# Lineage-specific evolution of LTR retrotransposons under natural selection drives genomic divergence in diploid *Oryza* species

**DOI:** 10.64898/2026.06.16.732638

**Authors:** Rong-jing Xu, Li-zhi Gao

## Abstract

LTR retrotransposons (LTR-RTs) are major drivers of plant genome evolution. However, the principles governing how natural selection shapes their lineage-specific dynamics across closely related species remain elusive. Here, we performed a comprehensive comparative analysis of LTR-RTs across 15 diploid *Oryza* species, representing all major genome types. We reconstructed the spatiotemporal distribution and evolutionary trajectories of LTR-RTs, revealing three distinct evolutionary patterns influenced by natural selection: lineage-specific expansion under purifying selection, balanced co-evolution under balancing selection, and lineage-specific retention under strong positive selection. We further demonstrated that species-specific LTR-RT families significantly contribute to highly diverged regions (HDRs) and non-aligned regions (NOTALs), drive segmental duplications, and influence genome size variation. Notably, the removal of LTR-RTs, particularly from medium-removal-rate families, is a key factor in genome size contraction. Our study provides a population-genetic perspective on LTR-RT evolution and highlights their critical role in shaping genomic divergence and adaptation in *Oryza*. Our findings provide a population-genetic framework for understanding how the co-evolutionary dynamics between TEs and their hosts shape genome architecture and adaptation in closely related species.

## 1. Introduction

Transposable elements (TEs) are ubiquitous repetitive sequences that constitute a major fraction of eukaryotic genomes and serve as a substantial source of genetic variation and evolutionary innovation [**1–4**]. Among TEs, long terminal repeat retrotransposons (LTR-RTs) represent the predominant component in plant genomes. For instance, LTR-RTs occupy approximately 74.81% of the maize genome, 27.8% of the rice genome, and 65.51% of the wheat genome [**5–7**]. Based on the order of reverse transcriptase (RT) and integrase (INT) domains within the *pol* gene, plant LTR-RTs are primarily classified into two superfamilies, Ty1/*Copia* and Ty3/*Gypsy*, each of which can be further divided into distinct lineages based on sequence and structural features [**8–10**]. Among closely related species, the species-specific amplification and contraction of distinct lineages drive genomic differentiation, influencing centromere structure, gene regulatory networks, and genome size [**11,12**]. Examples include CRM lineage amplification driving centromeric variation in rice [**13**], solo-LTRs acting as novel cis-regulatory elements [**14**], and bursts of specific LTR-RT lineages contributing to genome size variation in *Oryza* [**15**]. Comparative genomics of wild rice species such as *O. rufipogon* and *O. longistaminata* has further revealed that repetitive elements may have contributed to the genomic features [**16**]. Similarly, analyses of the upland wild rice *O. granulata* and the adaptive *O. rufipogon* genome have underscored the role of LTR-RTs in genome size dynamics [**17,18**].

Independent of host replication, LTR-RTs proliferate akin to asexual organisms, generating clonal lineages that diversify into families [**1,19,20**]. Their evolution depends not only on intrinsic activity but also on natural selection within the host genome [**17**], which shapes their distribution and evolutionary dynamics [**21–24**]. For instance, purifying selection influences LTR-RT distribution in *Brachypodium distachyon* [**25,26**], and selective pressures underlie their non-random distribution in *Arabidopsis thaliana* [**22**]. The genus *Oryza*—with its diverse genome types, high-quality sequences, significant variation in genome size, and well-resolved phylogeny—provides an ideal system for studying LTR-RT evolution [**15,27–37**]. Early surveys of LTR retrotransposons in *O. sativa* laid the groundwork for understanding their evolutionary history [**38**], while more recent studies on tea tree and rubber tree genomes have demonstrated how bursts of non-autonomous LTR-RTs and their lineage-specific evolution drive genome size expansion, offering parallels to processes in *Oryza* [**39–41**]. In rice, natural selection drives diverse “life-histories” among LTR-RT families, affecting copy number, density, genic distribution, and evolutionary strategy [**21,35,36,37,42**]. Previous studies have revealed diverse “life-histories” of LTR-RT families within species like rice [**35,42–44**]. Yet, a systematic, population-genetic analysis of lineage-specific dynamics across a whole genus—necessary to disentangle the roles of selection, drift, and lineage-specific innovation—is still lacking.

LTR-RTs are key drivers of structural variations (SVs), generating insertions, deletions, inversions, duplications, and presence-absence variations (PAVs) through mechanisms such as turnover, illegitimate recombination, and ectopic recombination, thereby promoting interspecific and intraspecific differentiation [**15,45–47**]. In *Oryza*, PAVs are among the most abundant SVs [**15,32**] and often correspond to highly diverged regions (HDRs) and non-aligned regions (NOTALs) [**48–50**]. Such regions are associated with species differentiation, adaptation, and domestication in various taxa [**51–54**], yet the specific role of LTR-RTs in their formation remains poorly understood. Beyond these SVs, segmental duplications (SDs) represent another important SV class, contributing to gene copy number variation and genome evolution [**6,55–62**]. While transposable elements like LINEs and *Alu* elements are established drivers of SDs in primates [**63–65**], their role in plants is less clear. Recent studies suggest LTR-RTs influence the formation, expansion, and evolution of SDs in plants [**9,66,67**], but the nature of this relationship requires further investigation.

Genome size is a fundamental genomic trait and an important outcome of evolution, yet its determinants remain incompletely resolved [**68–70**]. Beyond polyploidization, LTR-RT activity is a primary factor influencing genome size variation among species [**29,66,71–75**]. While recent bursts of LTR-RTs can drive genome expansion, the role of their removal in genome contraction is debated: some studies suggest LTR-RT removal significantly reduces genome size [**42,75**], whereas others argue it has limited impact [**76,77**]. Investigations into the genus *Eragrostis* have revealed that genome size variation and polyploidy prevalence are associated with global dispersal in arid areas, highlighting the ecological dimension of genome size evolution influenced by repetitive elements [**78**].

In this study, we performed a comprehensive analysis of LTR-RTs across 15 high-quality genomes representing the major diploid types in *Oryza*. Our work had two main objectives: (1) to elucidate how natural selection shapes the lineage-specific evolutionary dynamics of LTR-RTs across closely related species from a population genetics perspective, and (2) to evaluate the contribution of species-specific LTR-RT families to genomic divergence by examining their roles in HDRs/NOTALs formation, SD evolution, and genome size variation. This integrated approach provides new insights into the co-evolutionary interplay between TEs and their host genomes.

## 2. Results

### 2.1. Landscape and lineage-specific activity of LTR-RTs

The abundance and distribution of LTR-RTs within host genomes reflect their evolutionary dynamics and activity across species. To investigate the variation in LTR-RT content, we characterized transposable elements (TEs) from 15 diploid *Oryza* species using a comprehensive pipeline (**Fig. 1a-b; Additional file 2: Table S1**). A total of 6,219,917 TE fragments were identified, spanning 3,860,480,528 base pairs (bp). We observed considerable variation in TE content among species. For example, *O. brachyantha* possessed the lowest total TE length, with DNA transposons and LTR-RTs occupying 34.43 Mb and 43.67 Mb, respectively. In contrast, *O. australiensis* exhibited the highest total TE content (616.63 Mb), including 85.68 Mb of DNA transposons and 484.91 Mb of LTR-RTs. LTR-RTs were the most abundant TE type across all diploid *Oryza* species and showed the greatest degree of interspecific variation (**Additional file 2: Table S2**).

**Figure 1.**
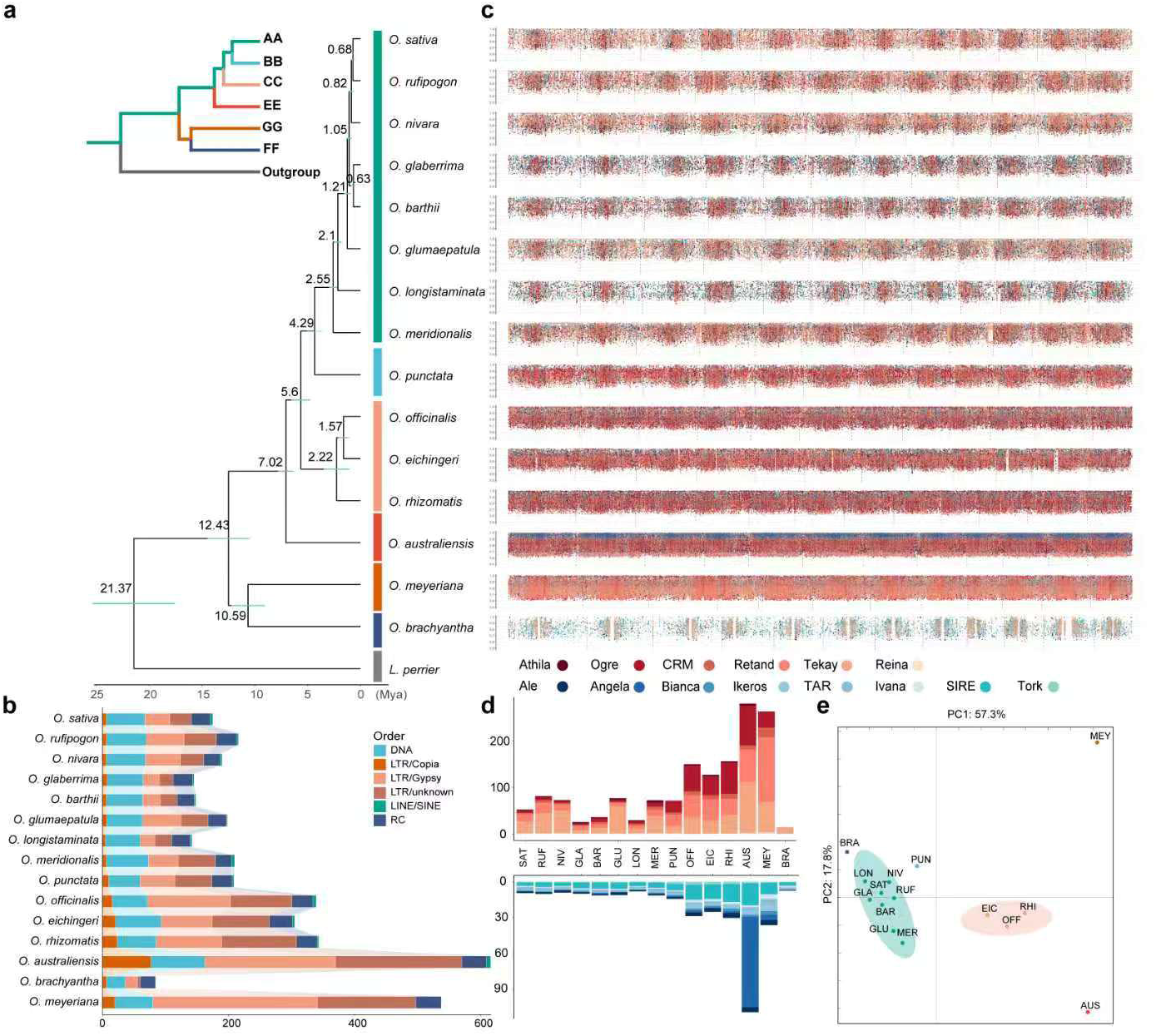
Distribution and composition of LTR-RTs across 15 *Oryza* diploid species. **(a)** Phylogeny of the 15 diploid *Oryza* species, with *L. perrieri* as the outgroup. The inset in the upper left depicts the phylogeny of different genome types in *Oryza*; the timeline is shown at the bottom. **(b)** Total content of transposable elements (TEs) from different orders in each species. The x-axis indicates TE length in base pairs (bp). “DNA” refers to Class I DNA transposons; “LTR/Copia” indicates LTR-RTs from the Ty1/*Copia* superfamily; “LTR/Gypsy” indicates LTR-RTs from the Ty3/*Gypsy* superfamily; “LINE/SINE” includes elements from LINE and SINE orders; “RC” represents rolling-circle (Class II) DNA transposons. **(c)** Genome-wide distribution landscape of LTR-RT sequences. Each point corresponds to one LTR-RT. The y-axis represents the Kimura 2-parameter (K2P) divergence transformed as (100 – K2P distance)/100. Grey dashed lines separate the 12 chromosomes; within each interval, the left and right sides correspond to the start and end of the chromosome, respectively. The color legend indicates the lineage origin of each LTR-RT and is shared with panel (d). **(d)** Cumulative length of LTR-RTs belonging to different lineages. The y-axis indicates length in bp. **(e)** Principal component analysis (PCA) based on the total length of LTR-RTs from different lineages across species.

Following integration, most TEs accumulate random mutations and lose activity. The age of each TE fragment was estimated using the Kimura 2-parameter (K2P) distance between the fragment and its corresponding consensus sequence. To assess how LTR-RT activity influences content variation, we constructed activity landscapes for different TE types (**Additional file 1: Fig. S1, S2**). These analyses revealed that both recent bursts and long-term activity of LTR-RTs contribute to content variation across species. In AA-type genomes, recent LTR-RT expansions were detected in *O. sativa*, *O. rufipogon*, *O. nivara*, *O. glumaepatula*, and *O. meridionalis*. In contrast, *O. glaberrima*, *O. barthii*, and *O. longistaminata* showed no evidence of recent bursts; instead, their LTR-RT activity followed a long-term stable expansion pattern (**Fig. S1, S2**). A similar long-term stable pattern was observed in the BB-type genome of *O. punctata*. Within CC-type genomes, recent expansion events occurred in *O. officinalis* and *O. rhizomatis*, while expansion in *O. eichingeri* was more moderate. In EE-type (*O. australiensis*) and GG-type (*O. meyeriana*) genomes, LTR-RT activity indicated accelerated accumulation and recent bursts, with the burst magnitude being substantially larger in *O. australiensis*. Conversely, in the FF-type genome of *O. brachyantha*, recent LTR-RT accumulation was limited, and long-term accumulation levels were lower than in other species. These findings suggest that LTR-RT activity and differentiation contribute to the divergence of genome types within *Oryza*.

To examine the distribution, accumulation, and activity of different LTR-RT lineages across species, we constructed spatiotemporal distribution landscapes for both LTR-RT fragments and intact elements (**Fig. 1c; Additional file 1: Fig. S3–S14; Additional file 2: Table S3**). The distribution of LTR-RTs among lineages and the spatiotemporal patterns of intact LTR-RTs varied across species. In AA-, BB-, CC-, EE-, and GG-type genomes, recent expansion of the Ty3/*Gypsy* superfamily exceeded that of Ty1/*Copia*. Within AA-type genomes, Tekay and Retand were the primary expanding lineages, both having undergone recent bursts. Distinct distribution patterns of Ty3/*Gypsy* and Ty1/*Copia* superfamilies were observed in *O. glaberrima*, *O. barthii*, and *O. longistaminata* compared to other AA-type species. In the BB-type genome, Ogre and Tekay were the most recently expanded lineages, with Tekay showing a pronounced expansion within the last 0–1 million years (Mya). The SIRE lineage (Ty1/*Copia*) also expanded in the BB genome around 0–0.8 Mya. In CC-type genomes, Ogre, Retand, and Tekay lineages showed continuous expansion and accumulation, with Ogre displaying the largest increase. SIRE also expanded within 0–2 Mya in these genomes. In EE-type genomes, Ogre and Tekay lineages exhibited the most extensive expansion, with Tekay undergoing a burst within 0–0.1 Mya. Additionally, the Angela lineage (Ty1/*Copia*) underwent a large-scale expansion in EE genomes within 0–2 Mya, predominantly accumulating in centromeric regions. In GG-type genomes, Retand was the primary accumulating lineage. In contrast, the FF-type genome showed the most moderate expansion and accumulation across all LTR-RTs, with a more dispersed distribution. In *O. brachyantha*, low levels of expansion and accumulation were observed for the SIRE, Ivana, and Tekay lineages.

We further analyzed the distribution of LTR-RTs from different lineages in centromeric regions and in the 2 kb regions upstream and downstream of genes. Quantification of LTR-RT length in centromeres revealed variation in their activity and accumulation across species (**Additional file 1: Fig. S15-S16**). In all diploid species, the total length of Ty3/*Gypsy* elements in centromeres exceeded that of Ty1/*Copia*. In AA-type genomes, CRM, Tekay, and Retand lineages were major components of the centromeres in *O. sativa*, *O. rufipogon*, *O. nivara*, *O. glaberrima*, and *O. barthii*, with CRM being the most abundant. In the centromeres of *O. glumaepatula*, *O. longistaminata*, and *O. meridionalis*, Tekay, Athila, CRM, and Ogre lineages showed higher abundance. Notably, in *O. glumaepatula*, Tekay, Athila, and Ogre lineages were more abundant than CRM, while in *O. meridionalis*, Tekay substantially outnumbered CRM. The composition of centromeric LTR-RTs in the BB-type genome was similar to AA-type genomes, though the Reina lineage was less abundant and Athila was absent. In CC-type genomes, centromeres contained more SIRE lineage elements compared to other *Oryza* species. The EE-type genome centromeres were distinct in containing a substantial number of Angela lineage elements. In the FF-type genome, overall centromeric LTR-RT content was lower than in other species, though Tekay was relatively abundant. In the GG-type genome, Tekay constituted the predominant centromeric component.

Analysis of LTR-RT distribution in the 2 kb flanking regions of genes revealed that Ty3/*Gypsy* fragments were more abundant than Ty1/*Copia* across all species (**Additional file 1: Fig. S17**). In *O. sativa*, *O. rufipogon*, and *O. nivara*, Retand and Tekay were the most abundant lineages near genes. In *O. glaberrima*, *O. barthii*, and *O. longistaminata*, Tekay, Retand, and Ogre showed higher abundance. In *O. glumaepatula*, only Tekay was predominant in these regions. *O. meridionalis* contained notably fewer LTR-RTs near genes compared to other AA-type genomes. In the BB-type genome, Ogre was the most abundant lineage near genes. Across CC-type genomes, Retand, Tekay, and Ogre were predominant; the distribution patterns in *O. officinalis* and *O. rhizomatis* were more similar to each other than to *O. eichingeri*. In the EE-type genome, Ogre, Tekay, and Retand remained primary, though Angela and SIRE were also more abundant compared to other species. In *O. brachyantha*, LTR-RTs near genes were generally scarce, primarily consisting of Tekay and SIRE, and their abundance increased with distance from gene boundaries.

Finally, principal component analysis (PCA) based on the content of LTR-RTs from different lineages across species indicated that the differentiation and activity of LTR-RTs drive variation in their content and distribution, significantly influencing the formation and differentiation of genomic types in *Oryza* (**Fig. 1d-e**).

### 2.2. Evolutionary patterns of LTR-RTs under natural selection

To reconstruct the dynamic evolutionary history of LTR-RTs in *Oryza* diploid species, we constructed phylogenetic trees for the Ty3/*Gypsy* and Ty1/*Copia* superfamilies based on reverse transcriptase (RT) sequences from 29,178 and 10,876 intact LTR-RTs, respectively (**Fig. 2a-b**). These analyses revealed distinct levels of differentiation and expansion for each lineage across different species. Within the Ty1/*Copia* superfamily, the Angela lineage displayed a unique pattern, primarily differentiating and undergoing continuous expansion in the genome of *O. australiensis*. The SIRE lineage predominantly differentiated and expanded in the three CC-type genome species. In contrast, the Ivana and Ikeros lineages expanded and evolved mainly in non-AA-type genomes. The Ale, Tork, and Bianca lineages maintained continuous differentiation and remained active across all diploid species. The TAR lineage expanded primarily in AA-type genomes. In the Ty3/*Gypsy* superfamily, the Athila lineage showed low levels of differentiation and expansion. The Reina and CRM lineages continued to expand and evolve across all species. The Tekay lineage exhibited higher levels of differentiation and expansion in AA-, CC-, and EE-type genomes compared to others, while the Retand and Ogre lineages displayed more complex evolutionary patterns (**Fig. 2a-b; Additional file 1: Fig. S18-S19**).

**Figure 2.**
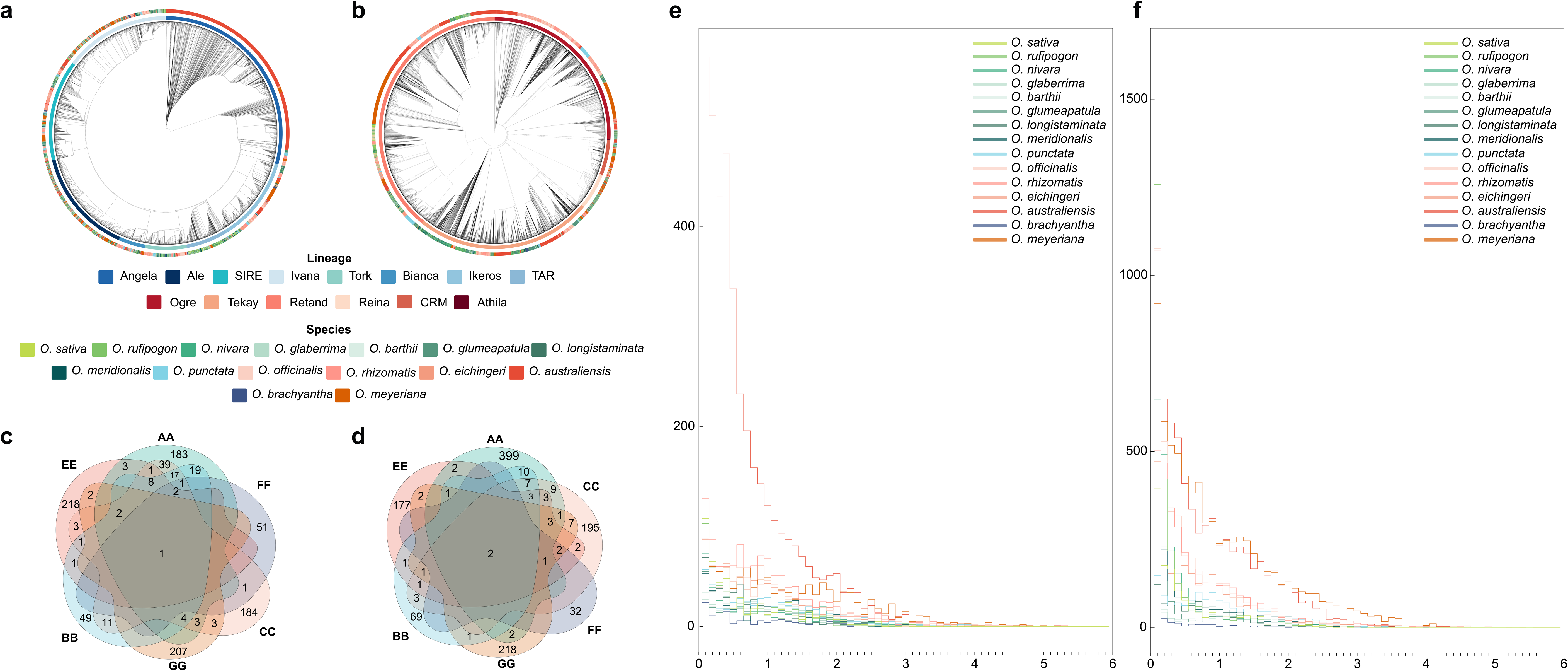
Dynamic evolutionary history of intact LTR-RTs in *Oryza* diploid species. **(a)** Phylogenetic tree of reverse transcriptase (RT) sequences from intact Ty1/*Copia*LTR-RTs. **(b)** Phylogenetic tree of RT sequences from intact Ty3/*Gypsy* LTR-RTs. **(c)** Venn diagram showing the sharing of intact Ty1/*Copia* LTR-RT families across different genomic types. **(d)** Venn diagram showing the sharing of intact Ty3/*Gypsy* LTR-RT families across different genomic types. **(e)** Temporal dynamics of intact Ty1/*Copia* LTR-RT accumulation. The y-axis shows the count of intact LTR-RTs, and the x-axis represents time in millions of years ago (Mya). Each unit corresponds to 1 Mya. **(f)** Temporal dynamics of intact Ty3/*Gypsy* LTR-RT accumulation. The y-axis shows the count of intact LTR-RTs, and the x-axis represents time in millions of years ago (Mya). Each unit corresponds to 1 Mya.

To further investigate LTR-RT differentiation across species, we clustered intact LTR-RTs into families based on sequence similarity in their 5’ LTR regions, yielding 1,015 Ty1/*Copia* and 1,157 Ty3/*Gypsy* families. The distribution of these families indicated that very few are shared across different genome types or species (**Fig. 2c-d; Additional file 1: Fig. S20–S22**). Beyond differentiation levels, variation in LTR-RT activity among species was also evident. By calculating the insertion times of intact LTR-RTs, we identified four primary activity patterns (**Fig. 2e-f**). The first pattern, characterized by recent bursts of LTR-RTs, was observed in *O. sativa*, *O. rufipogon*, *O. nivara*, *O. glumaepatula*, *O. meridionalis*, and all three CC-type genome species. In these genomes, Ty3/*Gypsy* elements began expanding 0.1–0.7 Mya, with activity peaking within the last 0.1 Mya. The variation in activity was greater in these species than in others, and the content of intact Ty3/*Gypsy* LTR-RTs was particularly high in *O. glumaepatula* and *O. rufipogon*. The second pattern, featuring a gradual but slower increase in LTR-RT accumulation, included *O. barthii*, *O. glaberrima*, *O. longistaminata*, and *O. punctata*. This pattern was also followed by Ty1/*Copia* elements in all species except *O. australiensis*. The third pattern, marked by long-term high activity of Ty3/*Gypsy* LTR-RTs, was seen in *O. australiensis* and *O. meyeriana*; in *O. australiensis*, Ty1/*Copia* activity also followed this pattern. The fourth pattern, displaying long-term low LTR-RT activity, was specific to *O. brachyantha* (**Fig. 2e-f**).

To elucidate how natural selection shapes these complex evolutionary dynamics across lineages and species, we constructed phylogenetic trees for each lineage based on 5’ LTR sequences. Members clustered on a single branch were defined as subfamilies. We then characterized the length, domain content, and insertion time of major LTR-RT families within each lineage. Based on these results, we classified the evolutionary histories of LTR-RT lineages into three overarching patterns across the studied species.

The first and primary pattern in *Oryza* is that differential selective pressures drive divergent evolutionary histories for the same lineage across species. Positive or balancing selection can promote small-scale expansion and differentiation, leading to higher diversity within a single species but lower differentiation between species. In contrast, negative (purifying) selection can drive long-term large-scale expansion, recent bursts, or contraction of LTR-RT lineages, resulting in higher differentiation between species (**Additional file 1: Fig. S23–S40; Additional file 3: Table S4; Additional file 4: Table S5**). Lineages following this pattern include Ogre, Tekay, Retand, Ikeros, Ivana, TAR, Tork, Athila, and Bianca.

Specifically, the Ogre lineage showed high differentiation (Fst ≥ 0.15) between species of different genome types, and moderate to low differentiation (0 < Fst < 0.15) within groups. This lineage expanded, differentiated, and evolved predominantly in non-AA-type genomes, where purifying selection drove the formation of new families and recent proliferation bursts. In AA-type genomes, under balancing selection, differentiation and expansion levels were lower, as seen in the Ogre4 and Ogre7 families (**Additional file 1: Fig. S23-S24**). The Tekay lineage differed, being most abundant in AA-type genomes. Purifying selection drove high differentiation between species of different genome types, fostering the formation, expansion, and differentiation of new families in species including *O. sativa*, *O. rufipogon*, *O. nivara*, *O. longistaminata*, *O. meridionalis*, *O. officinalis*, *O. eichingeri*, *O. rhizomatis*, and *O. australiensis* (**Additional file 1: Fig. S25-S26**). However, differentiation between *O. australiensis* and *O. glumaepatula* was moderate (Fst = 0.146), likely due to the expansion of the Tekay1 family after its divergence into two subfamilies in these species. Balancing selection influenced Tekay elements in *O. meyeriana*, *O. glaberrima*, *O. barthii*, and *O. glumaepatula*, leading to similar activity patterns (**Additional file 1: Fig. S25-S26**). The Retand lineage (Ty3/*Gypsy*) contained more intact elements in non-AA-type genomes. There, purifying selection drove long-term proliferation and recent bursts, enhancing differentiation between species. In *O. meyeriana*, however, balancing selection led multiple autonomous and non-autonomous Retand families to maintain similar long-term expansion and differentiation patterns (**Additional file 1: Fig. S27-S28**). In AA-type genomes, the Retand lineage sustained similar expansion patterns across species with moderate to low inter-species differentiation (**Additional file 3: Table S4; Additional file 4: Table S5**). The TAR lineage was influenced by purifying selection in *O. barthii*, *O. glaberrima*, *O. meyeriana*, and *O. rhizomatis*, where two families-maintained expansion and differentiation. In other species, positive selection resulted in multiple families exhibiting a mixed pattern of long-term proliferation and recent bursts (**Additional file 1: Fig. S31-S32**). Similarly, the Tork lineage experienced expansion and differentiation under varying selective pressures (**Additional file 1: Fig. S33-S34**). In the Athila and Bianca lineages, more non-autonomous LTR-RTs underwent continuous expansion and differentiation under balancing selection. Within the Bianca lineage, differentiated families expanded in EE- and CC-type genomes under purifying selection (**Additional file 1: Fig. S35–S38**). The Ivana lineage also showed distinct histories under different selective pressures; for example, the Ivana4, Ivana6, and Ivana14 families exhibited similar expansion and differentiation under balancing selection in *O. meyeriana* (**Additional file 1: Fig. S39-S40**).

The second pattern is characterized by all LTR-RTs within a lineage being influenced by either balancing or purifying selection across all species. In the Reina and Ale lineages, LTR-RTs were influenced by balancing selection, with several important families driven to diverge and expand on a small scale across species (**Additional file 1: Fig. S41–S44**). Lineages influenced by purifying selection contained more members than those under positive selection. Notably, these purifying selection lineages also contained more non-autonomous LTR-RTs, which themselves were under purifying selection, leading to specific differentiation and expansion of certain subfamilies or phylogenetic branches (**Additional file 1: Fig. S45–S48**). For instance, differentiated families from the SIRE and CRM lineages maintained rapid differentiation and expansion, or formed species-specific subfamilies undergoing continuous evolution, under purifying selection (**Additional file 1: Fig. S45–S48**).

The third and final pattern is defined by the majority (≥ 90%) of a lineage’s LTR-RTs being retained in only one genome type. In *Oryza*, this pattern is unique to the Angela lineage. We identified 3,167 intact Angela LTR-RTs in *O. australiensis*, where the lineage is under purifying selection. Based on all-to-all BLAST of 5’ LTRs, 3,313 of these intact elements were clustered into the Angela1 family, which maintains long-term expansion and differentiation under purifying selection in *O. australiensis* (**Additional file 1: Fig. S49-S50**).

Collectively, these three evolutionary patterns demonstrate that natural selection acts differentially on LTR-RT lineages across the *Oryza* phylogeny. We next investigated how these lineage-specific evolutionary dynamics, and particularly the proliferation of species-specific families, translate into tangible impacts on genome architecture and variation.

### 2.3. Species-specific LTR-RTs are major components and potential drivers of genomic divergence in HDRs and NOTALs

LTR-RT activity is a key driver of structural variation (SV) in *Oryza*. To assess its impact, we identified a total of 2,063,975 SVs across the studied species using the *O. sativa*genome as a reference. These SVs comprised 1,923,111 insertions and deletions, 23,613 duplications, 1,817 inversions, 11,493 translocations, 55,822 highly diverged regions (HDRs), and 48,119 non-aligned regions (NOTALs) (**Fig. 3a-b; Additional file 2: Table S6**).

**Figure 3.**
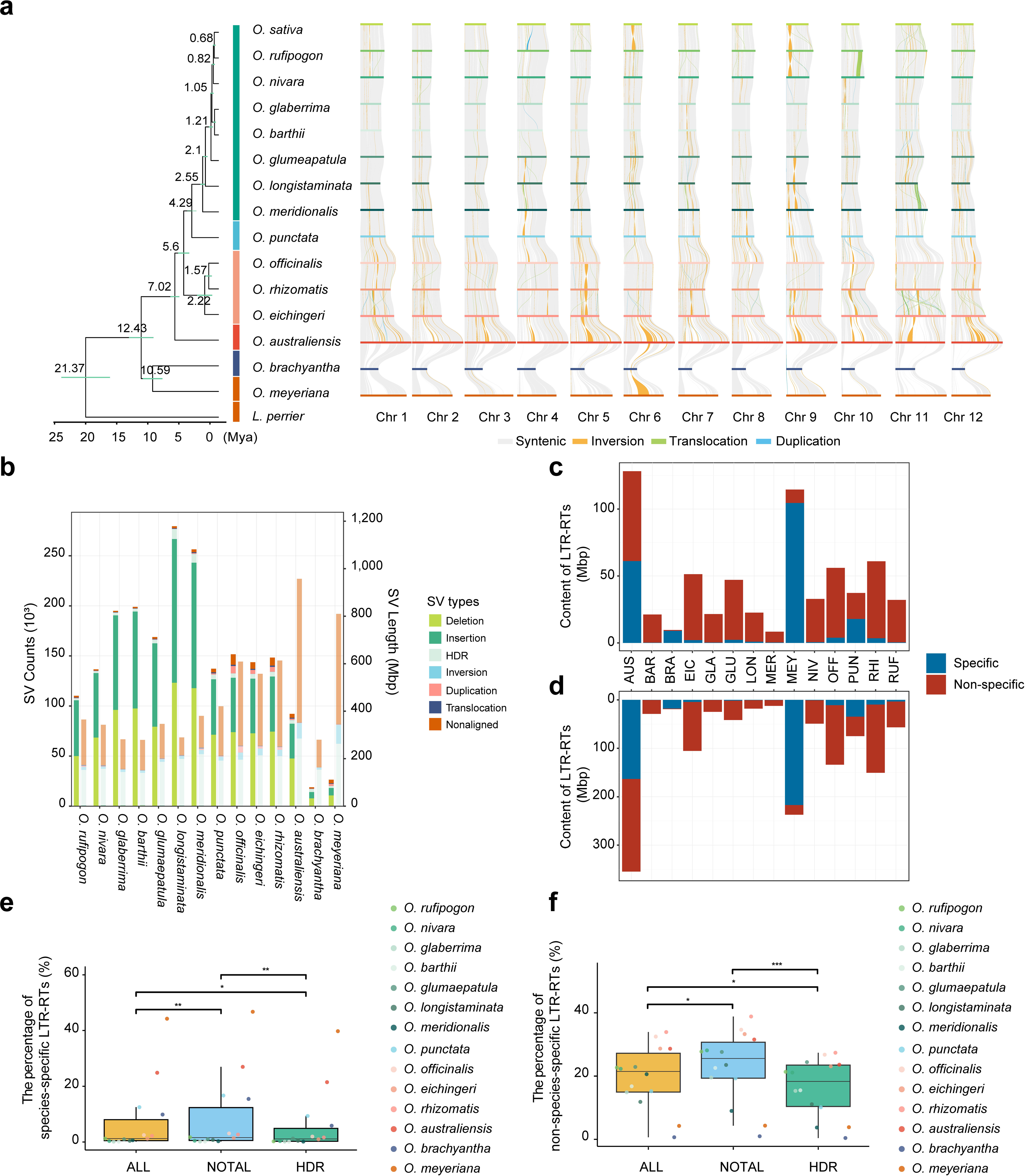
Contribution of LTR-RTs to highly diverged (HDR) and non-aligned (NOTAL) regions in *Oryza* diploid species. **(a)** Chromosomal synteny among 14 *Oryza* diploid species. **(b)** Counts (left bars) and total length (right bars) of structural variations (SVs) identified in each species. **(c)** Total length of LTR-RTs located within HDRs. **(d)** Total length of LTR-RTs located within NOTALs. **(e)** Percentage of species-specific LTR-RT families across different genomic contexts: whole genome (ALL), non-aligned regions (NOTAL), and highly diverged regions (HDR). **(f)** Percentage of non-species-specific LTR-RT families across the same genomic contexts. In panels (**e**) and (**f**), asterisks indicate statistical significance: \**p*<0.05, \*\**p*<0.01, \*\*\**p*<0.005.

To evaluate the contribution of LTR-RT divergence to the formation of HDRs and NOTALs, we quantified both species-specific and shared (non-species-specific) LTR-RT families within these regions. Substantial amounts of both LTR-RT types were detected. In HDRs, species-specific LTR-RTs spanned 206,396,816 bp (range: 150,829 to 104,510,766 bp per species), while shared LTR-RTs spanned 437,571,921 bp (range: 626,826 to 67,167,662 bp per species) (**Figure 3c; Additional file 2: Table S7**). In NOTALs, species-specific LTR-RTs accounted for 464,556,914 bp (range: 200,271 to 216,975,177 bp per species), and shared LTR-RTs accounted for 841,157,940 bp (range: 1,117,712 to 191,122,636 bp per species) (**Fig. 3d; Additional file 2: Table S7**).

We then compared the relative abundance (percentage) of these LTR-RT categories within HDRs and NOTALs to their genome-wide levels (**Fig. 3e-f**). In NOTALs, the relative content of species-specific LTR-RTs was significantly higher than in both the whole genome and in HDRs. Similarly, the relative content of shared LTR-RTs in NOTALs was significantly higher than in the whole genome and markedly higher than in HDRs. Within HDRs, the relative content of species-specific LTR-RTs was significantly elevated compared to the whole genome, whereas the relative content of shared LTR-RTs was significantly reduced (**Fig. 3e-f**). These results indicate a distributional bias of species-specific LTR-RTs toward both NOTALs and HDRs. Shared LTR-RTs also showed a preferential distribution within NOTALs.

To further examine the relationship between LTR-RTs and these highly diverged genomic regions, we performed correlation analyses between the content of species-specific/shared LTR-RTs and the features of HDRs and NOTALs (**Additional file 1: Fig. S51-52**). The total length of HDRs showed a significant negative correlation with the percentage of shared LTR-RTs within HDRs (p = 0.021). Conversely, the proportion of the genome occupied by HDRs was strongly positively correlated with the percentage of species-specific LTR-RT families, both genome-wide and within HDRs themselves (**Additional file 1: Fig. S51**). While the total length of NOTALs did not show a significant correlation with either LTR-RT type, the genomic proportion of NOTALs was significantly positively correlated with the percentage of species-specific LTR-RT families, measured both genome-wide and within NOTALs (**Additional file 1: Fig. S52**).

Together, these findings demonstrate that the formation and expansion of HDRs and NOTALs are influenced by the differentiation and proliferation of species-specific LTR-RTs. However, the activity of species-specific LTR-RTs alone does not fully account for the emergence and variation of these divergent genomic regions.

### 2.4. Reciprocal interactions between LTR-RTs and segmental duplications

Segmental duplications (SDs) represent an important category of structural variations and play a significant role in the adaptive evolution of *Oryza* species. To investigate the reciprocal impacts between LTR-RTs and SDs, we identified SDs across the studied genomes. In total, we detected 16,845 intra-chromosomal SDs (spanning 148,155,634 bp) and 61,773 inter-chromosomal SDs (spanning 178,941,900 bp). Among these, 8,805 intra-chromosomal and 33,725 inter-chromosomal SDs were found to contain LTR-RTs (**Fig. 4a; Additional file 2: Table S8**). The number of SDs containing LTR-RTs varied among species, with *O. officinalis*, *O. rhizomatis*, and *O. meyeriana* each containing more than 1,000 intra-chromosomal SDs that include LTR-RTs. These findings indicate that LTR-RT activity influences the expansion and divergence of SDs across different species.

**Figure 4.**
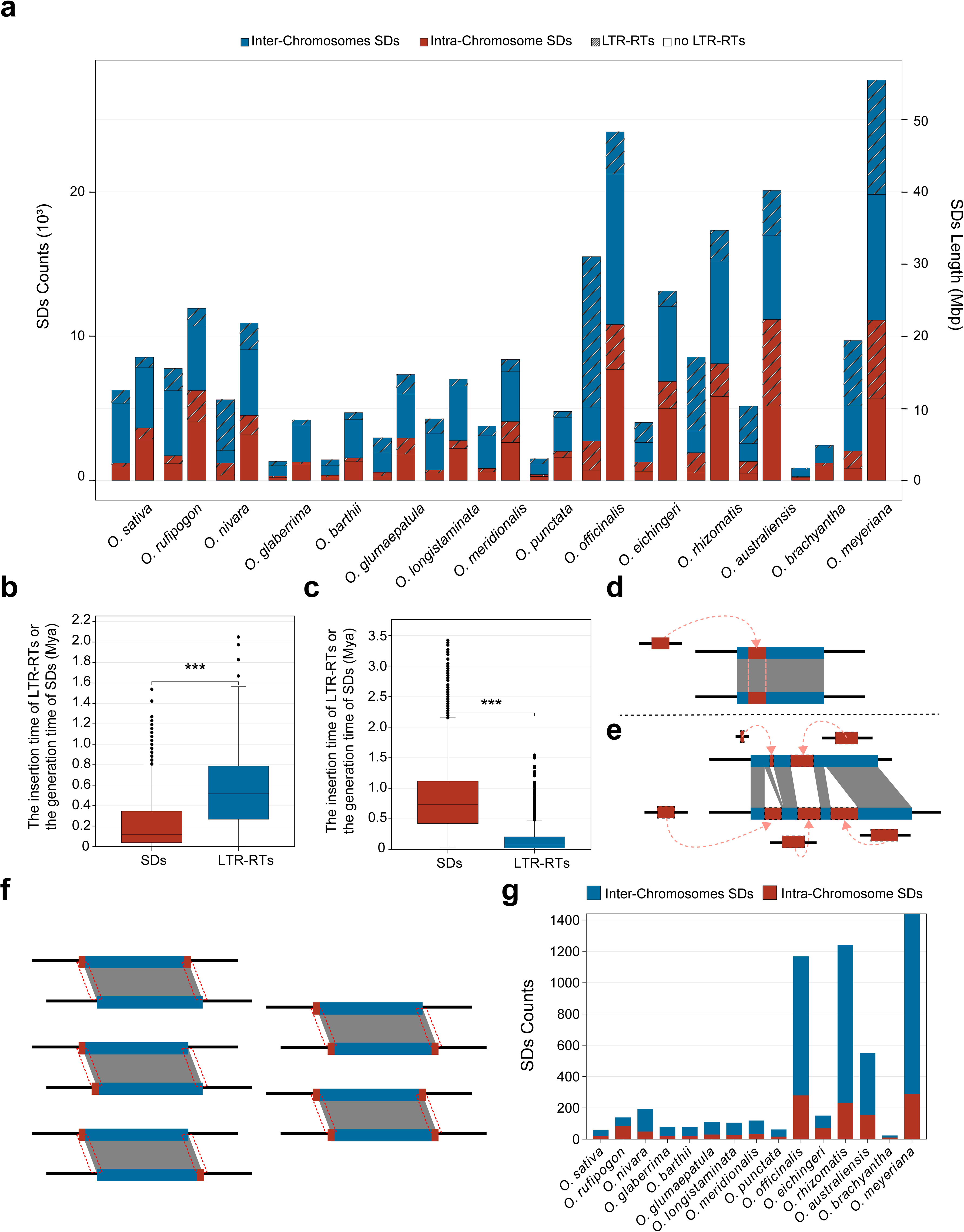
Influence of LTR-RTs on the expansion and formation of segmental duplications in *Oryza* diploid species. **(a)** Number (left bar) and total length (right bar) of SDs identified in the 15 *Oryza* diploid species. **(b)** Comparison between the insertion time of 731 intact LTR-RTs and the formation time of the SDs that contain them, showing that LTR-RT insertion significantly preceded SD generation. **(c)** Comparison between the insertion time of 5,373 intact LTR-RTs and the formation time of the SDs that contain them, showing that LTR-RT insertion occurred significantly after SD generation. Asterisks denote statistical significance: *p<0.05, **p<0.01, ***p<0.005. Two models depicting reciprocal interactions between LTR-RTs and SDs: **(d)** LTR-RT expansion accompanies SD occurrence; **(d)** LTR-RTs insert into SD regions and promote their enlargement. Red rectangles represent LTR-RTs, blue rectangles represent SD regions, and black lines indicate chromatin. **(e)** Schematic of the criterion used to identify SDs potentially driven by LTR-RTs. Each SD has four breakpoint junctions (5lbp flanks); SDs with two or more junctions overlapping LTR-RTs were classified as LTR-RT-potentiated. **(g)** Proportion of SDs classified as potentially driven by LTR-RTs in each *Oryza* diploid species.

Next, we compared the estimated formation time of SDs containing intact LTR-RTs with the insertion time of those intact elements (**Fig. 4b-c; Additional file 2: Table S9**). We identified 730 intact LTR-RTs that inserted *before* the associated SDs were formed (**Fig. 4b**), and 5,374 intact LTR-RTs that inserted *after* the formation of the SDs in which they reside (**Fig. 4c**). These results reveal two distinct patterns of interaction between LTR-RTs and SDs (**Fig. 4d**): one in which the creation of new genomic regions via SDs provides novel insertion sites for LTR-RTs, and another in which SDs occur within genomic regions that already contain LTR-RTs. The former pattern suggests that LTR-RTs can contribute to the expansion of SD regions, while the latter highlights how SDs can amplify the copy number of pre-existing LTR-RTs.

To further assess the influence of LTR-RTs on SD formation, we identified SDs that were potentially driven by LTR-RTs based on the criterion that two or more breakpoint junctions of an SD overlap with LTR-RT sequences (**Fig. 4e**). A total of 5,518 such SDs were characterized. In most *Oryza* species examined, these LTR-RT-associated SDs constituted between 1% and 10% of the total SD content. However, in *O. rhizomatis*, *O. australiensis*, and *O. meyeriana*, this proportion was notably higher, reaching 14.5%, 10.67%, and 14.85%, respectively (**Fig. 4f; Additional file 2: Table S10**). Furthermore, compared to intra-chromosomal SDs, a greater number of inter-chromosomal SDs were identified as potentially driven by LTR-RTs. Collectively, these results demonstrate that LTR-RTs can influence the formation of SDs, potentially through mechanisms such as providing homologous sequences for non-allelic recombination with a particularly pronounced effect on the generation of inter-chromosomal duplications.

### 2.5. LTR-RT removal contributes to genome size variation

Variation in genome size represents one of the most significant impacts of LTR-RT activity on host genomes. To assess the contribution of LTR-RTs to genome size in *Oryza*, we characterized the population dynamics of LTR-RTs alongside the genome sizes of the studied species (**Fig. 5a; Additional file 5: Table S11**). The results indicate that the dynamic trend of genome size across species closely parallels the trend in the birth counts of LTR-RTs. In contrast, other metrics—including the ratio of intact LTR-RTs to birth counts, the ratio of solo LTRs to intact LTR-RTs, and the ratio of combined solo and truncated LTRs to intact LTR-RTs—did not show a clear similar or opposite relationship with genome size trends.

**Figure 5.**
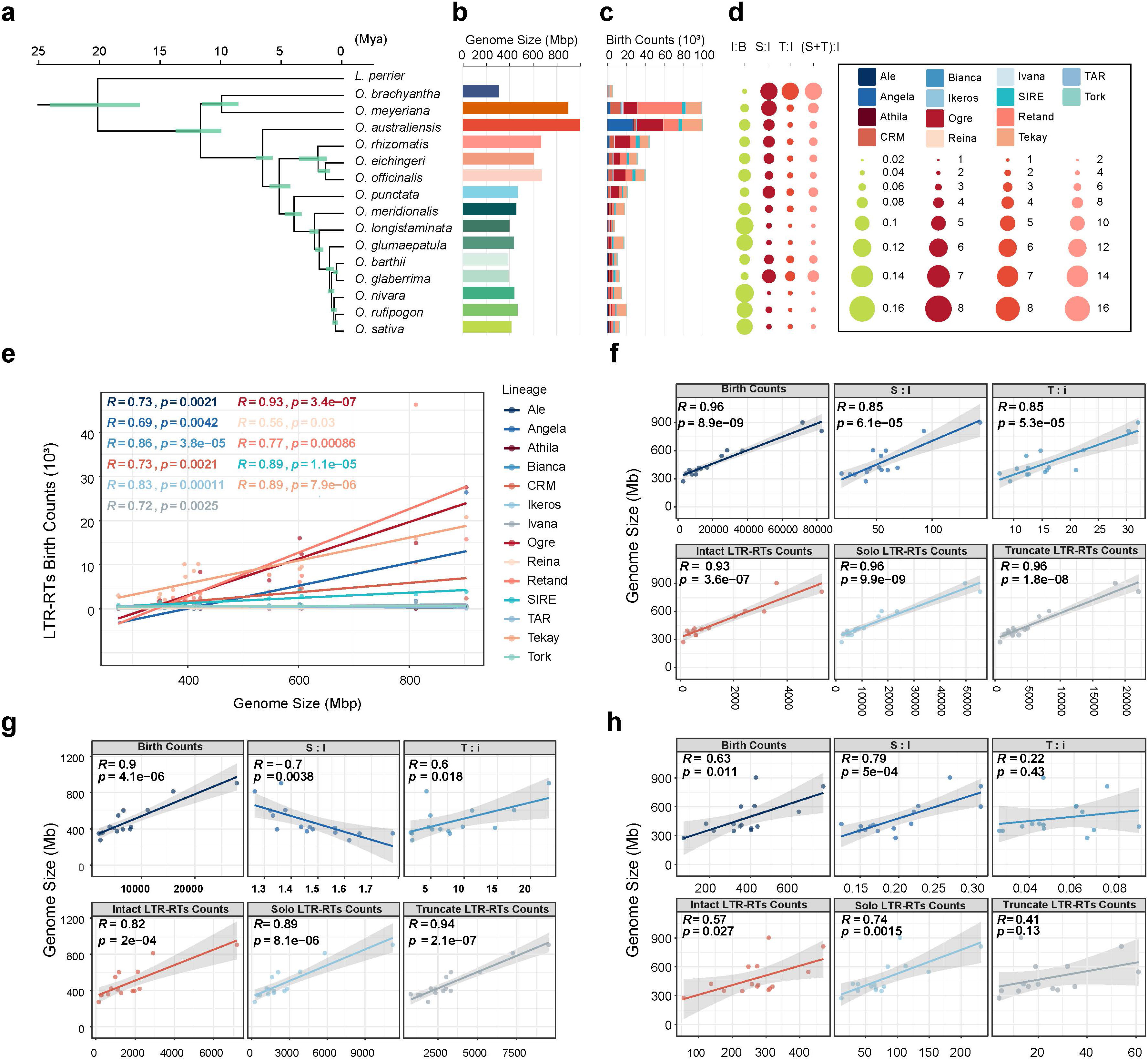
Contribution of LTR-RT removal to genome size variation in *Oryza* diploid species. **(a)** Population dynamics of LTR-RTs and genome size variation. **(b)** Genome size variation across species. **(c)** Birth counts of LTR-RTs. **(d)** Key LTR-RT metrics: I:B, ratio of intact LTR-RTs to birth counts; S:I, ratio of solo LTRs to intact LTR-RTs; T:I, ratio of truncated LTRs to intact LTR-RTs; (S+T):I, ratio of solo and truncated LTRs to intact LTR-RTs. **(e)** Pearson correlation between the birth counts of different LTR-RT lineages and genome size. **(f)–(h)** Pearson correlation analyses for LTR-RTs grouped by removal rate: high (f), medium (g), and low (h). For each group, correlations are shown between genome size and: (top) total counts of all LTR-RT types; (middle) S:I ratio; (bottom) T:I ratio.

To further examine the effects of LTR-RT proliferation and removal on genome size, we performed Pearson correlation analyses between the birth counts of different lineages and genome size, as well as between the solo-LTR-to-intact-LTR-RT ratio (a proxy for deletion rate) and genome size (**Fig. 5b; Additional file 1: Fig. S53**). These analyses revealed that the birth counts of different lineages have varying degrees of influence on genome size. Lineages such as Ogre, Retand, and CRM showed a significant positive correlation with genome size. However, the overall deletion rate of LTR-RTs was not significantly correlated with genome size (**Fig. 5b; Additional file 1: Fig. S53**).

To investigate the impact of LTR-RT removal more specifically, we classified LTR-RT families into high, medium, and low removal-rate categories based on the ratio of solo LTRs to intact LTR-RTs (**Additional file 1: Fig. S54–56; Additional file 5: Table S11**). The results show that high removal-rate families contain more members and constitute a greater number of families overall. Furthermore, the proportion of species-specific LTR-RT families classified as high removal-rate was greater than that of species-specific families in the medium and low removal-rate categories. This suggests that high removal-rate families experience more frequent activity cycles.

We then conducted Pearson correlation analyses between the abundance of LTR-RTs within each removal-rate category and genome size, and between the removal rate itself and genome size (**Fig. 5c–e**). For high removal-rate LTR-RTs, all measured factors (counts and removal rate) showed a significant positive correlation with genome size (**Fig. 5c**). In contrast, for medium removal-rate LTR-RTs, the removal rate was significantly negatively correlated with genome size (**Fig. 5d**). For low removal-rate LTR-RTs, both the birth counts and the total counts of all LTR-RT types within this category were significantly positively correlated with genome size (**Fig. 5e**).

Collectively, these results demonstrate that LTR-RT removal is a significant, lineage-dependent contributor to genome size variation within *Oryza*. Notably, the removal of LTR-RTs from medium removal-rate families appears to have the most significant impact on genome size contraction.

## 3. Discussion

Our systematic analysis of LTR retrotransposons across 15 diploid *Oryza* species uncovers three primary evolutionary patterns shaped by natural selection: lineage-specific expansion under purifying selection, balanced co-evolution under balancing selection, and lineage-specific retention under strong positive selection. These patterns mechanistically link species-specific LTR-RT dynamics to key aspects of genomic divergence: their significant accumulation in highly diverged (HDRs) and non-aligned regions (NOTALs), their reciprocal interactions with segmental duplications (SDs), and their collective impact on genome size variation—where removal rates, especially from medium-removal-rate families, emerge as a critical factor for genome contraction. These findings establish LTR-RTs as central drivers of genomic divergence and adaptation in *Oryza*, offering a novel population-genetic perspective on TE-host genome co-evolution.

### 3.1. Lineage-specific evolution of LTR-RTs: A population-genetic perspective

Transposable elements (TEs) are fundamental components of eukaryotic genomes, playing crucial roles in shaping regulatory networks, epigenetic landscapes, genome topology, and providing a rich source of genetic variation [**14,79–83**]. In plants, long terminal repeat retrotransposons (LTR-RTs) constitute a major genomic fraction and significantly influence chromatin state, three-dimensional architecture, gene regulation, and centromere structure [**13,81,83**]. Extensive research across various plant genera, including *Oryza*, *Triticum*, and *Zea*, has explored these dynamics [**13,19,74,79,84**]. In this study, we identified 2.47 Gb of LTR-RT sequences, including 50,625 intact elements, across 15 diploid *Oryza* species representing all major genomic types. We constructed comprehensive distribution and spatiotemporal landscapes for these LTR-RTs. Our analyses revealed lineage-specific activity variations among species and their differential impacts on genomic regions, notably highlighting the influence of the Tekay lineage in AA-type genomes, the Angela lineage in *O. australiensis*, and the Ogre and Retand lineages in non-AA genomes. Consistent with prior work, our findings confirm variations in LTR-RT abundance across whole genomes, centromeres, and gene-flanking regions among diploid *Oryza* species [**15,32**]. Moreover, our data provide new evidence that the activity dynamics of different LTR-RT lineages vary significantly across distinct genomic types within *Oryza*. This is consistent with findings in other plant systems, such as tea tree and rubber tree, where recent bursts of specific LTR-RT lineages have been shown to drive genome size evolution and genome innovation [**39–41**]. Furthermore, the rapid diversification of AA-genome *Oryza* species has been linked to recent LTR-RT evolution, underscoring the tempo of TE-driven genome change [**85**].

The influence of natural selection on the evolutionary trajectories of LTR-RTs across species is complex and not fully understood [**20,21,86–90**]. Several conceptual frameworks have been proposed. The parasitic DNA hypothesis posits that LTR-RT variation is driven by diverse selective pressures acting on host genomes [**91,92**]. The transposition-selection balance model suggests TEs reach an equilibrium transposition rate under stabilizing selection [**93–96**]. In contrast, the transposition-burst model (or Red Queen hypothesis) proposes that novel TEs from horizontal transfer or recent formation can trigger replication bursts, leading to long-term antagonistic co-evolutionary cycles between hosts and TEs [**88,90,93,97–99**]. By reconstructing the evolutionary histories of LTR-RT lineages across 15 diploid species from six genomic types, our study offers a new population-genetic perspective on how natural selection operates at the scale of the genomic “community ecology” [**20**] (**Additional file 3: Table S4; Additional file 4: Table S5**). Aligning with previous findings in rice [**21**], our results underscore the significant roles of both negative (purifying) and positive selection in LTR-RT evolution. We further demonstrate that purifying selection is associated with lineages undergoing long-term expansion or recent bursts within a genome, whereas positive/balancing selection drives moderate, coordinated expansion across multiple LTR-RT families in several species. We identified three overarching patterns of LTR-RT evolution under natural selection in *Oryza*, providing empirical support consistent with the transposition-burst model.

The three evolutionary patterns we identified—selective expansion, balanced co-evolution, and lineage-specific retention—may represent generalizable modes of LTR-RT evolution in plant genomes subject to different genomic and selective environments. Testing this framework in other plant lineages with diverse genome sizes and life histories will reveal its broader applicability.

### 3.2. LTR-RTs as engines of structural genomic divergence

LTR-RTs are a major driver of structural variation (SV), contributing significantly to genomic differentiation among diploid *Oryza* species [**15,32,45,46**]. The availability of high-quality genomes has spurred recent investigations into LTR-RT-mediated SV formation in this genus [**6,13,15,27,30,31,45,46**]. Our study identified a comprehensive set of SVs (Supplementary Table S6) and, consistent with prior work, confirmed the abundance of highly diverged regions (HDRs) and non-aligned regions (NOTALs)—often categorized as presence-absence variations (PAVs) [**15,32,49**]—among these species. We show that the content of species-specific LTR-RT families significantly influences these complex divergent regions. Furthermore, our findings indicate that activities of various LTR-RT lineages drive the formation of intricate SVs and contribute to differentiation between genomic types. However, LTR-RT activity alone does not fully account for the genesis of all HDRs and NOTALs, suggesting the involvement of additional evolutionary mechanisms.

Segmental duplications (SDs) are a key mechanism for increasing gene copy number, providing vital genetic resources for evolution [**57,100,101**]. While LINEs and *Alu* elements are established drivers of SDs in primate genomes [**64,65**], their role in plants is less clear. Emerging evidence suggests LTR-RTs can influence SD regions in plants [**66,67,102**]. Our analysis of the timing of SD formation relative to the insertion of intact LTR-RTs within them (**Additional file 2: Table S9**) revealed a reciprocal relationship (**Fig. 4d**): LTR-RTs can insert into regions created by SDs, and SDs can occur within pre-existing LTR-RT clusters, occasionally amplifying them. This aligns with reports of LTR-RT integration into SDs [**67**]. Additionally, we found that LTR-RTs are particularly associated with a greater number of inter-chromosomal SDs (**Fig. 4f**). These insights offer new evidence for understanding TE-SD interactions in plant genomes. Such reciprocal interactions echo findings in the tea tree genome, where bursts of non-autonomous LTR-RTs were associated with genome size evolution, potentially mediated through duplication events [**39,40**].

### 3.3. The dual role of LTR-RT dynamics in genome size evolution

Genome size variation in eukaryotes is largely attributed to LTR-RT dynamics, aside from polyploidization [**71,75,86**]. Within *Oryza*, LTR-RT activity is a primary determinant of genome size diversity [**15,28,29,33,35,37,103**]. Our study confirms that recent bursts of LTR-RTs significantly contribute to genome size variation. While LTR-RT removal via mechanisms like solo-LTR formation can lead to DNA loss [**35,42,75,103**], we found no genome-wide negative correlation between the solo-LTR-to-intact-LTR-RT ratio and genome size (**Additional file 1: Fig. S51**). Instead, our novel classification of LTR-RT families based on removal rate revealed a nuanced relationship: the removal of elements from *medium* removal-rate families shows the most significant negative correlation with genome size, highlighting their particular importance in genome contraction. This finding provides a refined perspective on how LTR-RT deletion contributes to genome size evolution. This nuanced view complements studies in other genera, such as *Eragrostis*, where genome size variation and polyploidy are linked to ecological dispersal, suggesting that both expansion and contraction mechanisms are subject to evolutionary pressures tied to environmental adaptation [**78**]. Similarly, early surveys of LTR-RTs in cultivated rice highlighted their role in recent genome size change [**38**].

### 3.4. Conclusions and future perspectives

In conclusion, our study provides a comprehensive, population-genetic dissection of LTR-RT evolution and its genomic consequences across the diploid *Oryza* phylogeny. We demonstrate that natural selection operates in distinct modes—purifying and balancing selection—to shape lineage-specific expansion, co-evolution, and retention, forming the foundation for their diverse impacts. These differential evolutionary trajectories directly contribute to key facets of genomic divergence: the enrichment of species-specific LTR-RTs in HDRs and NOTALs underscores their role in creating inaccessible genomic variation; their reciprocal relationship with SDs reveals a dynamic mechanism for generating and amplifying structural complexity; and the nuanced link between removal rates and genome size highlights deletion as an active, lineage-dependent process in genome contraction. This integrated view firmly establishes LTR-RTs not merely as repetitive sequence components but as central, dynamic agents in the structural and adaptive evolution of *Oryza* genomes.

Looking forward, this work opens several promising avenues for research. First, the molecular mechanisms governing the distinct deletion rates among LTR-RT families, particularly the potent effect of medium-removal-rate families, remain unknown. Future studies combining epigenomic profiling (e.g., histone marks, DNA methylation) with comparative genomics could identify the host pathways that differentially target LTR-RT families for removal. Second, the functional consequences of LTR-RT-potentiated SDs deserve exploration. Do these SDs disproportionately generate novel gene copies or alter regulatory landscapes, contributing to phenotypic innovation? Third, while we focused on diploids, applying this framework to allopolyploid *Oryza* species could elucidate how genome merger and duplication reshape the evolutionary dynamics and genomic impact of LTR-RTs.

Finally, the population-level patterns of the identified species-specific LTR-RT families within wild *Oryza* populations could be explored to test their association with local adaptation, potentially linking specific TE-driven structural variants to ecological traits. Addressing these questions will further illuminate the intricate co-evolutionary dialogue between TEs and their plant hosts. Ultimately, understanding the rules governing TE evolution, as decoded here in *Oryza*, is fundamental to explaining the origins of genomic diversity, the constraints on genome architecture, and the raw material available for adaptation in flowering plants.

## 4. Materials and Methods

### 4.1. Genome sequences of fifteen *Oryza* diploid species

The reference genome of *O. sativa* ssp. *japonica* cv. Nipponbare (Release 7; abbreviated SAT) was obtained from the Rice Genome Annotation Project database (RGAP; https://rice.uga.edu). The genome sequences for the other 14 species—*O. rufipogon* (RUF), *O. nivara* (NIV), *O. glaberrima* (GLA), *O. barthii* (BAR), *O. glumaepatula*(GLU), *O. longistaminata* (LON), *O. meridionalis* (MER), *O. punctata* (PUN), *O. officinalis*(OFF), *O. eichingeri* (EIC), *O. rhizomatis* (RHI), *O. australiensis* (AUS), *O. brachyantha*(BRA), and *O. meyeriana* (MEY)—were sourced from recent publications (Supplementary Table S1) [**15,30,32,34,46,104,105**].

### 4.2. Identification, annotation, and classification of LTR-RTs

Transposable elements were identified through a combination of *de novo* prediction and homology-based searches. We used the Extensive *De-novo* TE Annotator (EDTA v2.2.2) with default parameters to generate a non-redundant TE library for each genome and to annotate all TEs [**106**]. These libraries were subsequently used to soft-mask the genome sequences with RepeatMasker v4.1.2, preparing them for segmental duplication analysis. Intact LTR-RTs were detected using LTRharvest v1.1, LTR_FINDER v1.1, and LTR_retriever v3.0.2 [**107–109**]. All intact LTR-RTs were then classified using TEsorter v1.4.7 with the REXdb v4.0 database [**10,110**]. To define families within each lineage, we performed an all-to-all BLAST comparison of the 5′ LTR sequences. A group was accepted as a family if >50% of its 5′ LTRs shared >80% sequence identity [**103**].

### 4.3. Population genetic analysis of LTR-RT lineages

For each lineage, a consensus sequence was constructed from all identified full-length intact LTR-RTs using MAFFT v7.525 and the *consensus* function in TEpop [**19,84,111**]. Members of major LTR-RT families were aligned to their respective consensus sequences using the EMBOSS *water* program with a gap opening penalty of 50 and a gap extension penalty of 0.1. These alignments were converted into variant call format (VCF) files using the *pair_to_vcf_2024.pl* script from TEpop [**19**]. Only nucleotide substitutions present in at least 80% of all copies were retained for analysis [**19**]. Principal component analysis (PCA) was performed on these VCF files, and results were visualized using the R packages vcfR v1.15.0, ggplot2 v4.0.1, ggfortify v0.1.2, cowplot v1.1.3, dplyr v1.1.4, and plotly v4.11.0. Population genetic statistics (FST, π, and Tajima’s D) were calculated for intact LTR-RTs across species using VCFtools v0.1.14 [**112**].

### 4.4. Identification of solo LTRs and truncated LTRs

Solo and truncated LTRs were identified by aligning the LTR sequences from intact LTR-RTs against the genomes using BLASTN, excluding regions already annotated as intact LTR-RTs, coding sequences (CDS), or non-LTR-RT TEs. Alignments covering ≥80% of the query length with ≥80% similarity were classified as solo LTRs. Regions with ≥80% alignment length but similarity between 50% and 80% were classified as truncated LTRs [**14,35,43**]. Birth counts of LTR-RTs were derived from matrix population models [**113**] and defined as the sum of intact LTR-RTs, solo LTRs, and truncated LTRs. The deletion rate was calculated as the ratio of solo LTRs to intact LTR-RTs. LTR-RT families were then categorized as follows: high removal rate (deletion rate ≥ 3), medium removal rate (deletion rate < 3 and (solo + truncated LTRs)/intact LTR-RTs ≥ 1), and low removal rate ((solo + truncated LTRs)/intact LTR-RTs < 1).

### 4.5. Identification of structural variation

The genomes of the 14 other *Oryza* species were aligned to the *O. sativa* reference genome using minimap2 v2.30 with the “-ax asm20” parameter set [**114**]. We identified structural variations (SVs) using SyRI v1.7.1 with default parameters [**49**].

### 4.6. Identification of segmental duplications and LTR-RT-associated SDs

Following RepeatMasker soft-masking, tandem repeats were identified using Tandem Repeats Finder (TRF) with parameters: 2 7 7 80 10 50 500 [**115**]. These repeat regions were then hard-masked. Segmental duplications (SDs) were detected using BISER v1.4 [**116**], retaining hits with ≥90% sequence identity. SDs potentially driven by LTR-RTs were identified based on their breakpoint junctions: each SD has four 5-bp junction sequences flanking its breakpoints. SDs with two or more junctions overlapping LTR-RT sequences were classified as LTR-RT-associated (**Fig. 4e**).

### 4.7. Estimation of LTR-RT insertion time and SD formation time

For each intact LTR-RT, the Kimura 2-parameter (K2P) distance (*d*) between its two LTRs was calculated using MEGA11 with the complete deletion option [**117**]. Insertion time (*t*) was estimated as *t = d / 2r*, where *r* is the neutral substitution rate (1.3 × 10LL substitutions per site per year) [**26,94**]. For SDs, the divergence (*K*) was calculated using the Jukes-Cantor model: *K = -(3/4) * ln(1 - (4/3) * (1 - identity))* [**118**]. The formation time (*T*) of an SD was then estimated as *T = K / 2μ*, where μ is the nucleotide substitution rate for rice (1.3 × 10LL per bp per year) [**119**].

### 4.8. Reconstruction of 15 *Oryza* diploid species phylogeny

*L. perreri* was chosen as the outgroup for inferring the species phylogeny. The 6,692 single-copy orthologous genes identified by Orthofinder were aligned using MAFFT [**120**]. The phylogenetic tree of Oryza was constructed using IQTree2 v.2.4.0 [**121**]. The Divergence time was estimated using the MCMCTree program in PAML v.4.10.9 [**122**].

### 4.9. Construction of the phylogenetic tree of LTR-RTs across *Oryza* diploid species phylogeny

The phylogenetic trees of 10,876 Ty1/Copia LTR-RTs and 29,178 Ty3/Gypsy LTR-RTs were constructed based on the RT sequences. These sequences were aligned using MAFFT. The phylogenetic trees of these LTR-RTs superfamilies were constructed using IQTree2. The phylogenetic trees of LTR-RTs from distinct lineages were constructed based on the 5’ LTR sequences. These sequences were aligned using MAFFT. The phylogenetic trees of these LTR-RTs superfamilies were constructed using IQTree2.

## Supporting information

Supplementary Table S4 The Fst of LTR-RTs from different lineages in Oryza diploid species

Supplementary Table S11 The information of counts and deletion rate of LTR-RTs in Oryza diploid species

## Author Contributions

L.-Z.G. designed the study. R.-J.X. analyzed the data. R.-J.X. drafted the manuscript. L.-Z.G. revised the manuscript.

## Acknowledgments

This work was supported by a start-up grant from Hainan University (to L.G.).

