## Supplementary Figures S1-S56 for "Lineage-specific evolution of LTR retrotransposons under natural selection drives genomic divergence in diploid *Oryza* species"

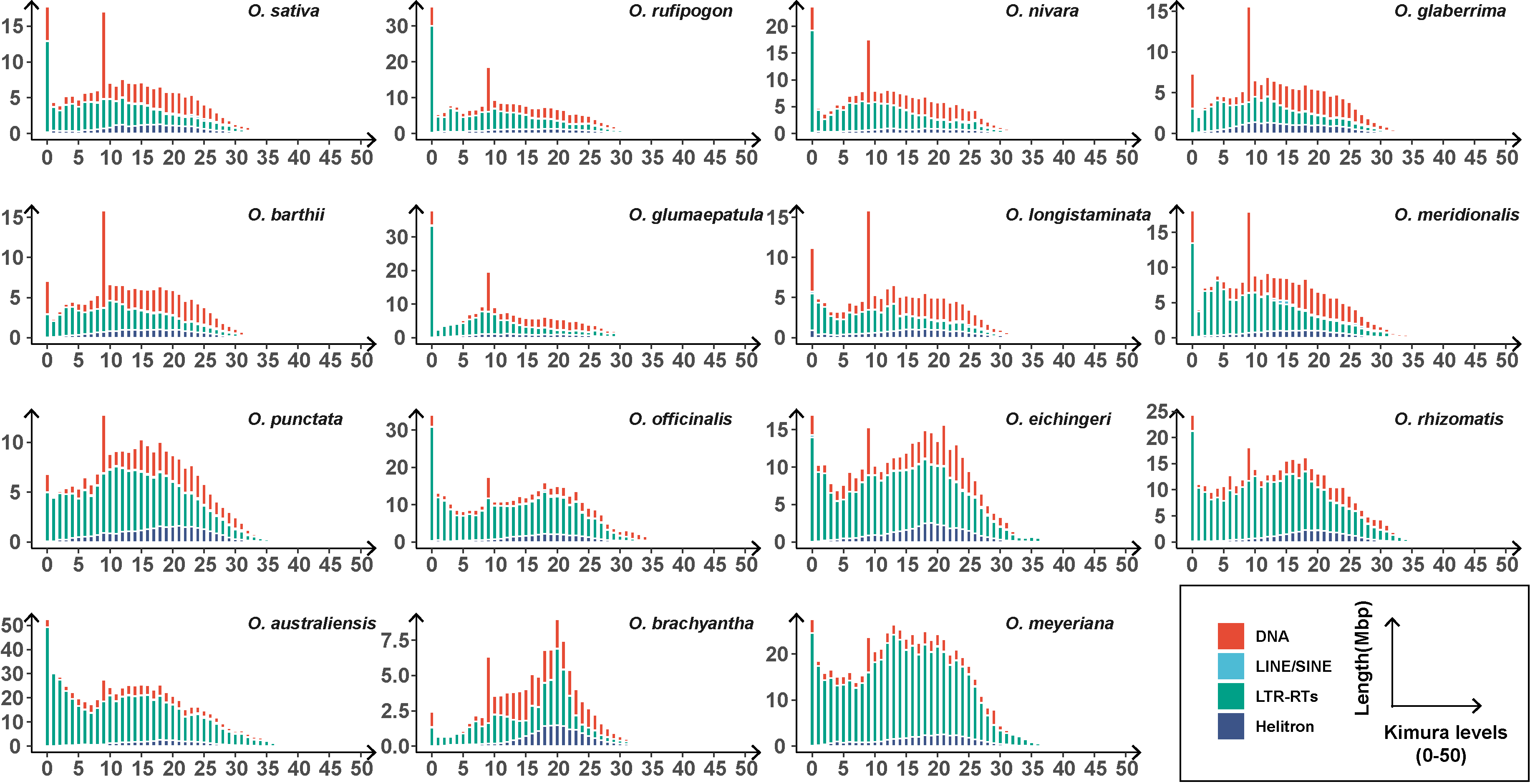

**Fig. S1.** **Kimura distance-based divergence analysis of TEs in Oryza diploid species genomes.** These graphs represent the total length for different types of TEs (LTR-RTs and LINE/SINE retrotransposons; DNA transposons and Helitron transposons) in the analyzed genomes.

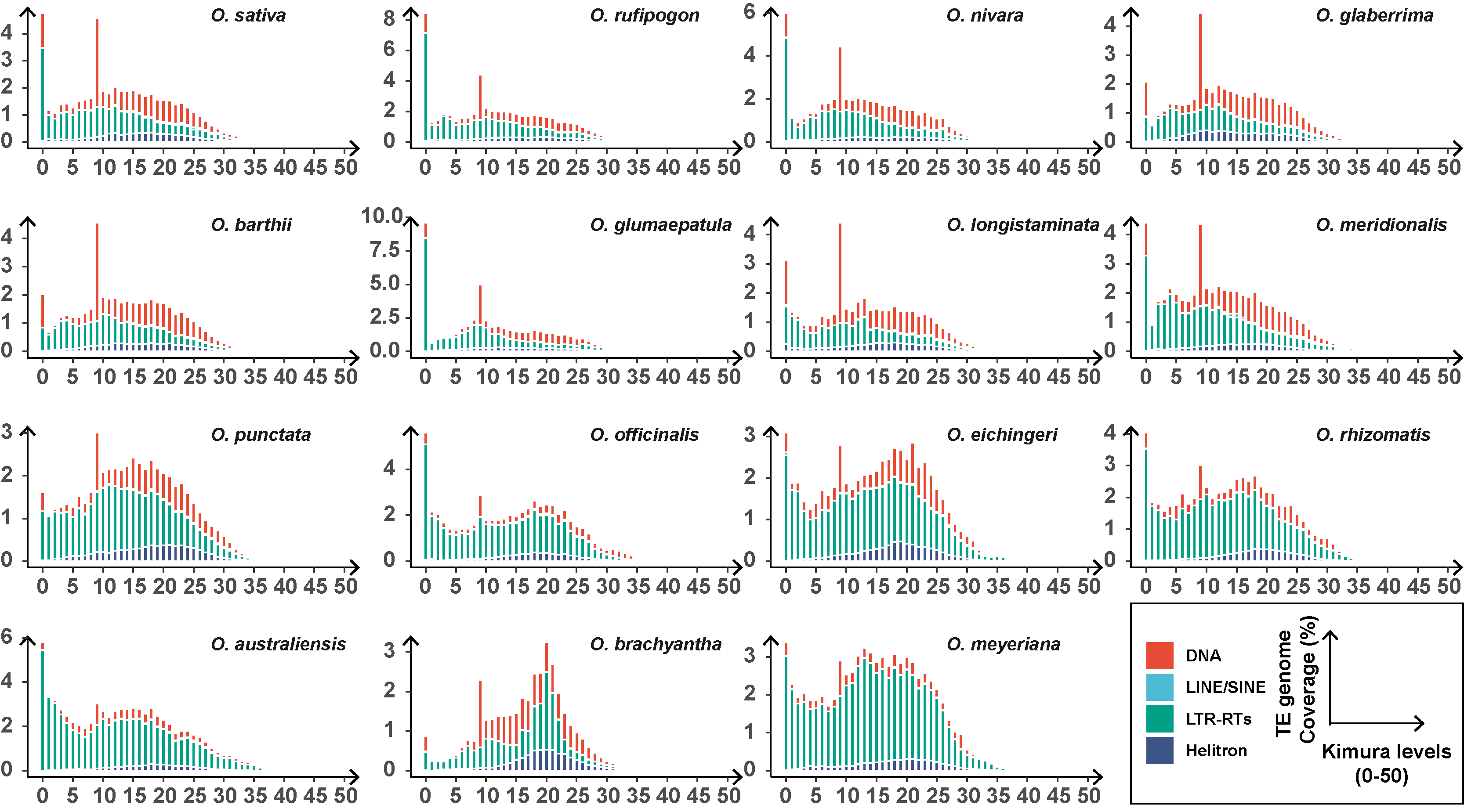

**Fig. S2.** **Kimura distance-based divergence analysis of TEs in Oryza diploid species genomes.** These graphs represent the genome coverage for different types of TEs (LTR-RTs and LINE/SINE retrotransposons; DNA transposons and Helitron transposons) in the analyzed genomes.

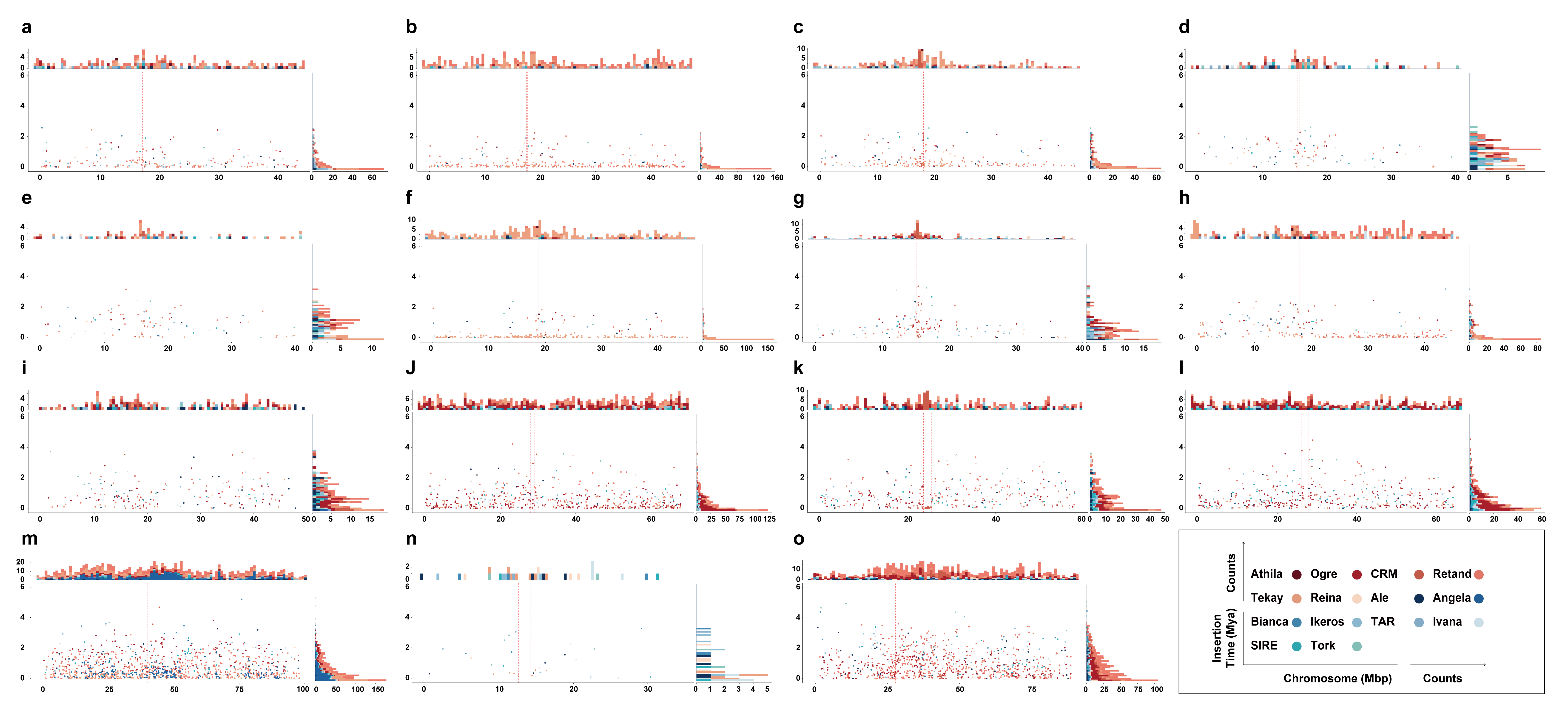

**Fig. S3.** **The spatiotemporal distribution of intact LTR-RTs from chromosome 1 of diploid Oryza species.** From **a** to **o** represents *O. sativa*, *O. rufipogon*, *O. nivara*, *O. glaberrima*, *O. barthii*, *O. glumaepatula*, *O. longistaminata*, *O. meridionalis*, *O. punctata*, *O. officinalis*, *O. eichingeri*, *O. rhizomatis*, *O. australiensis*, *O. brachyantha*, *O. meyeriana*, respectively.

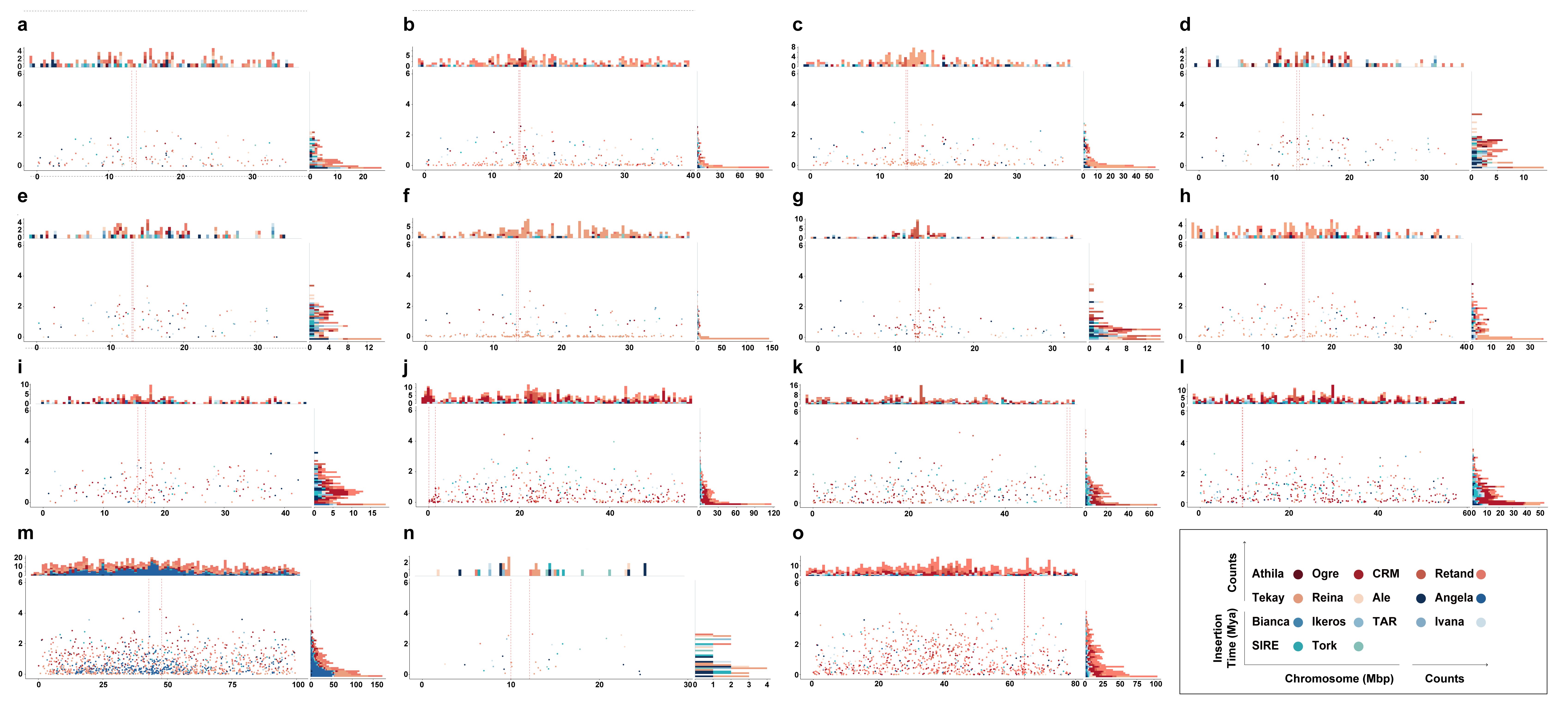

**Fig. S4.** **The spatiotemporal distribution of intact LTR-RTs from chromosome 2 of diploid Oryza species.** From **a** to **o** represents *O. sativa*, *O. rufipogon*, *O. nivara*, *O. glaberrima*, *O. barthii*, *O. glumaepatula*, *O. longistaminata*, *O. meridionalis*, *O. punctata*, *O. officinalis*, *O. eichingeri*, *O. rhizomatis*, *O. australiensis*, *O. brachyantha*, *O. meyeriana*, respectively.

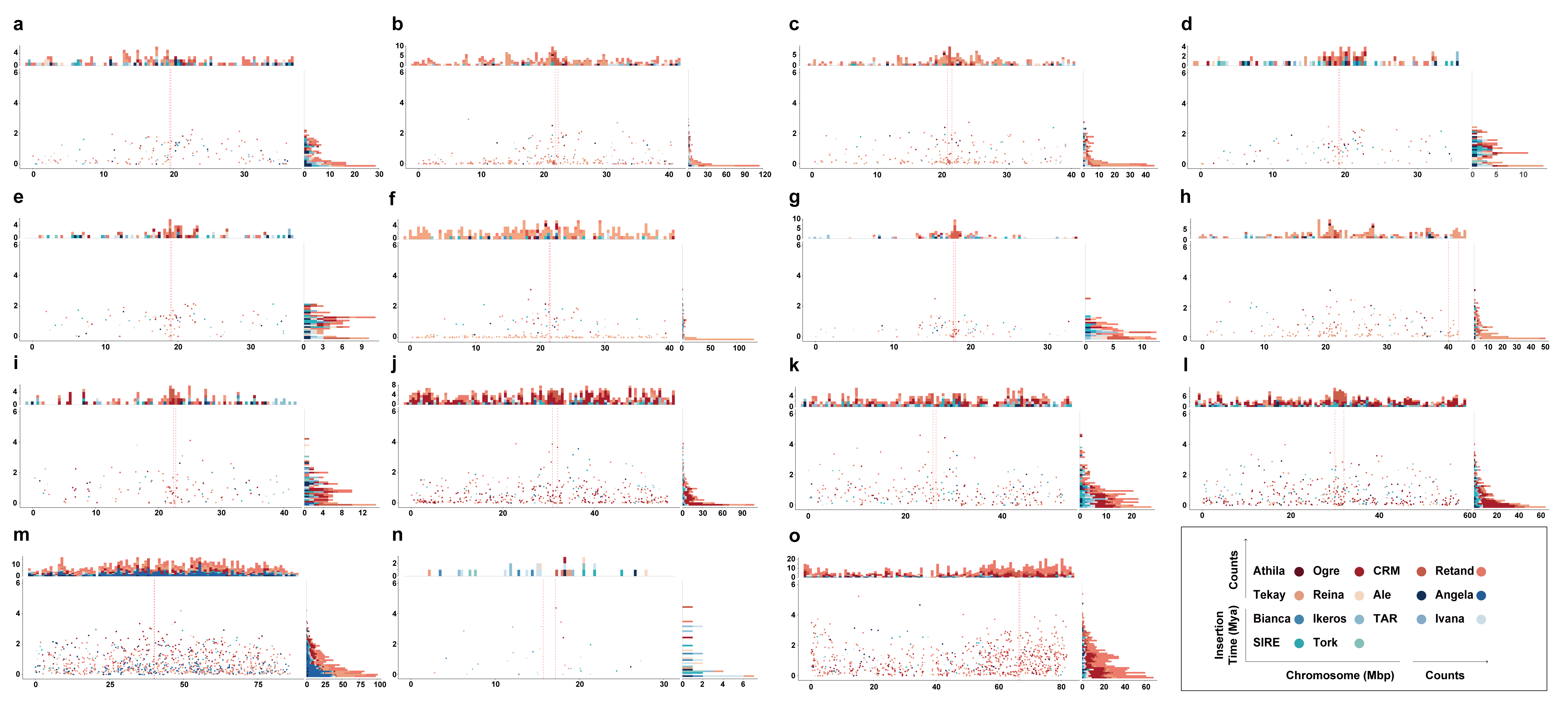

**Fig. S5.** **The spatiotemporal distribution of intact LTR-RTs from chromosome 3 of diploid Oryza species.** From **a** to **o** represents *O. sativa*, *O. rufipogon*, *O. nivara*, *O. glaberrima*, *O. barthii*, *O. glumaepatula*, *O. longistaminata*, *O. meridionalis*, *O. punctata*, *O. officinalis*, *O. eichingeri*, *O. rhizomatis*, *O. australiensis*, *O. brachyantha*, *O. meyeriana*, respectively.

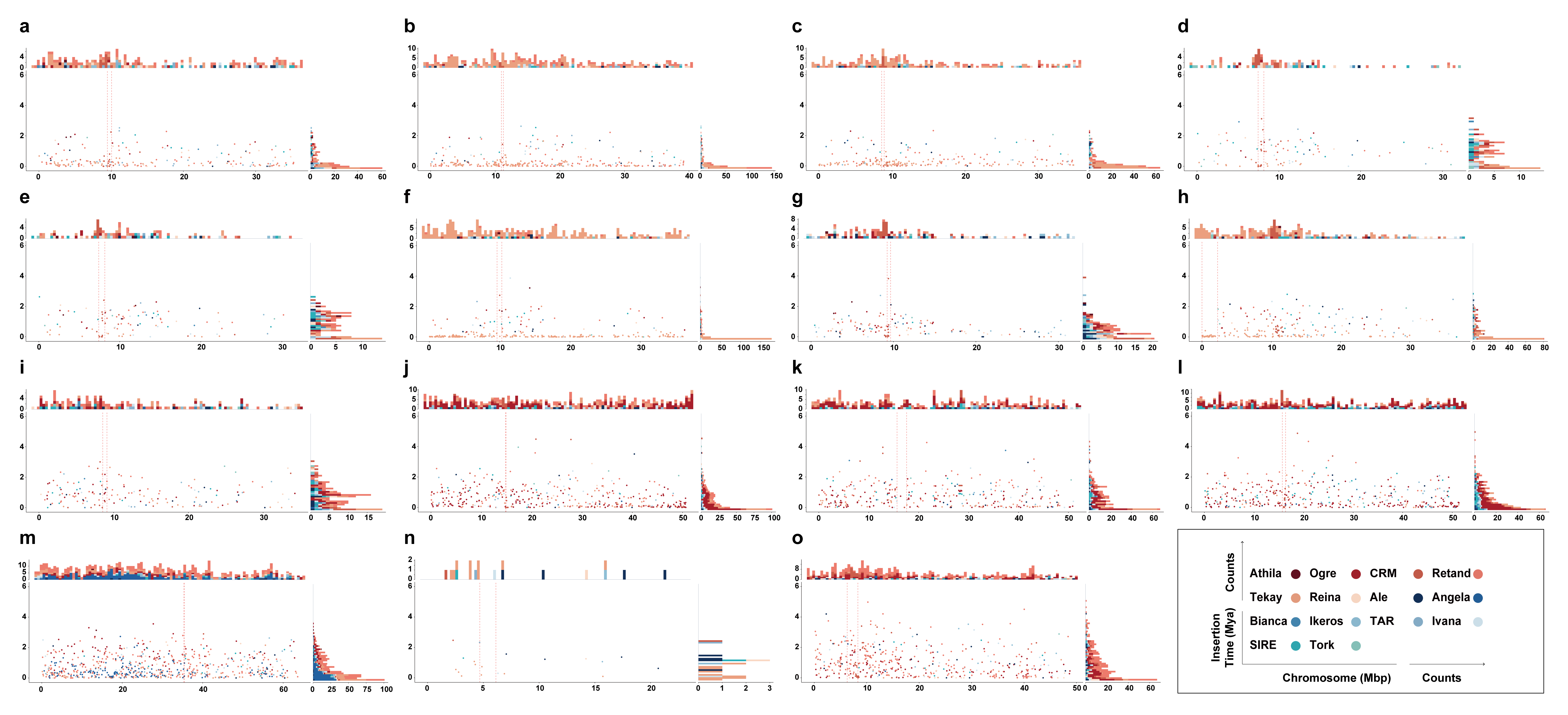

**Fig. S6.** **The spatiotemporal distribution of intact LTR-RTs from chromosome 4 of diploid Oryza species.** From **a** to **o** represents *O. sativa*, *O. rufipogon*, *O. nivara*, *O. glaberrima*, *O. barthii*, *O. glumaepatula*, *O. longistaminata*, *O. meridionalis*, *O. punctata*, *O. officinalis*, *O. eichingeri*, *O. rhizomatis*, *O. australiensis*, *O. brachyantha*, *O. meyeriana*, respectively.

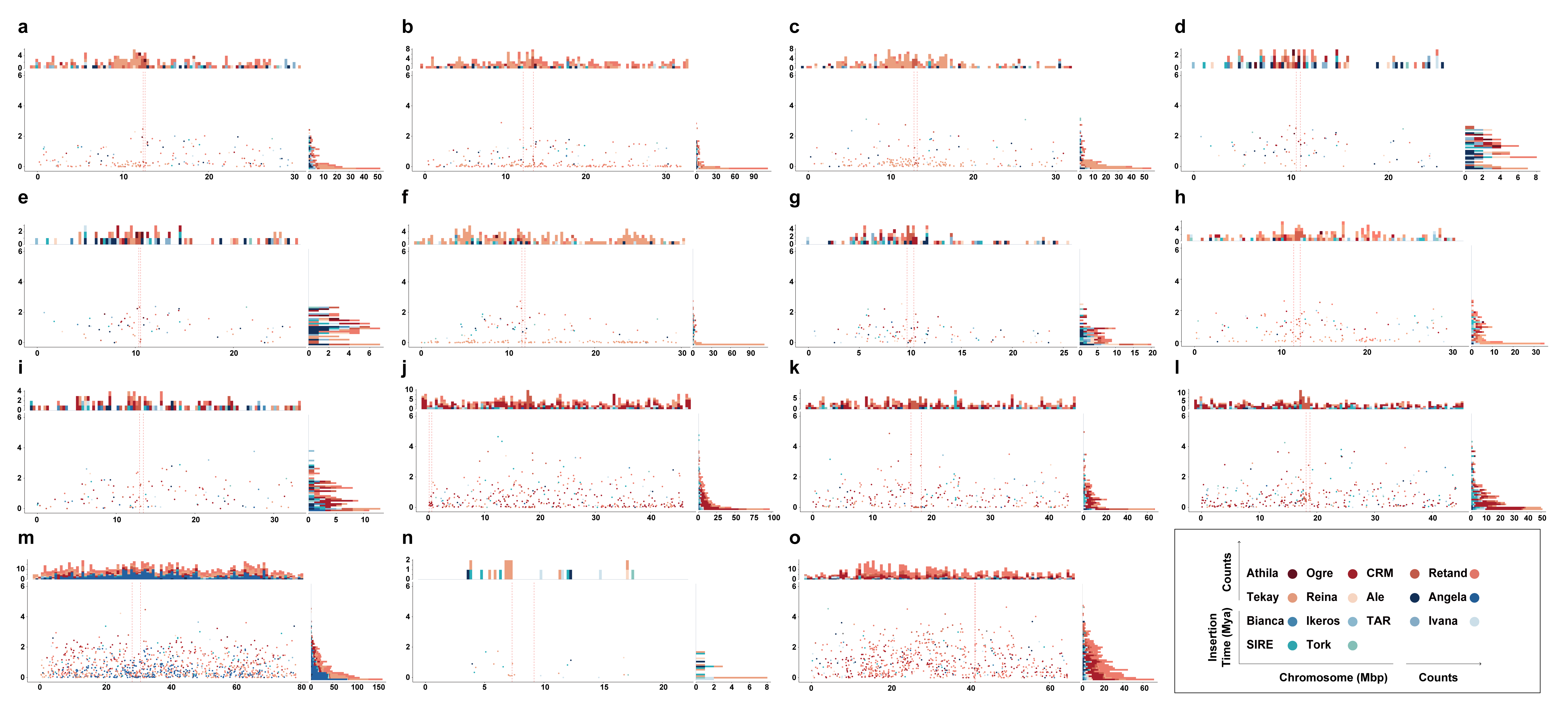

**Fig. S7.** **The spatiotemporal distribution of intact LTR-RTs from chromosome 5 of diploid Oryza species.** From **a** to **o** represents *O. sativa*, *O. rufipogon*, *O. nivara*, *O. glaberrima*, *O. barthii*, *O. glumaepatula*, *O. longistaminata*, *O. meridionalis*, *O. punctata*, *O. officinalis*, *O. eichingeri*, *O. rhizomatis*, *O. australiensis*, *O. brachyantha*, *O. meyeriana*, respectively.

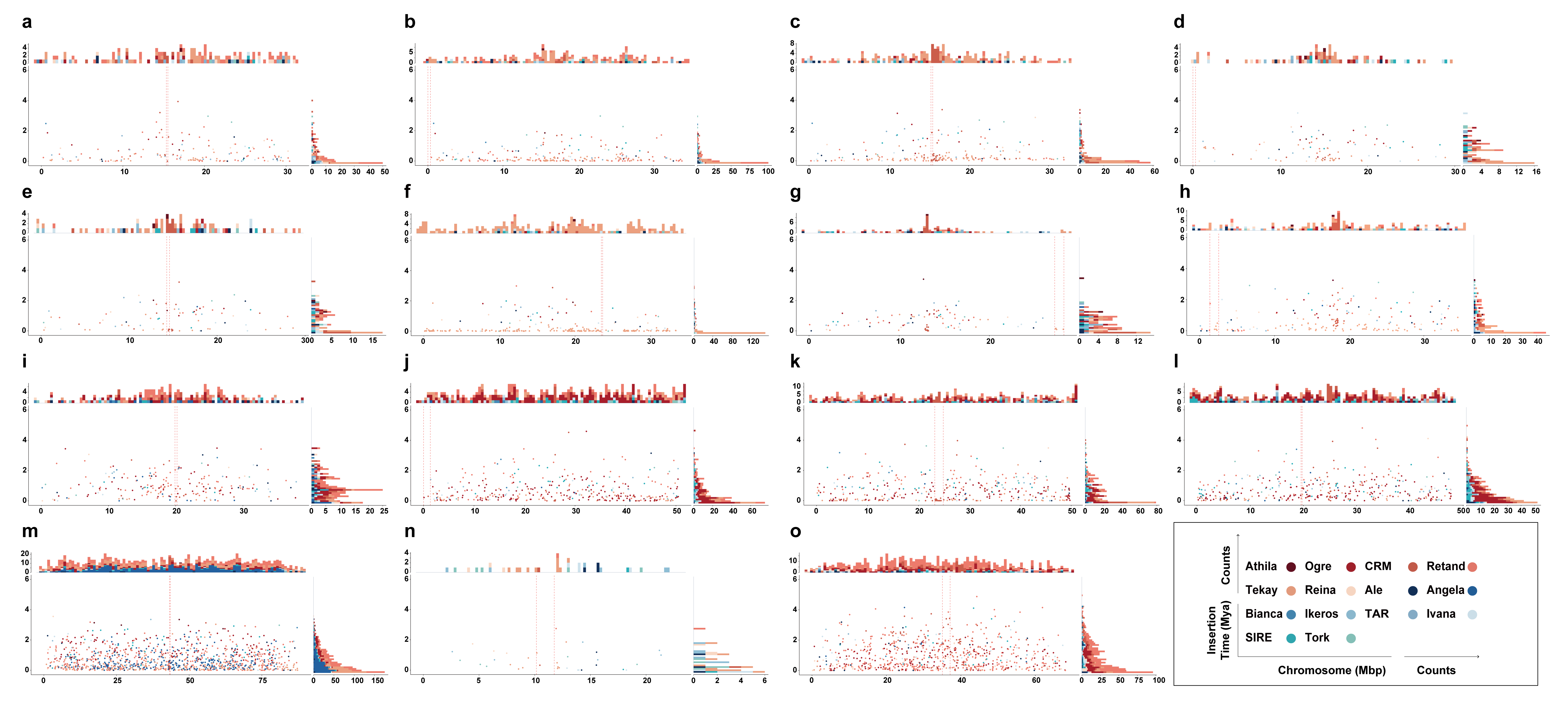

**Fig. S8.** **The spatiotemporal distribution of intact LTR-RTs from chromosome 6 of diploid Oryza species.** From **a** to **o** represents *O. sativa*, *O. rufipogon*, *O. nivara*, *O. glaberrima*, *O. barthii*, *O. glumaepatula*, *O. longistaminata*, *O. meridionalis*, *O. punctata*, *O. officinalis*, *O. eichingeri*, *O. rhizomatis*, *O. australiensis*, *O. brachyantha*, *O. meyeriana*, respectively.

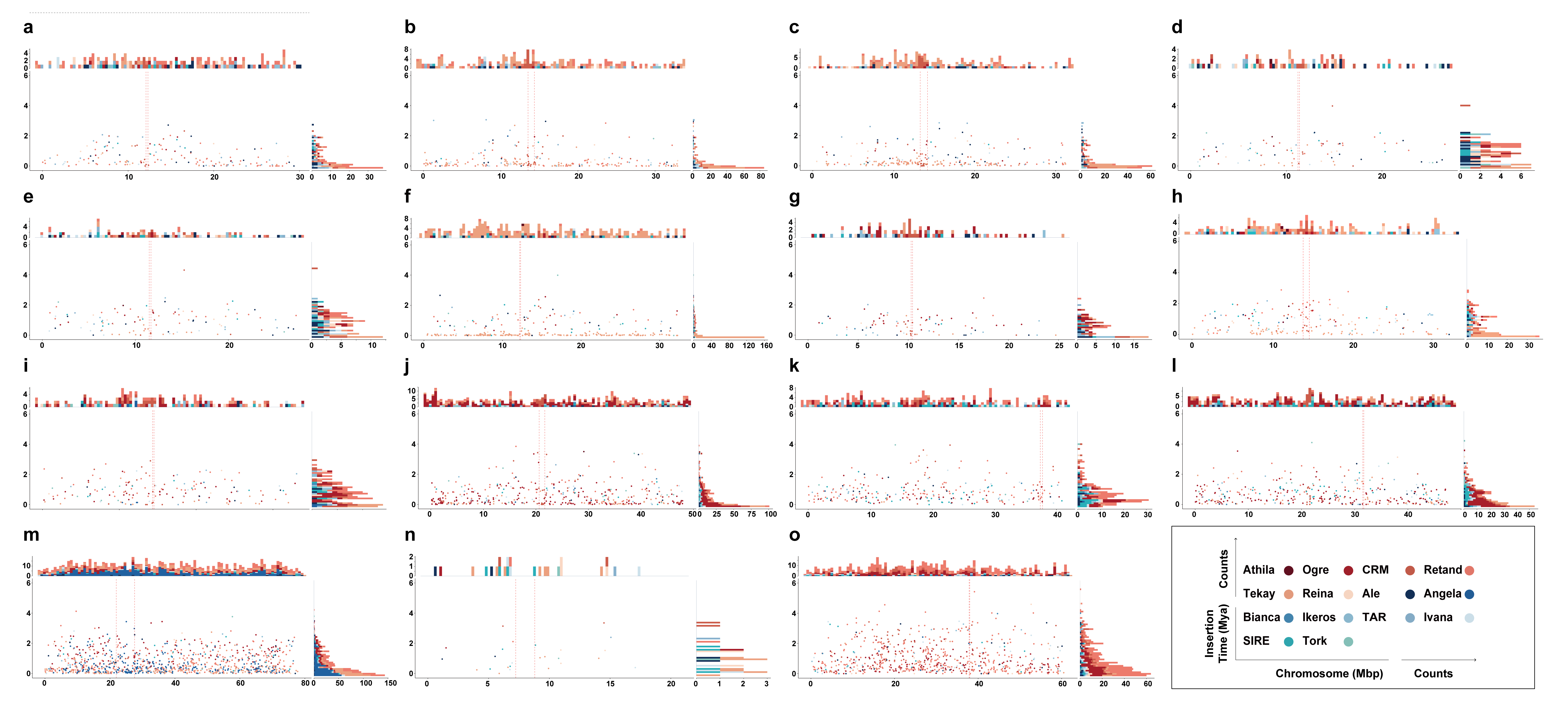

**Fig. S9.** **The spatiotemporal distribution of intact LTR-RTs from chromosome 7 of diploid Oryza species.** From **a** to **o** represents *O. sativa*, *O. rufipogon*, *O. nivara*, *O. glaberrima*, *O. barthii*, *O. glumaepatula*, *O. longistaminata*, *O. meridionalis*, *O. punctata*, *O. officinalis*, *O. eichingeri*, *O. rhizomatis*, *O. australiensis*, *O. brachyantha*, *O. meyeriana*, respectively.

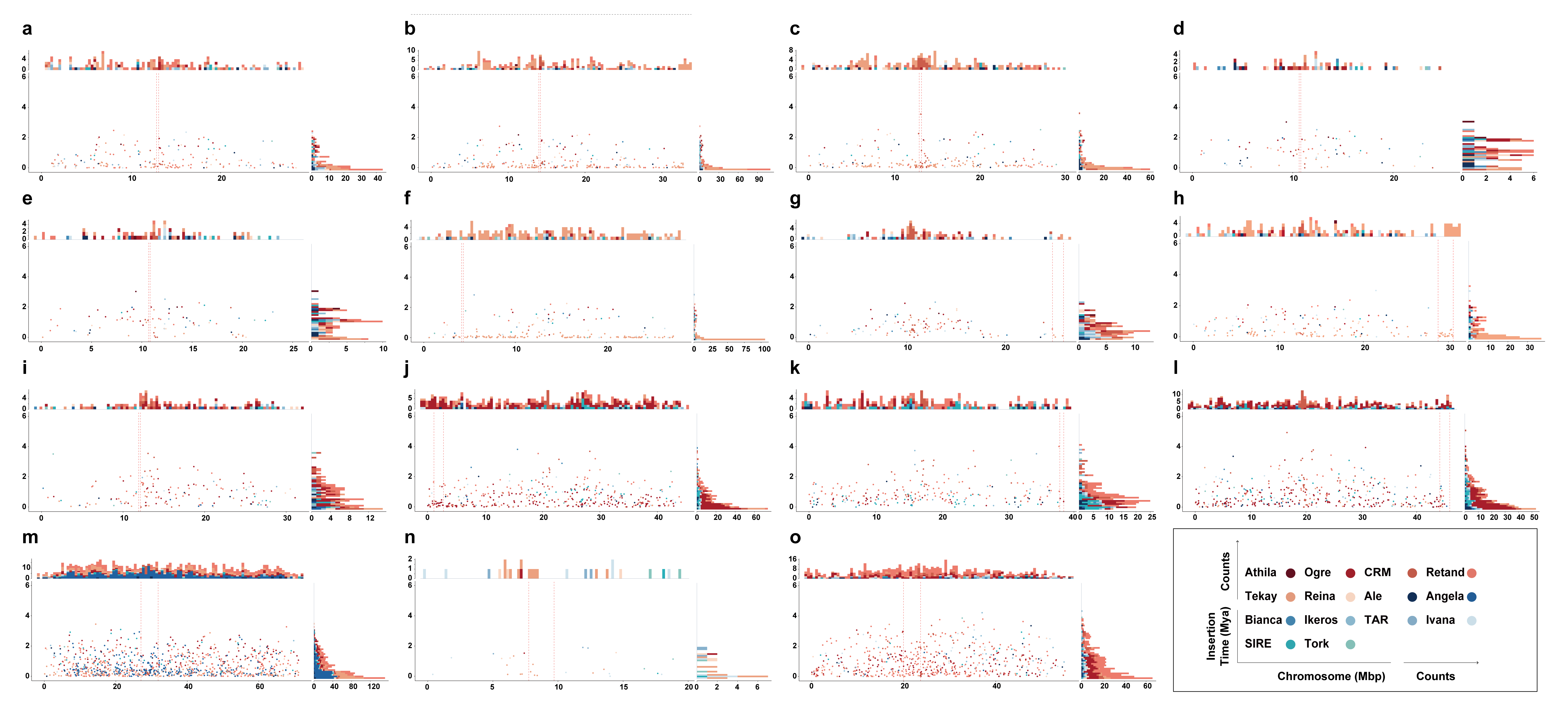

**Fig. S10.** **The spatiotemporal distribution of intact LTR-RTs from chromosome 8 of diploid Oryza species.** From **a** to **o** represents *O. sativa*, *O. rufipogon*, *O. nivara*, *O. glaberrima*, *O. barthii*, *O. glumaepatula*, *O. longistaminata*, *O. meridionalis*, *O. punctata*, *O. officinalis*, *O. eichingeri*, *O. rhizomatis*, *O. australiensis*, *O. brachyantha*, *O. meyeriana*, respectively.

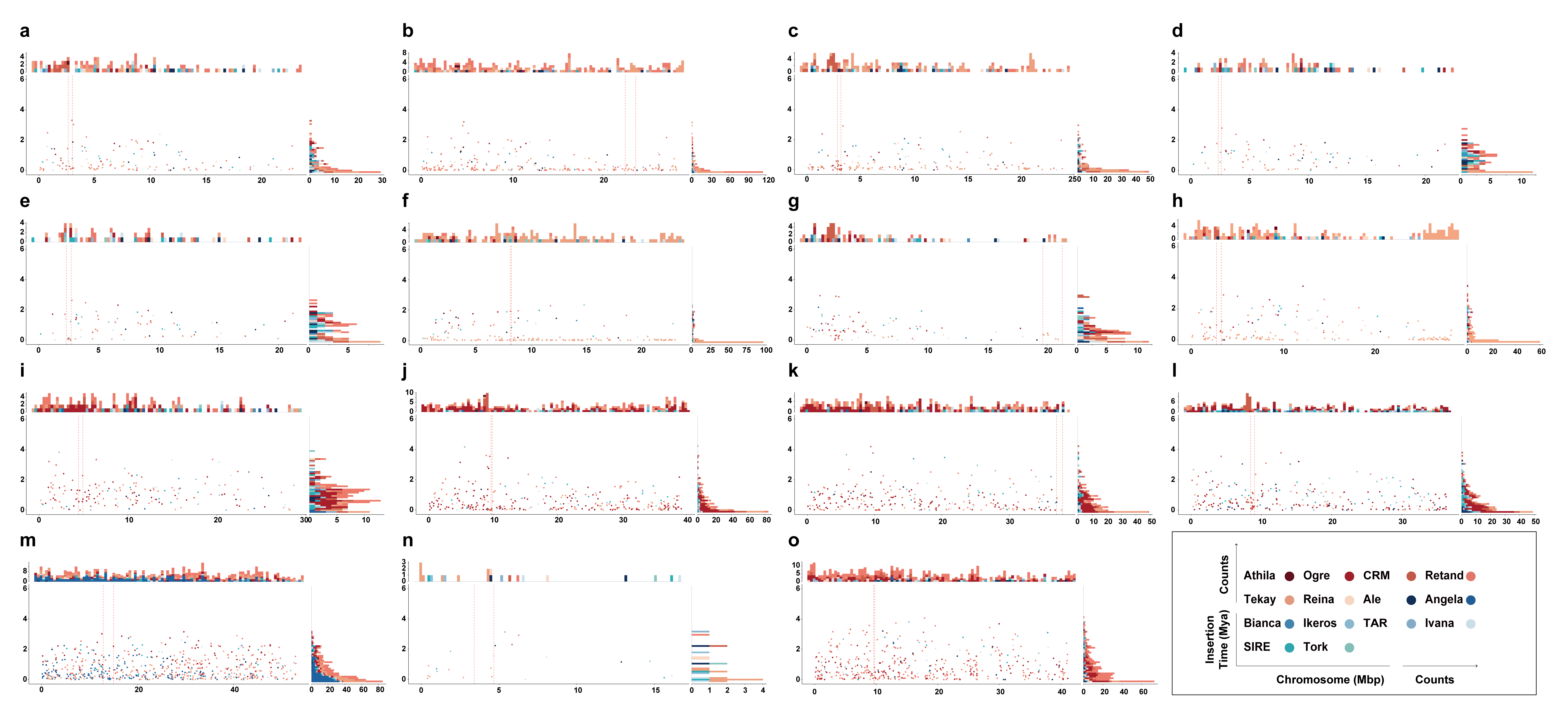

**Fig. S11.** **The spatiotemporal distribution of intact LTR-RTs from chromosome 9 of diploid Oryza species.** From **a** to **o** represents *O. sativa*, *O. rufipogon*, *O. nivara*, *O. glaberrima*, *O. barthii*, *O. glumaepatula*, *O. longistaminata*, *O. meridionalis*, *O. punctata*, *O. officinalis*, *O. eichingeri*, *O. rhizomatis*, *O. australiensis*, *O. brachyantha*, *O. meyeriana*, respectively.

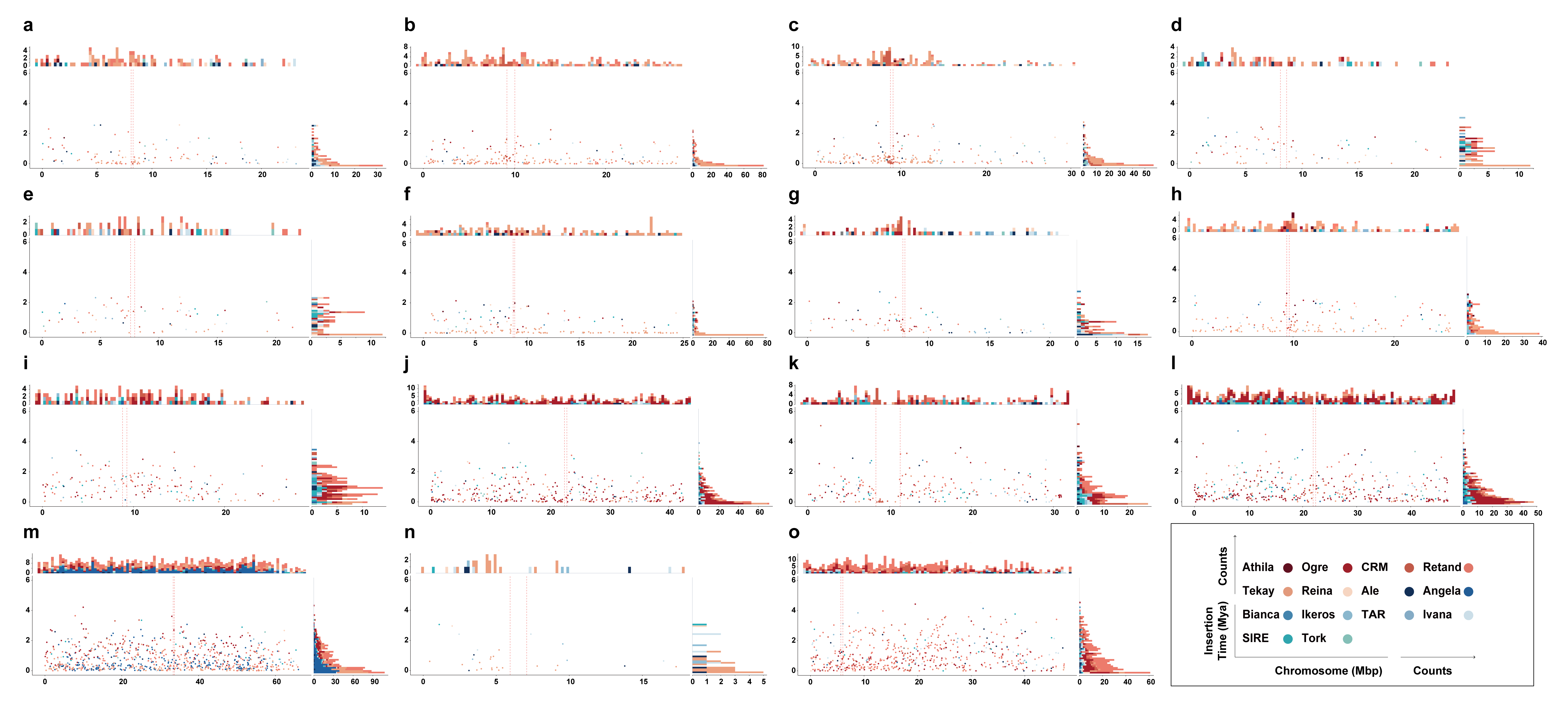

**Fig. S12.** **The spatiotemporal distribution of intact LTR-RTs from chromosome 10 of diploid Oryza species.** From **a** to **o** represents *O. sativa*, *O. rufipogon*, *O. nivara*, *O. glaberrima*, *O. barthii*, *O. glumaepatula*, *O. longistaminata*, *O. meridionalis*, *O. punctata*, *O. officinalis*, *O. eichingeri*, *O. rhizomatis*, *O. australiensis*, *O. brachyantha*, *O. meyeriana*, respectively.

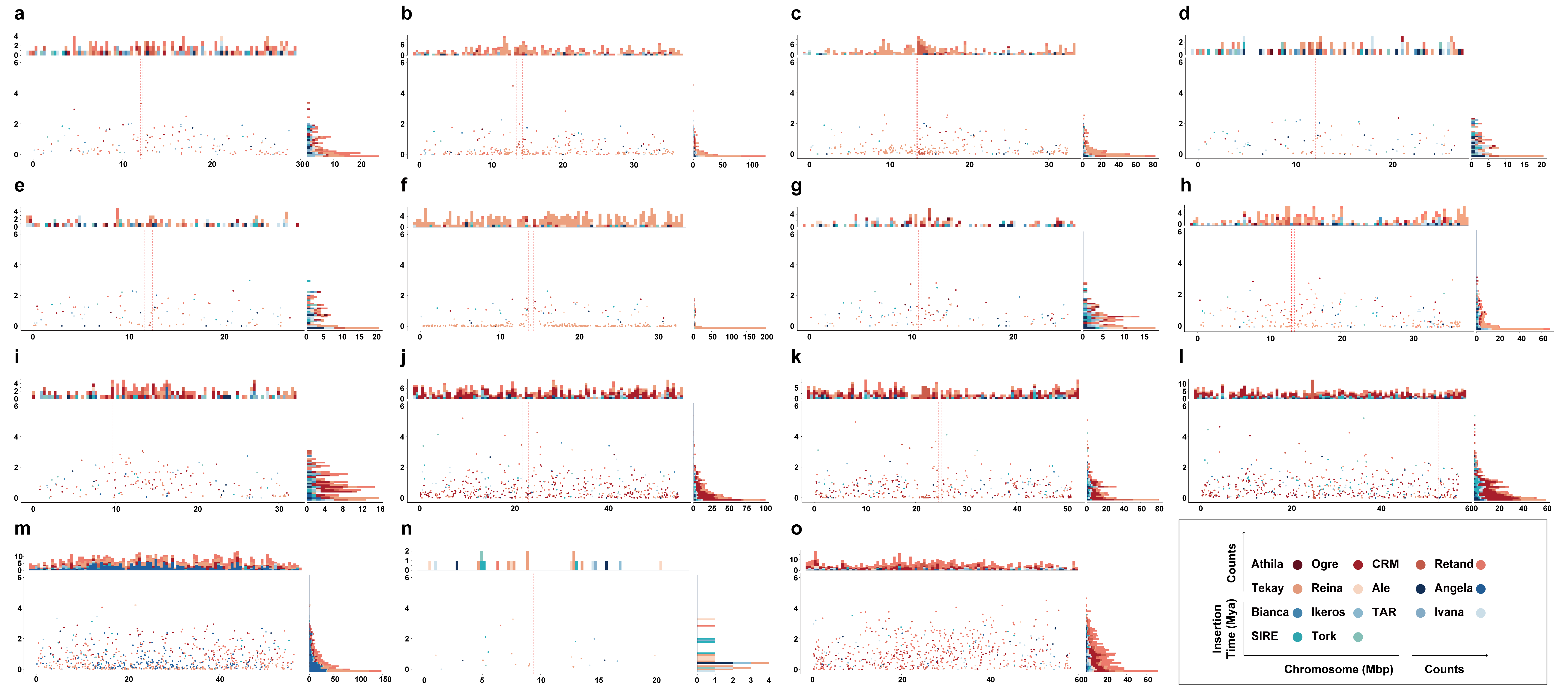

**Fig. S13.** **The spatiotemporal distribution of intact LTR-RTs from chromosome 11 of diploid Oryza species.** From **a** to **o** represents *O. sativa*, *O. rufipogon*, *O. nivara*, *O. glaberrima*, *O. barthii*, *O. glumaepatula*, *O. longistaminata*, *O. meridionalis*, *O. punctata*, *O. officinalis*, *O. eichingeri*, *O. rhizomatis*, *O. australiensis*, *O. brachyantha*, *O. meyeriana*, respectively.

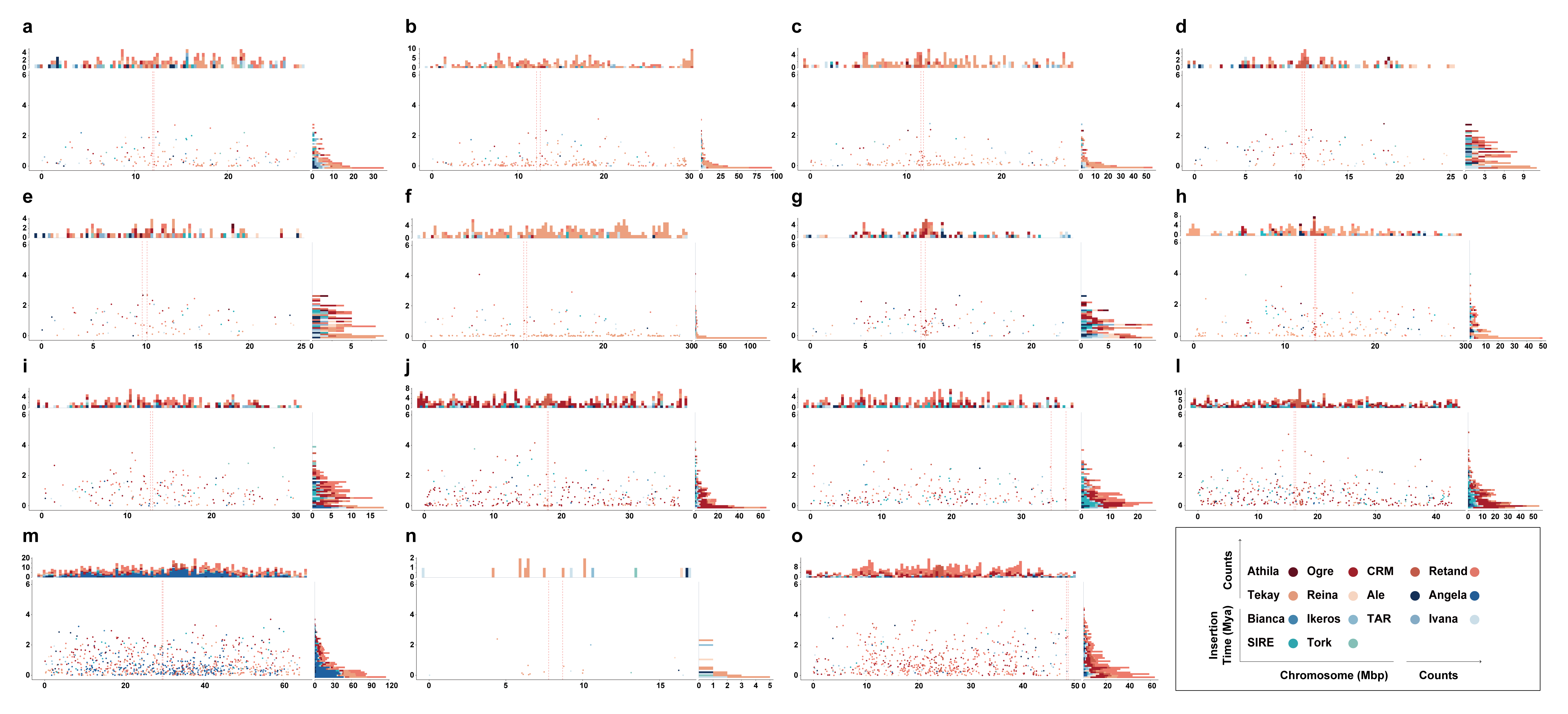

**Fig. S14.** **The spatiotemporal distribution of intact LTR-RTs from chromosome 12 of diploid Oryza species.** From **a** to **o** represents *O. sativa*, *O. rufipogon*, *O. nivara*, *O. glaberrima*, *O. barthii*, *O. glumaepatula*, *O. longistaminata*, *O. meridionalis*, *O. punctata*, *O. officinalis*, *O. eichingeri*, *O. rhizomatis*, *O. australiensis*, *O. brachyantha*, *O. meyeriana*, respectively.

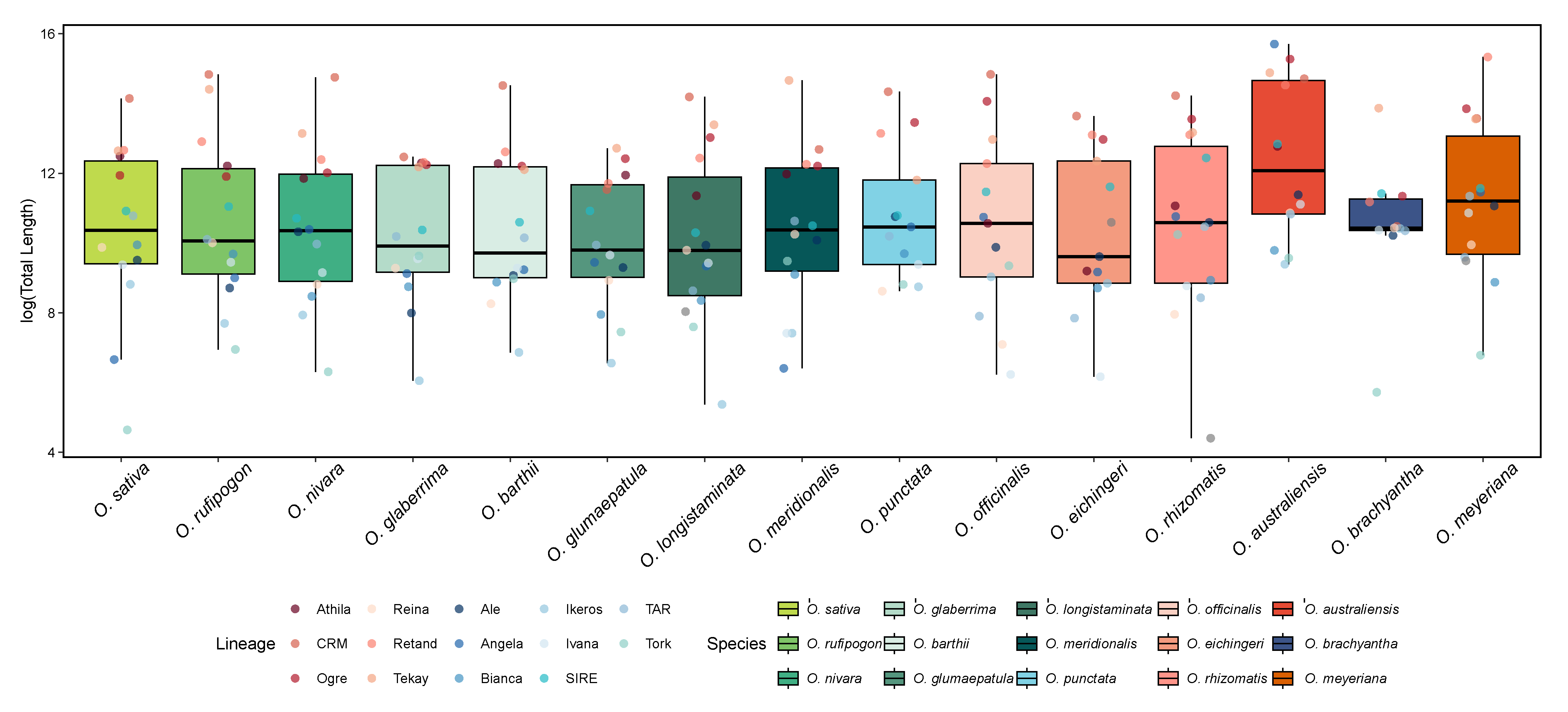

**Fig. S15.** **The total length of LTR-RTs in the centromeric regions of diploid Oryza species.**

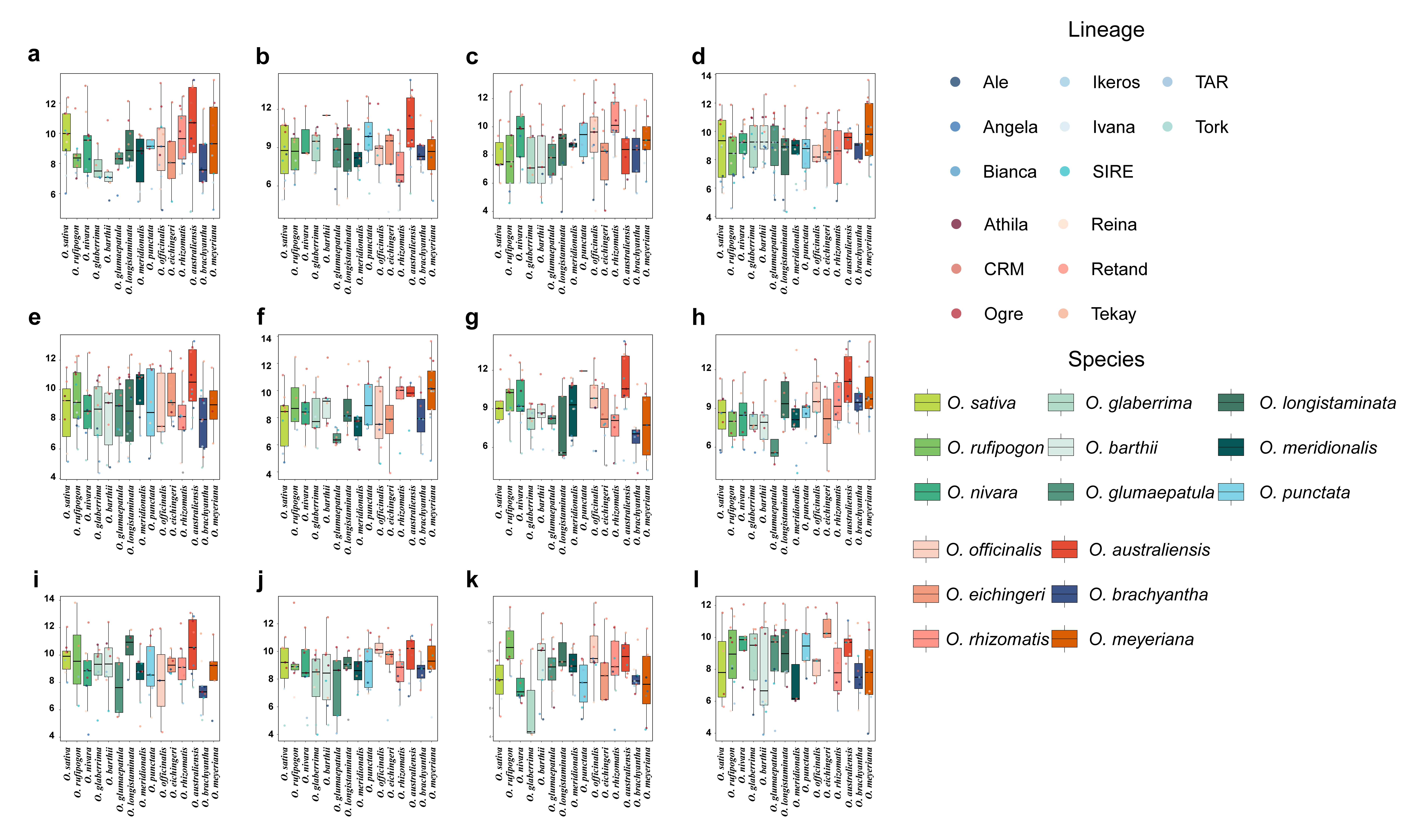

**Fig. S16.** **The total length of LTR-RTs in the centromeric region in different chromosomes across different species.** From **a** to **l** represented chromosome 1- chromosome 12, respectively.

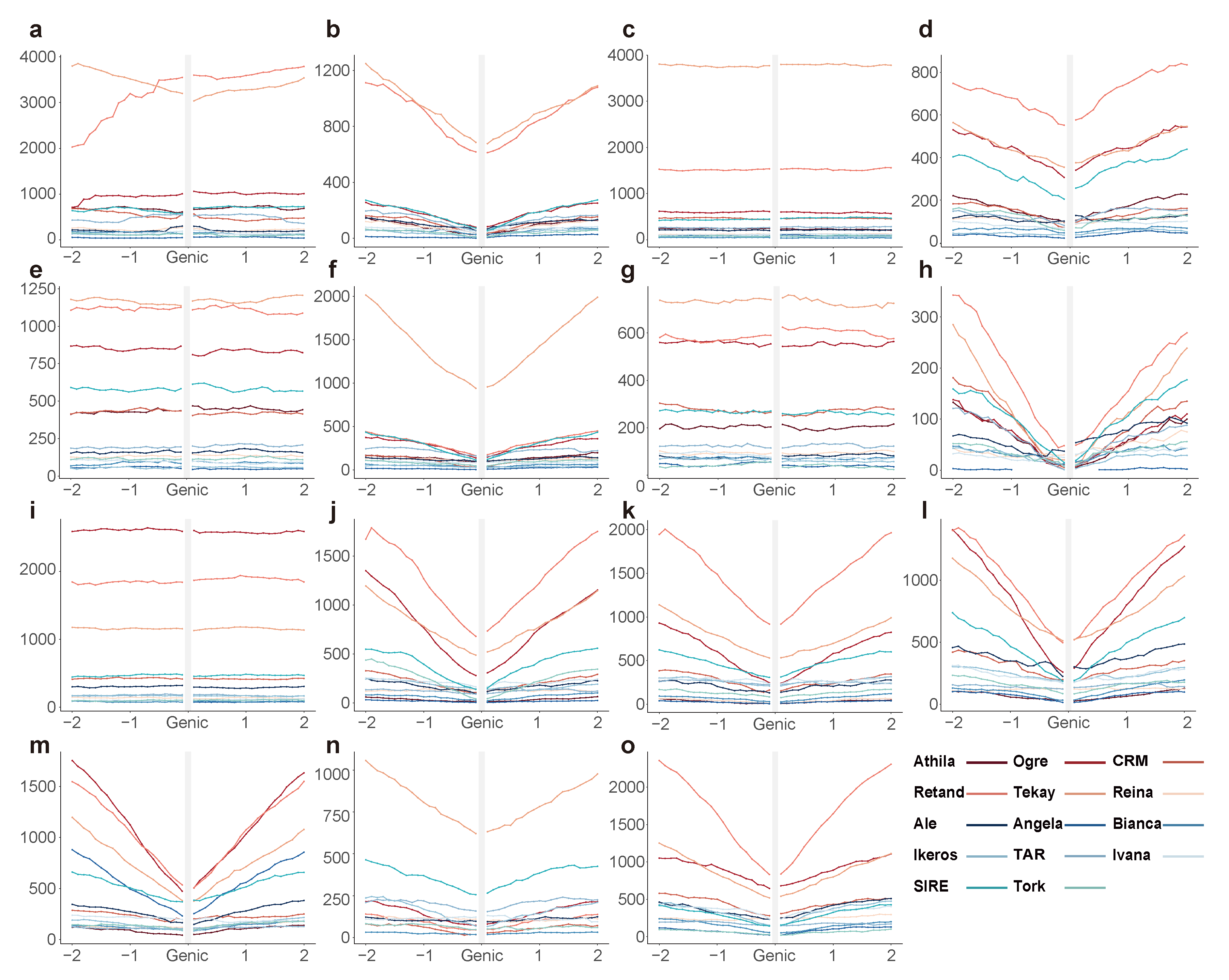

**Fig. S17.** **The LTR-RT content in the 2kb upstream and downstream flanking regions of genes in diploid Oryza species.** From **a** to **o** represents *O. sativa*, *O. rufipogon*, *O. nivara*, *O. glaberrima*, *O. barthii*, *O. glumaepatula*, *O. longistaminata*, *O. meridionalis*, *O. punctata*, *O. officinalis*, *O. eichingeri*, *O. rhizomatis*, *O. australiensis*, *O. brachyantha*, *O. meyeriana*, respectively.

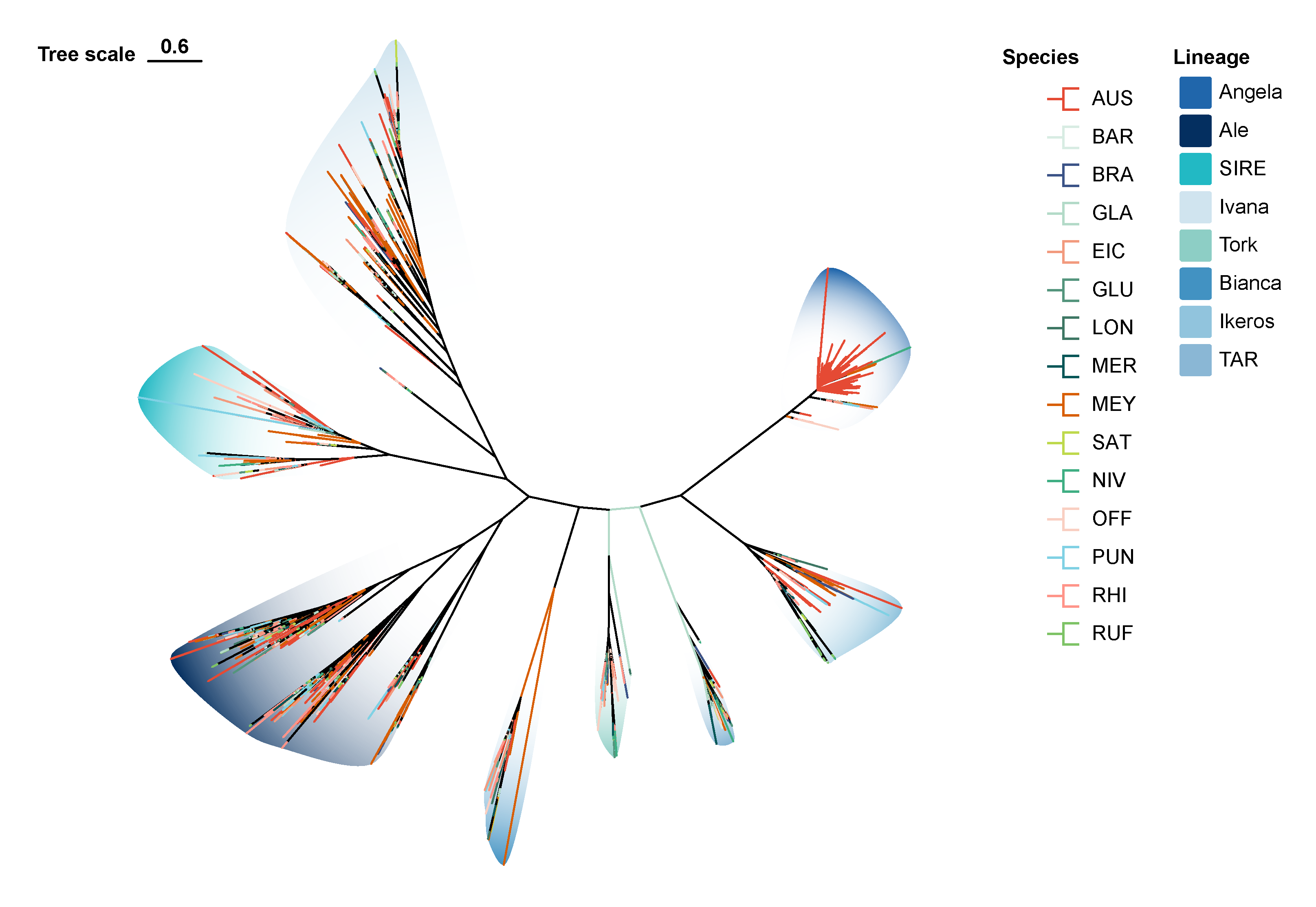

**Fig. S18.** **Unrooted gene tree of intact LTR-RTs from Ty1/Copia superfamily in diploid Oryza species.**

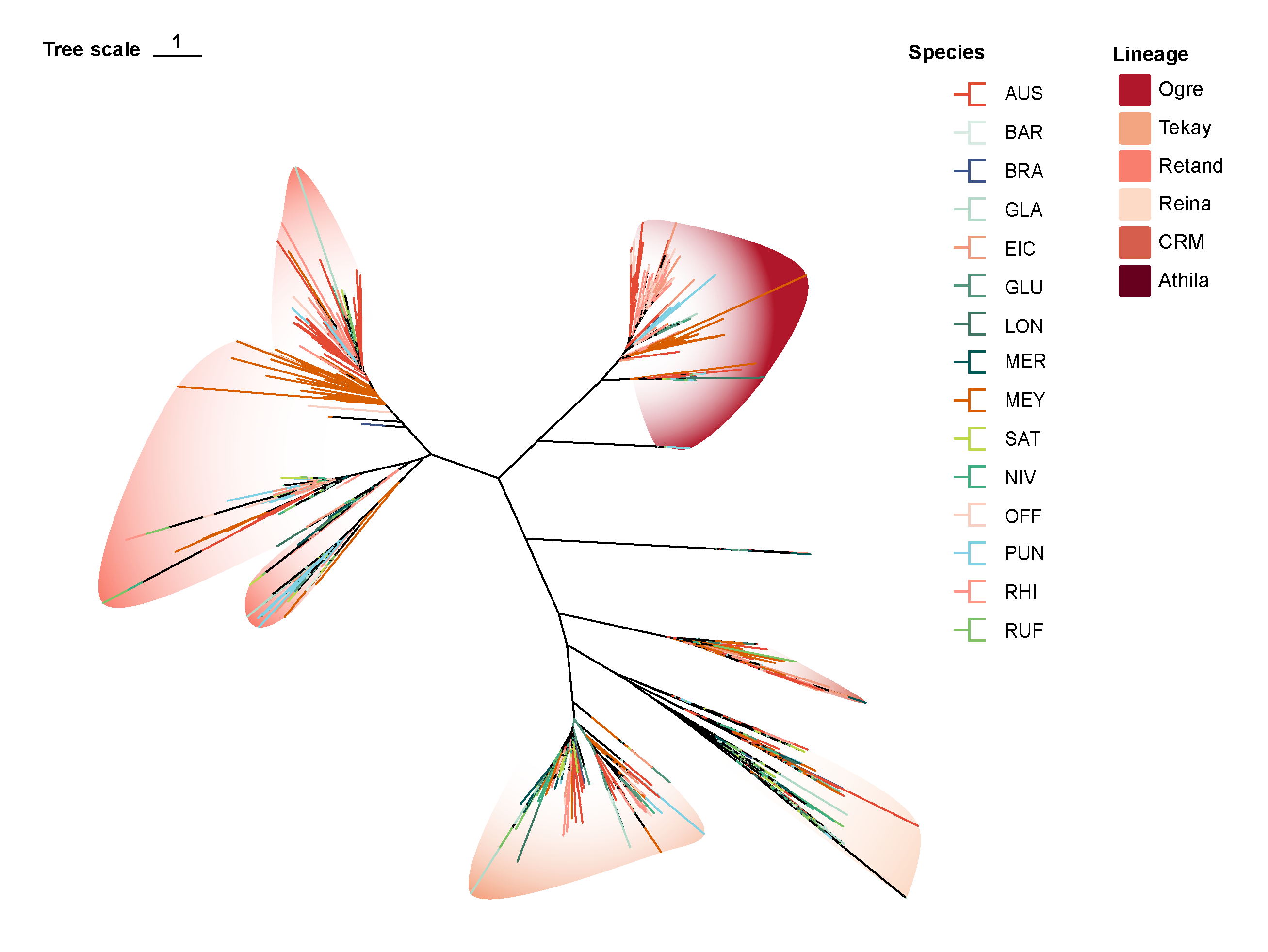

**Fig. S19.** **Unrooted gene tree of intact LTR-RTs from Ty3/Gypsy superfamily in diploid Oryza species.**

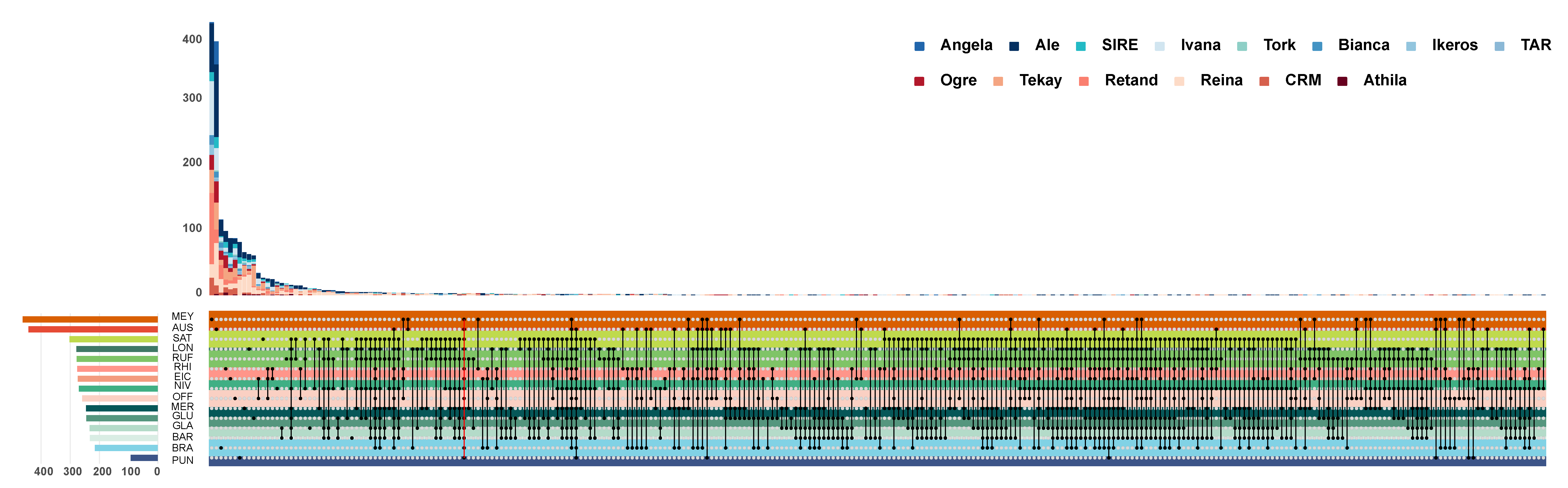

**Fig. S20.** **The upset plot of LTR-RTs families from different lineage in diploid Oryza species.**

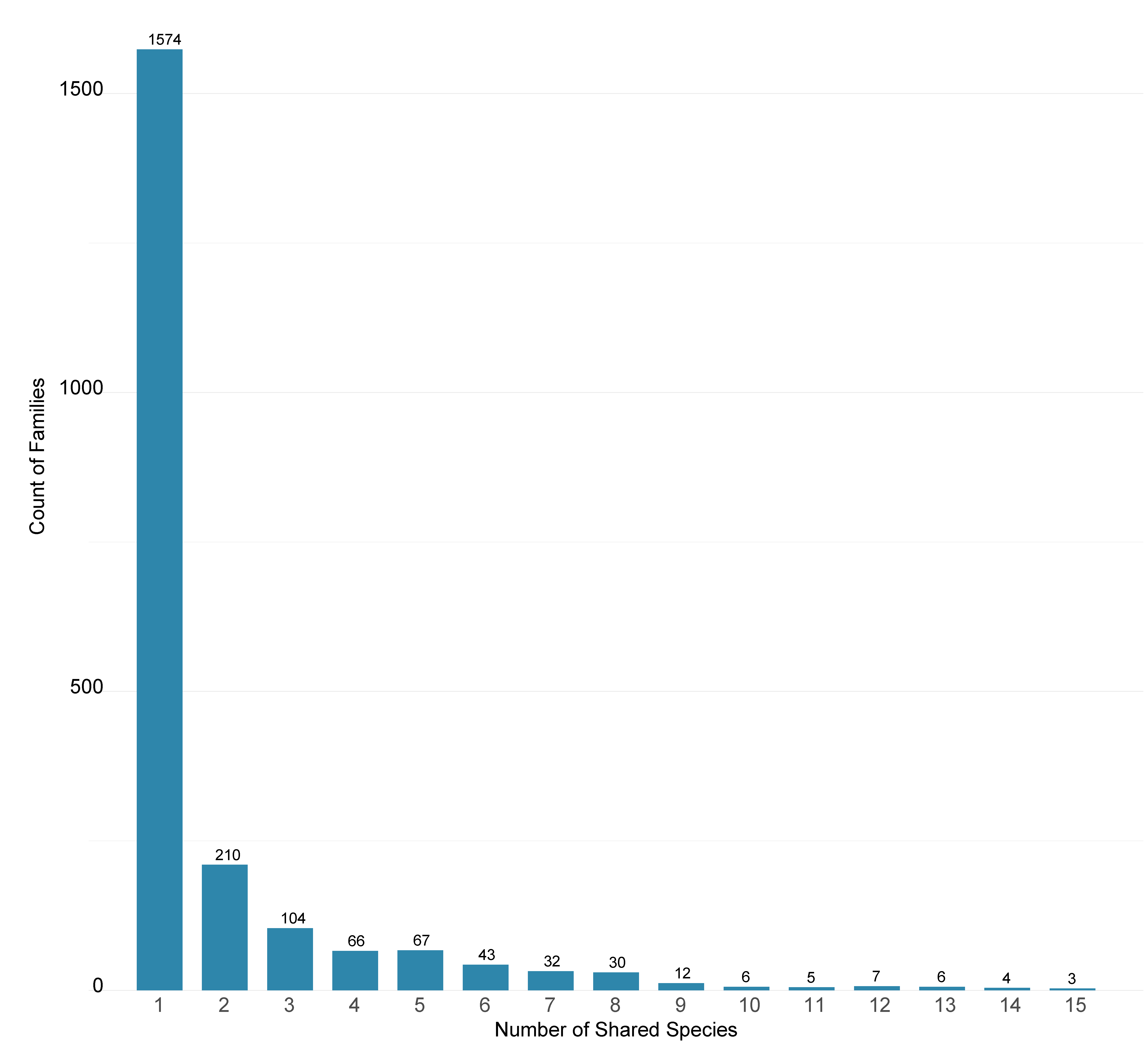

**Fig. S21.** **The Counts of species-specific and non-species-specific LTR-RTs families in Oryza diploid species.** The x-axis is the counts of species sharing these LTR-RTs families.

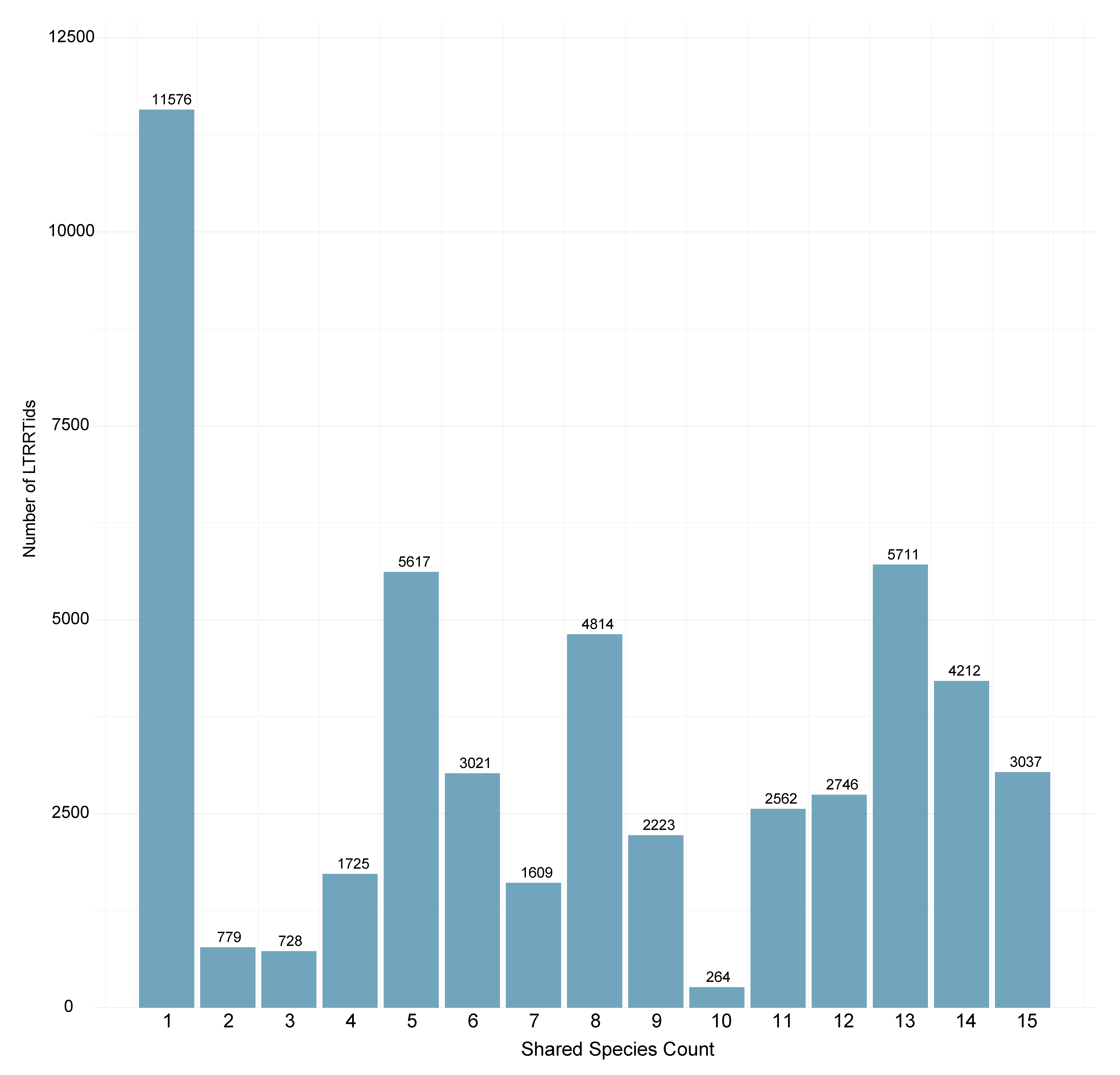

**Fig. S22.** **The Counts of the members of species-specific and non-species-specific LTR-RTs families in Oryza diploid species**. The x-axis is the counts of species sharing these LTR-RTs families.

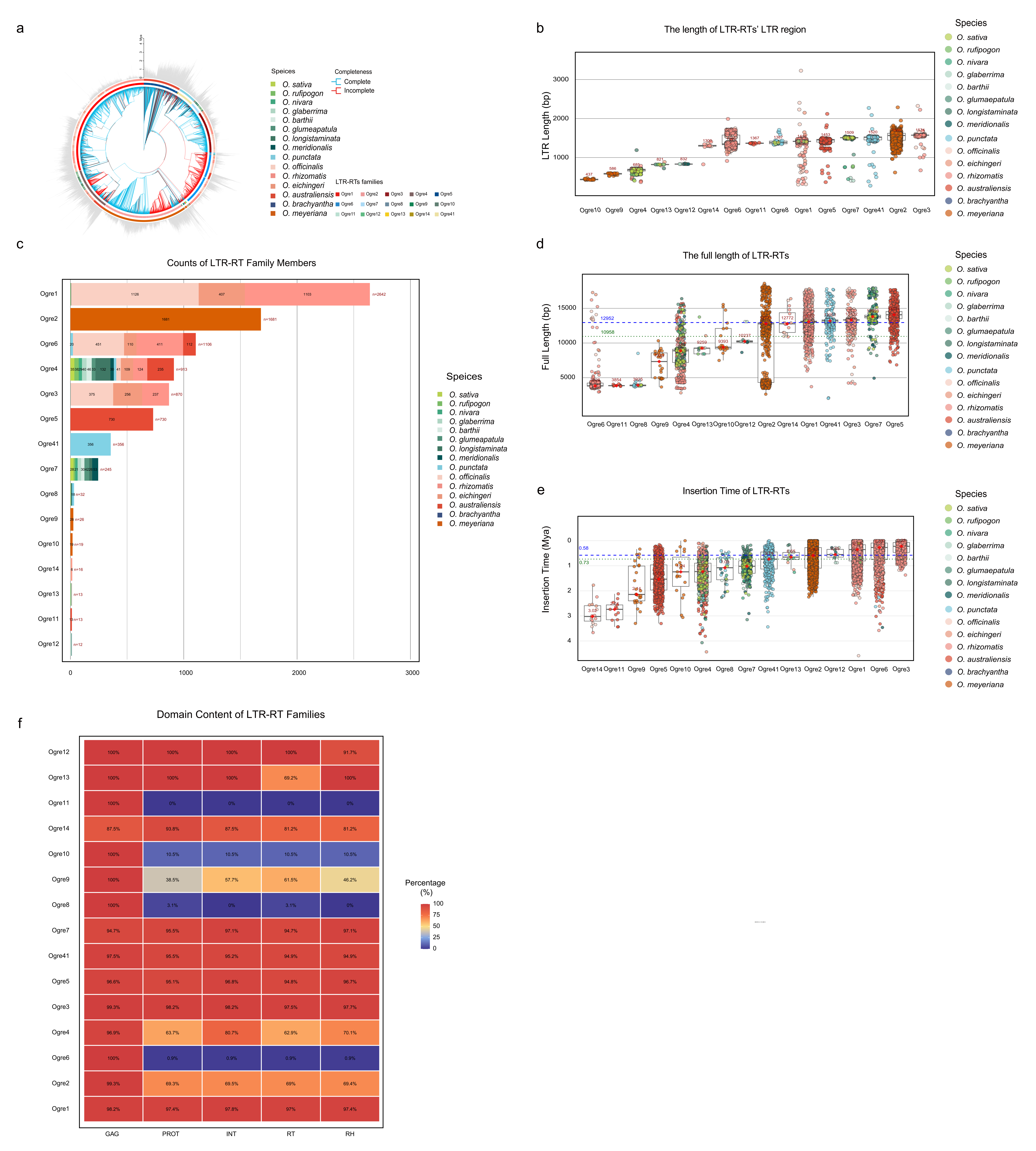

**Fig. S23. The phylogenetic tree of intact LTR-RTs and the information of main intact LTR-RTs families in Ogre lineage.** **a** is a phylogenetic tree of LTR-RTs, constructed based on the 5’ LTR region. The outermost bar plot indicates the insertion time of LTR-RTs, the middle ring layer shows the species where LTR-RTs are located, and the innermost ring represents the major families to which LTR-RTs belong. The branch colors indicate whether the domains are complete, and also distinguish whether the LTR-RTs are autonomous or non-autonomous. **b** is a boxplot showing the length of the 5’ LTR region for the major LTR-RT families. **c** shows the number and distribution of members in the major LTR-RT families. **d** is a boxplot displaying the length of members in the major LTR-RT families. **e** is a boxplot illustrating the insertion time of members in the major LTR-RT families. The green dashed line represents the mean value, while the blue dashed line indicates the median value. **f** shows the domain content of the major LTR-RT families.

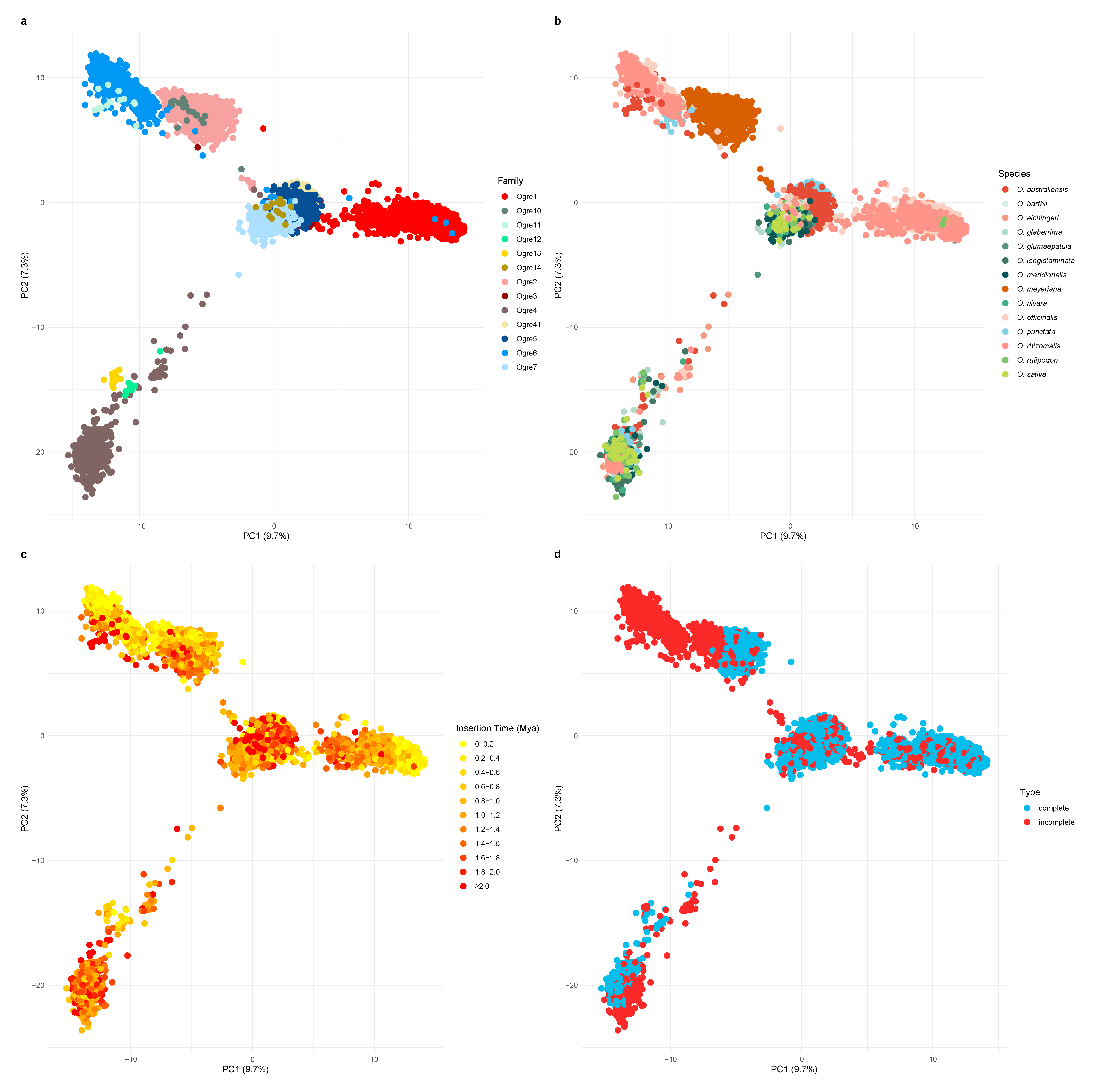

**Fig. S24. The dynamic evolution of the main LTR-RTs families in Ogre lineage, in Oryza species.** **a** indicate the PCA analysis of main intact LTR-RTs families. **b** indicates the species including these intact LTR-RTs. **c** indicates the insertion time of these intact LTR-RTs. **d** indicates the autonomous intact LTR-RTs and non-autonomous intact LTR-RTs.

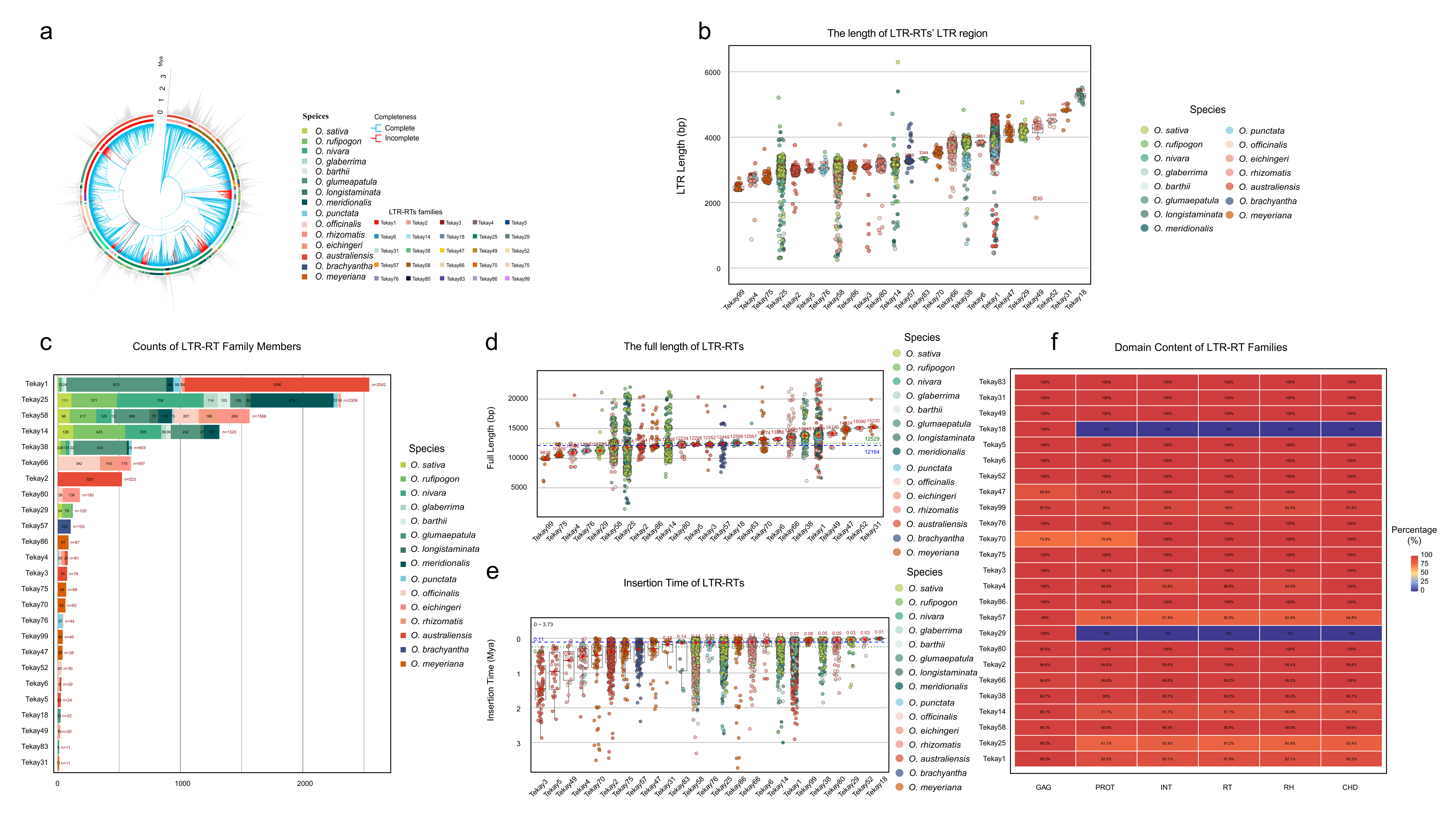

**Fig. S25. The phylogenetic tree of intact LTR-RTs and the information of main intact LTR-RTs families in Tekay lineage.** **a** is a phylogenetic tree of LTR-RTs, constructed based on the 5’ LTR region. The outermost bar plot indicates the insertion time of LTR-RTs, the middle ring layer shows the species where LTR-RTs are located, and the innermost ring represents the major families to which LTR-RTs belong. The branch colors indicate whether the domains are complete, and also distinguish whether the LTR-RTs are autonomous or non-autonomous. **b** is a boxplot showing the length of the 5’ LTR region for the major LTR-RT families. **c** shows the number and distribution of members in the major LTR-RT families. **d** is a boxplot displaying the length of members in the major LTR-RT families. **e** is a boxplot illustrating the insertion time of members in the major LTR-RT families. The green dashed line represents the mean value, while the blue dashed line indicates the median value. **f** shows the domain content of the major LTR-RT families.

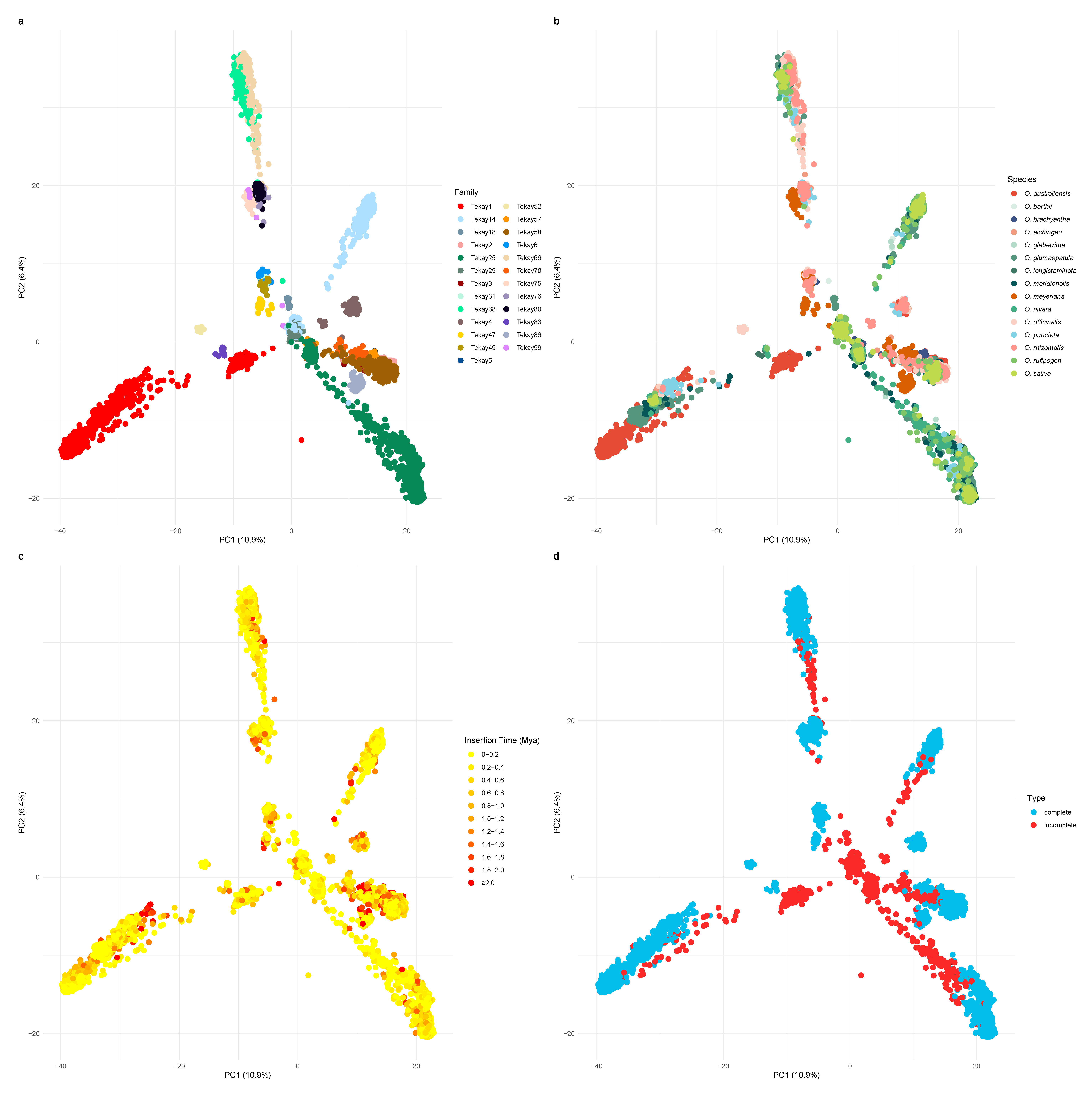

**Fig. S26. The dynamic evolution of the main LTR-RTs families in Tekay lineage, in Oryza species.** **a** indicate the PCA analysis of main intact LTR-RTs families. **b** indicates the species including these intact LTR-RTs. **c** indicates the insertion time of these intact LTR-RTs. **d** indicates the autonomous intact LTR-RTs and non-autonomous intact LTR-RTs.

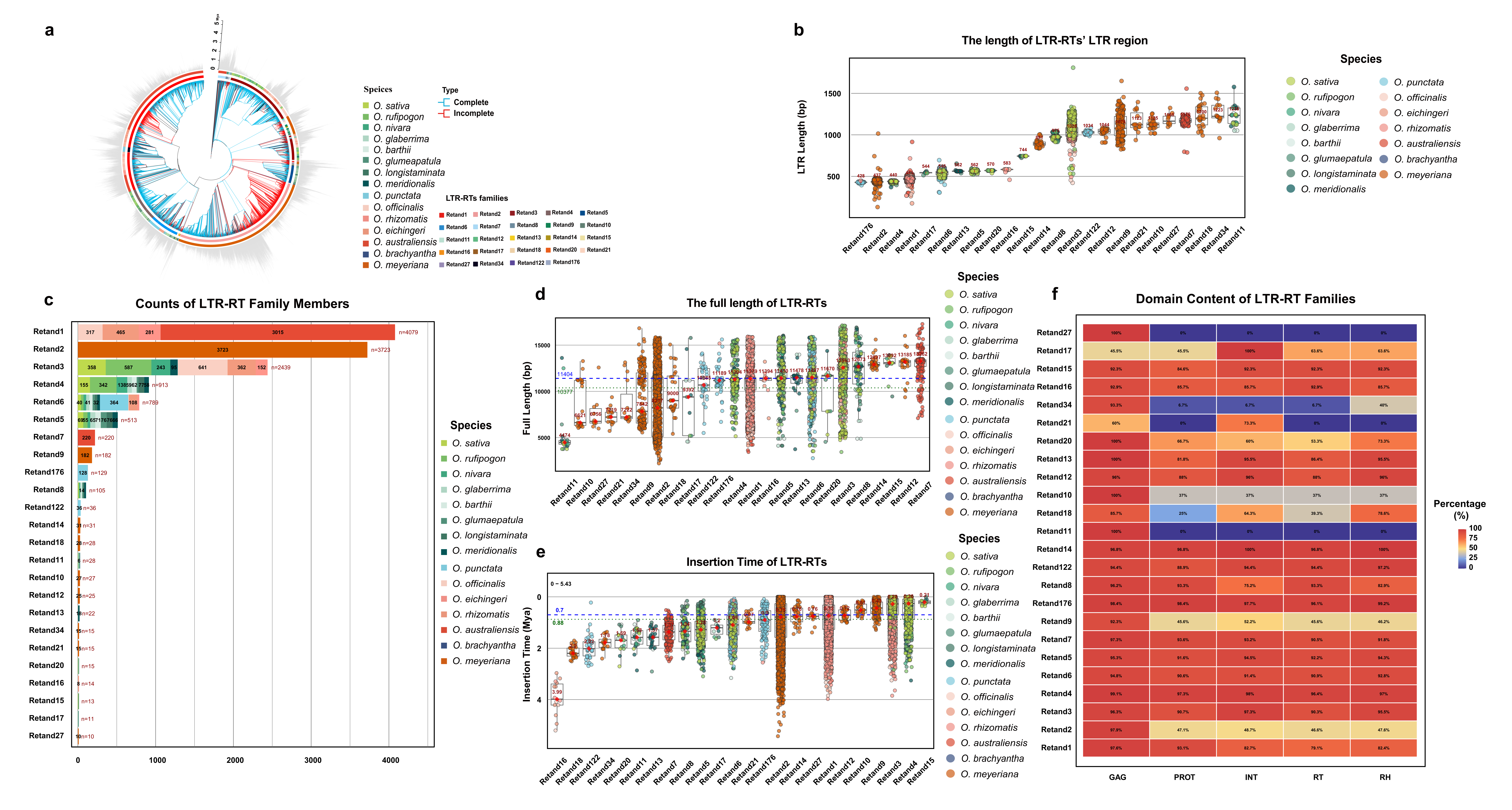

**Fig. S27. The phylogenetic tree of intact LTR-RTs and the information of main intact LTR-RTs families in Retand lineage.** **a** is a phylogenetic tree of LTR-RTs, constructed based on the 5’ LTR region. The outermost bar plot indicates the insertion time of LTR-RTs, the middle ring layer shows the species where LTR-RTs are located, and the innermost ring represents the major families to which LTR-RTs belong. The branch colors indicate whether the domains are complete, and also distinguish whether the LTR-RTs are autonomous or non-autonomous. **b** is a boxplot showing the length of the 5’ LTR region for the major LTR-RT families. **c** shows the number and distribution of members in the major LTR-RT families. **d** is a boxplot displaying the length of members in the major LTR-RT families. **e** is a boxplot illustrating the insertion time of members in the major LTR-RT families. The green dashed line represents the mean value, while the blue dashed line indicates the median value. **f** shows the domain content of the major LTR-RT families.

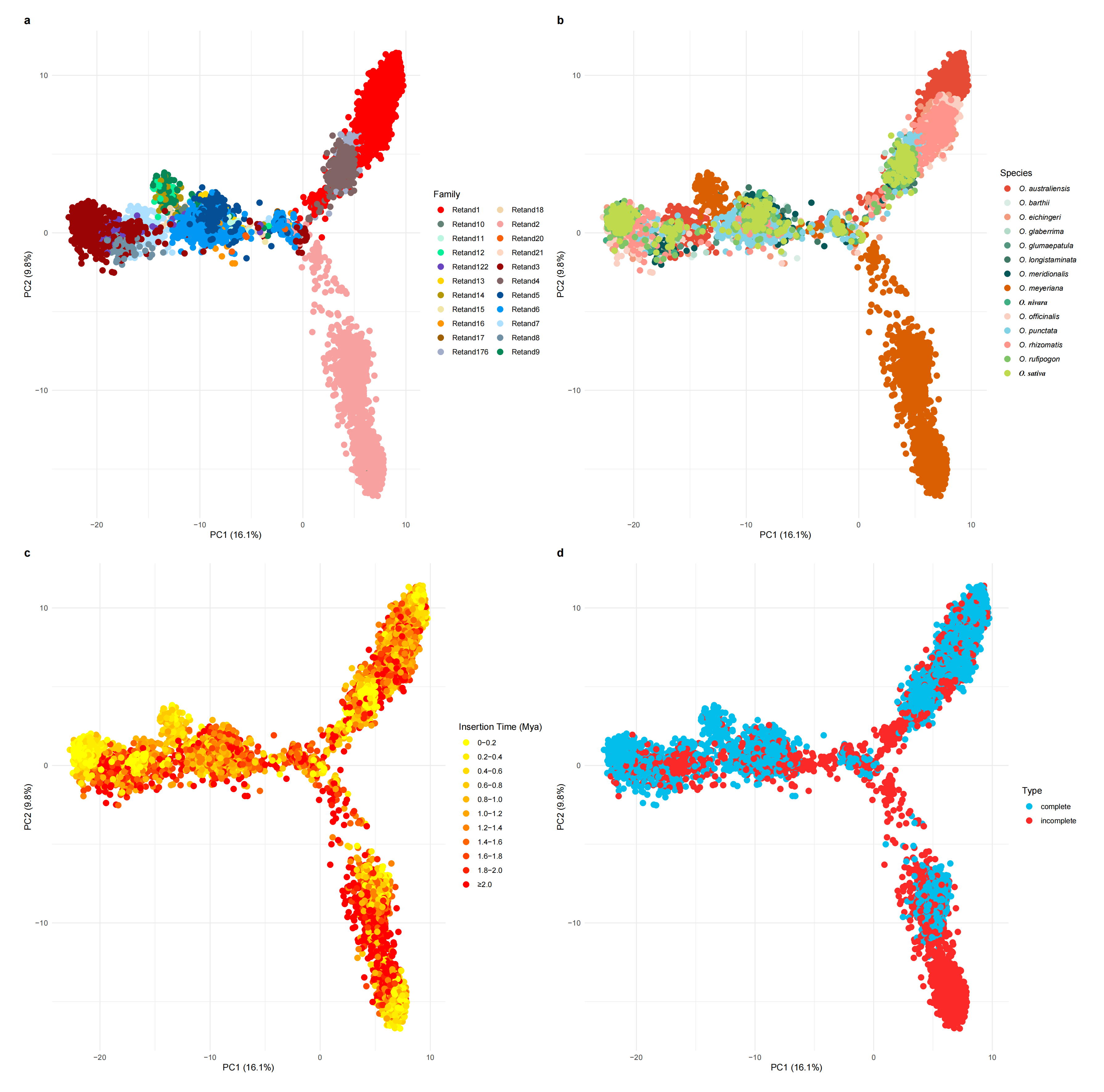

**Fig. S28. The dynamic evolution of the main LTR-RTs families in Retand lineage, in Oryza species.** **a** indicates the PCA analysis of main intact LTR-RTs families. **b** indicates the species including these intact LTR-RTs. **c** indicates the insertion time of these intact LTR-RTs. **d** indicates the autonomous intact LTR-RTs and non-autonomous intact LTR-RTs.

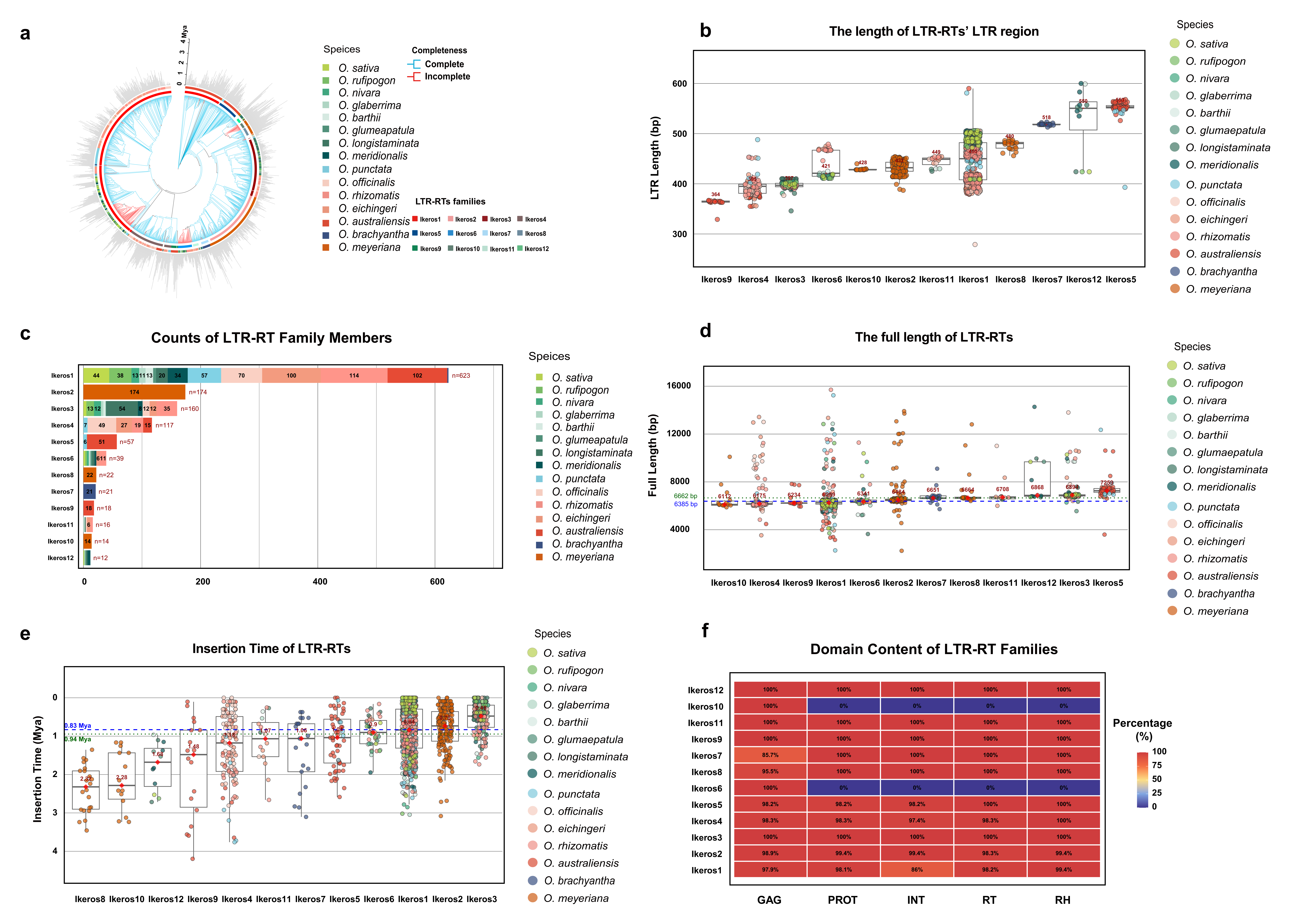

**Fig. S29. The phylogenetic tree of intact LTR-RTs and the information of main intact LTR-RTs families in Ikeros lineage.** **a** is a phylogenetic tree of LTR-RTs, constructed based on the 5’ LTR region. The outermost bar plot indicates the insertion time of LTR-RTs, the middle ring layer shows the species where LTR-RTs are located, and the innermost ring represents the major families to which LTR-RTs belong. The branch colors indicate whether the domains are complete, and also distinguish whether the LTR-RTs are autonomous or non-autonomous. **b** is a boxplot showing the length of the 5’ LTR region for the major LTR-RT families. **c** shows the number and distribution of members in the major LTR-RT families. **d** is a boxplot displaying the length of members in the major LTR-RT families. **e** is a boxplot illustrating the insertion time of members in the major LTR-RT families. The green dashed line represents the mean value, while the blue dashed line indicates the median value. **f** shows the domain content of the major LTR-RT families.

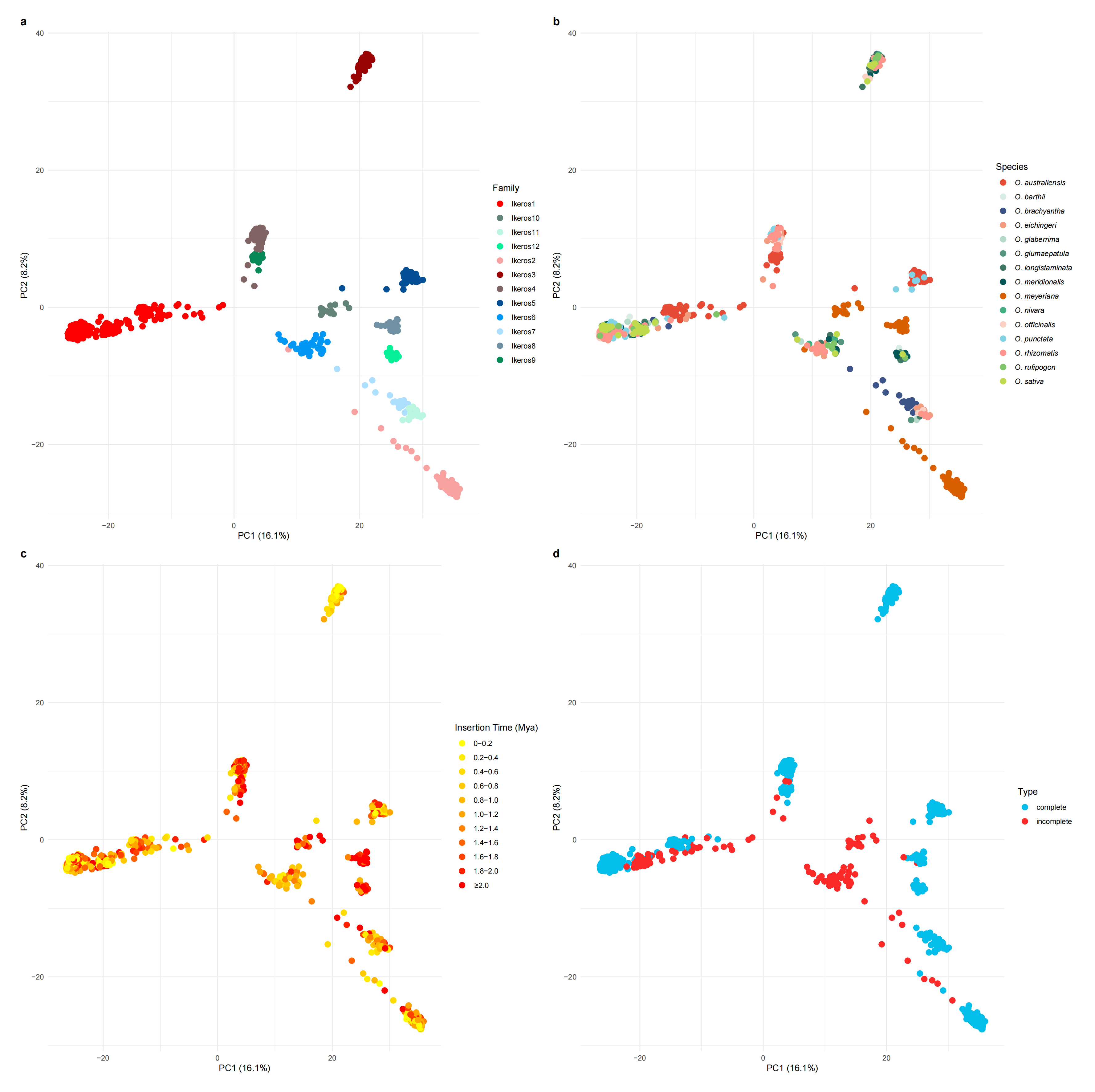

**Fig. S30. The dynamic evolution of the main LTR-RTs families in Ikeros lineage, in Oryza species.** **a** indicates the PCA analysis of main intact LTR-RTs families. **b** indicates the species including these intact LTR-RTs. **c** indicates the insertion time of these intact LTR-RTs. **d** indicates the autonomous intact LTR-RTs and non-autonomous intact LTR-RTs.

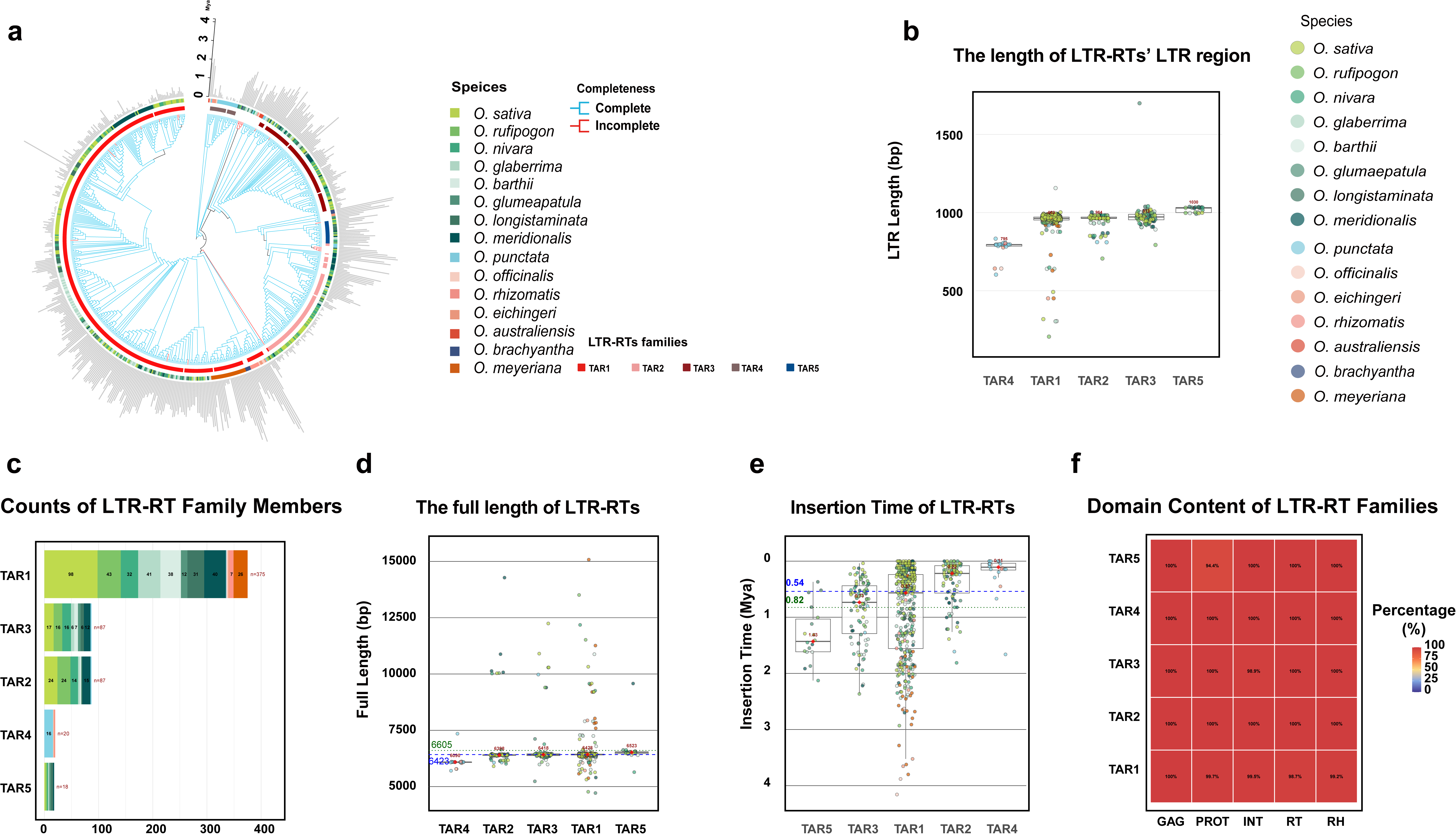

**Fig. S31. The phylogenetic tree of intact LTR-RTs and the information of main intact LTR-RTs families in TAR lineage.** **a** is a phylogenetic tree of LTR-RTs, constructed based on the 5’ LTR region. The outermost bar plot indicates the insertion time of LTR-RTs, the middle ring layer shows the species where LTR-RTs are located, and the innermost ring represents the major families to which LTR-RTs belong. The branch colors indicate whether the domains are complete, and also distinguish whether the LTR-RTs are autonomous or non-autonomous. **b** is a boxplot showing the length of the 5’ LTR region for the major LTR-RT families. **c** shows the number and distribution of members in the major LTR-RT families. **d** is a boxplot displaying the length of members in the major LTR-RT families. **e** is a boxplot illustrating the insertion time of members in the major LTR-RT families. The green dashed line represents the mean value, while the blue dashed line indicates the median value. **f** shows the domain content of the major LTR-RT families.

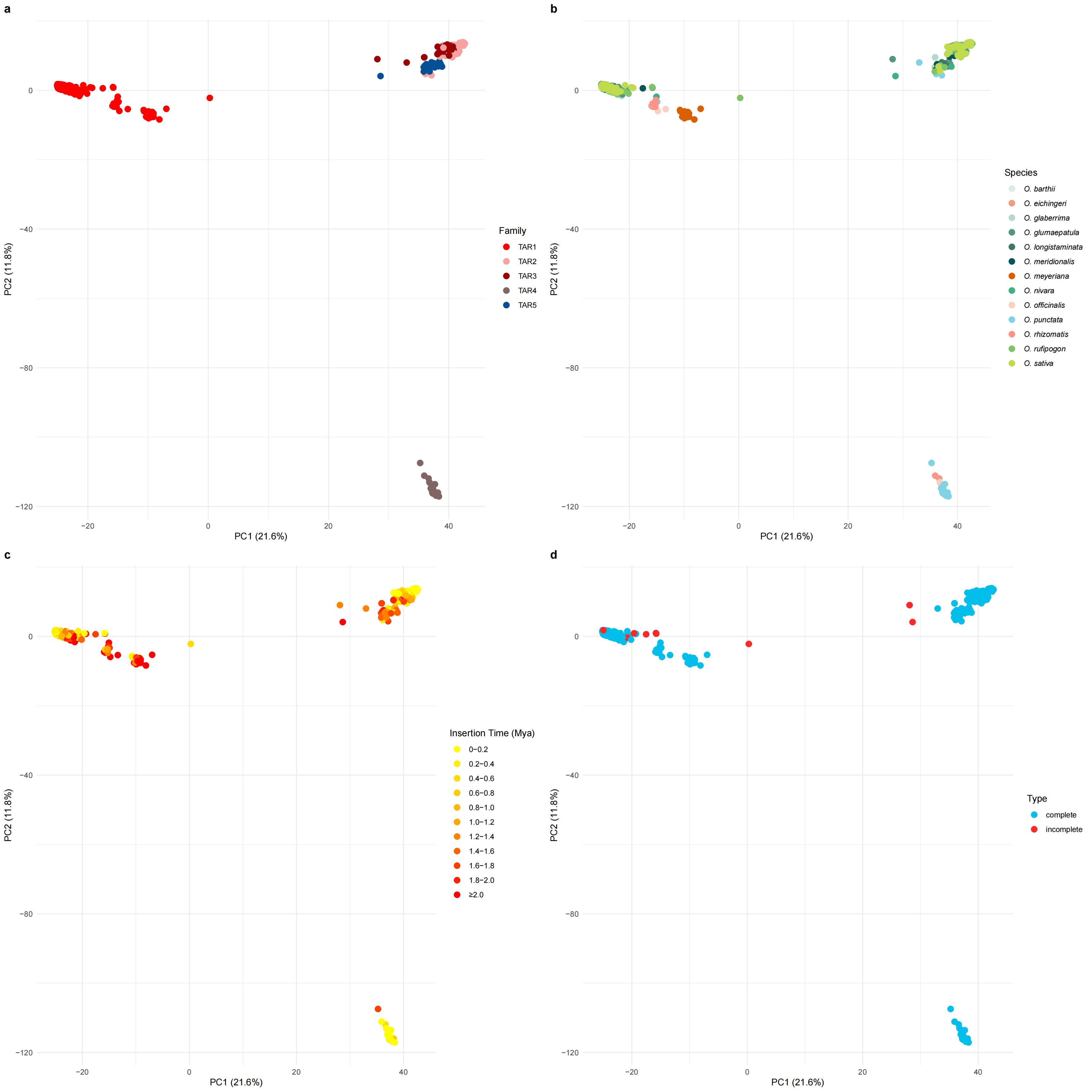

**Fig. S32. The dynamic evolution of the main LTR-RTs families in TAR lineage, in Oryza species.** **a** indicate the PCA analysis of main intact LTR-RTs families. **b** indicates the species including these intact LTR-RTs. **c** indicates the insertion time of these intact LTR-RTs. **d** indicates the autonomous intact LTR-RTs and non-autonomous intact LTR-RTs.

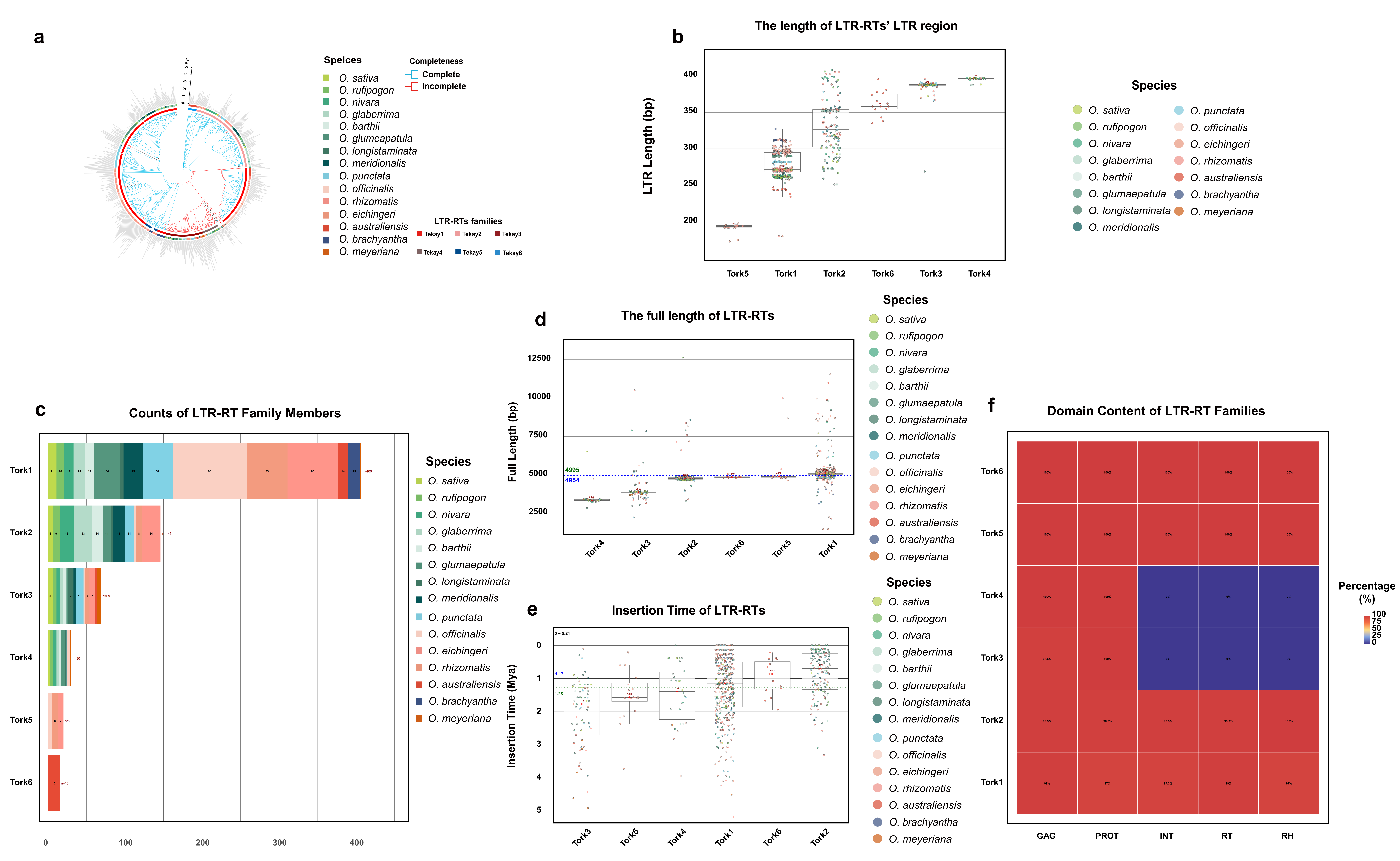

**Fig. S33. The phylogenetic tree of intact LTR-RTs and the information of main intact LTR-RTs families in Tork lineage.** **a** is a phylogenetic tree of LTR-RTs, constructed based on the 5’ LTR region. The outermost bar plot indicates the insertion time of LTR-RTs, the middle ring layer shows the species where LTR-RTs are located, and the innermost ring represents the major families to which LTR-RTs belong. The branch colors indicate whether the domains are complete, and also distinguish whether the LTR-RTs are autonomous or non-autonomous. **b** is a boxplot showing the length of the 5’ LTR region for the major LTR-RT families. **c** shows the number and distribution of members in the major LTR-RT families. **d** is a boxplot displaying the length of members in the major LTR-RT families. **e** is a boxplot illustrating the insertion time of members in the major LTR-RT families. The green dashed line represents the mean value, while the blue dashed line indicates the median value. **f** shows the domain content of the major LTR-RT families.

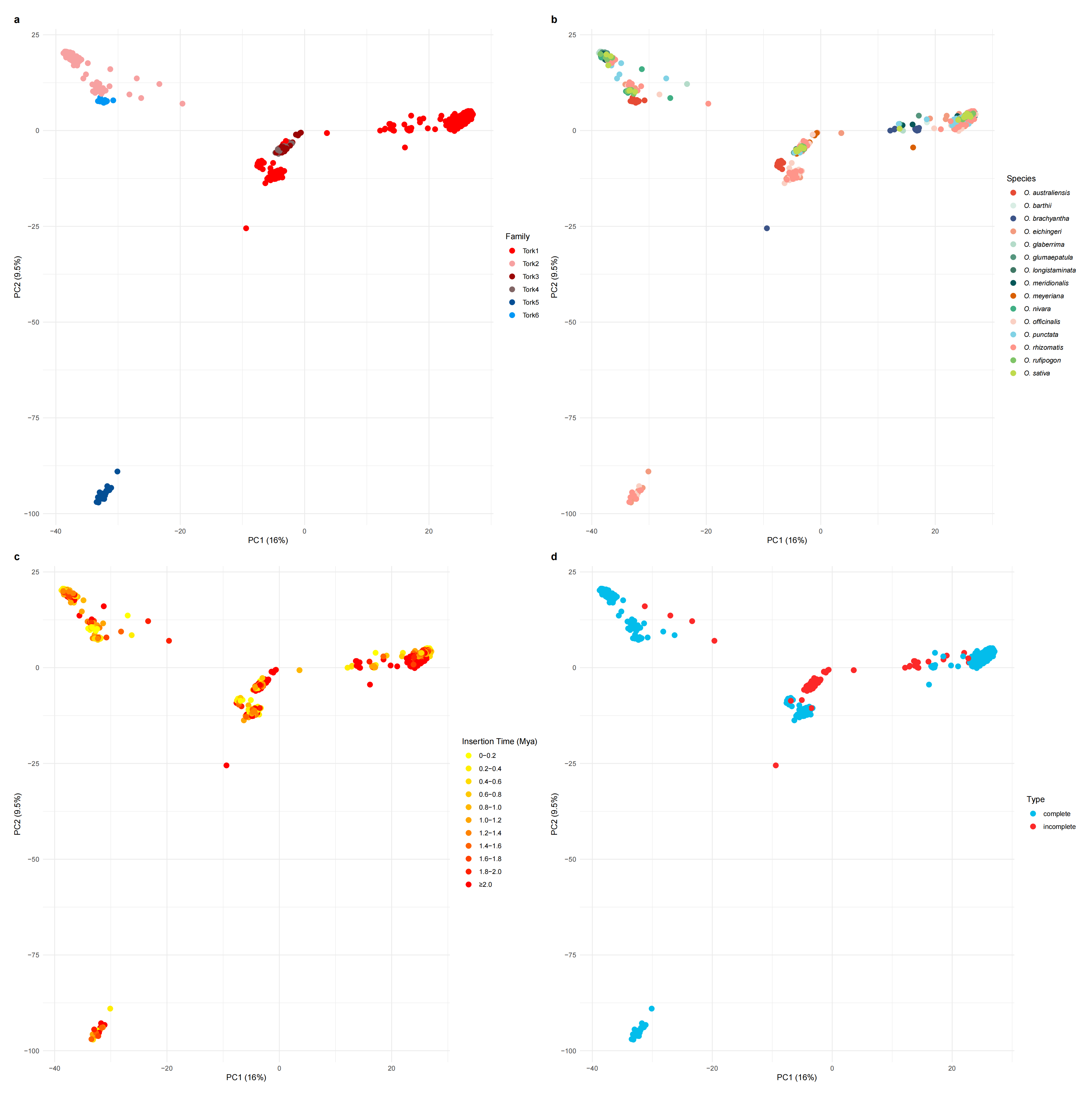

**Fig. S34. The dynamic evolution of the main LTR-RTs families in Tork lineage, in Oryza species.** **a** indicates the PCA analysis of main intact LTR-RTs families. **b** indicates the species including these intact LTR-RTs. **c** indicates the insertion time of these intact LTR-RTs. **d** indicates the autonomous intact LTR-RTs and non-autonomous intact LTR-RTs.

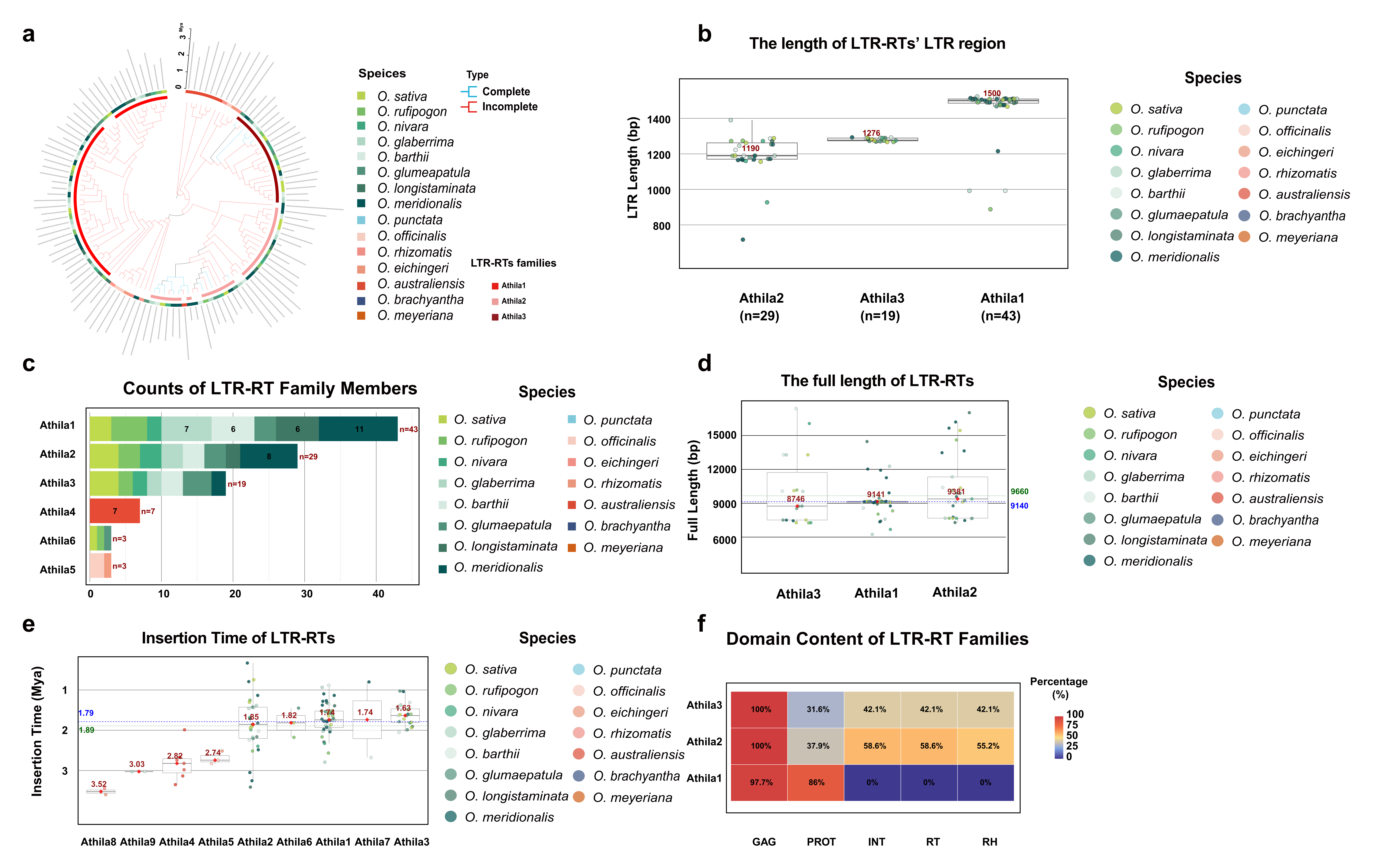

**Fig. S35. The phylogenetic tree of intact LTR-RTs and the information of main intact LTR-RTs families in Athila lineage.** **a** is a phylogenetic tree of LTR-RTs, constructed based on the 5’ LTR region. The outermost bar plot indicates the insertion time of LTR-RTs, the middle ring layer shows the species where LTR-RTs are located, and the innermost ring represents the major families to which LTR-RTs belong. The branch colors indicate whether the domains are complete, and also distinguish whether the LTR-RTs are autonomous or non-autonomous. **b** is a boxplot showing the length of the 5’ LTR region for the major LTR-RT families. **c** shows the number and distribution of members in the major LTR-RT families. **d** is a boxplot displaying the length of members in the major LTR-RT families. **e** is a boxplot illustrating the insertion time of members in the major LTR-RT families. The green dashed line represents the mean value, while the blue dashed line indicates the median value. **f** shows the domain content of the major LTR-RT families.

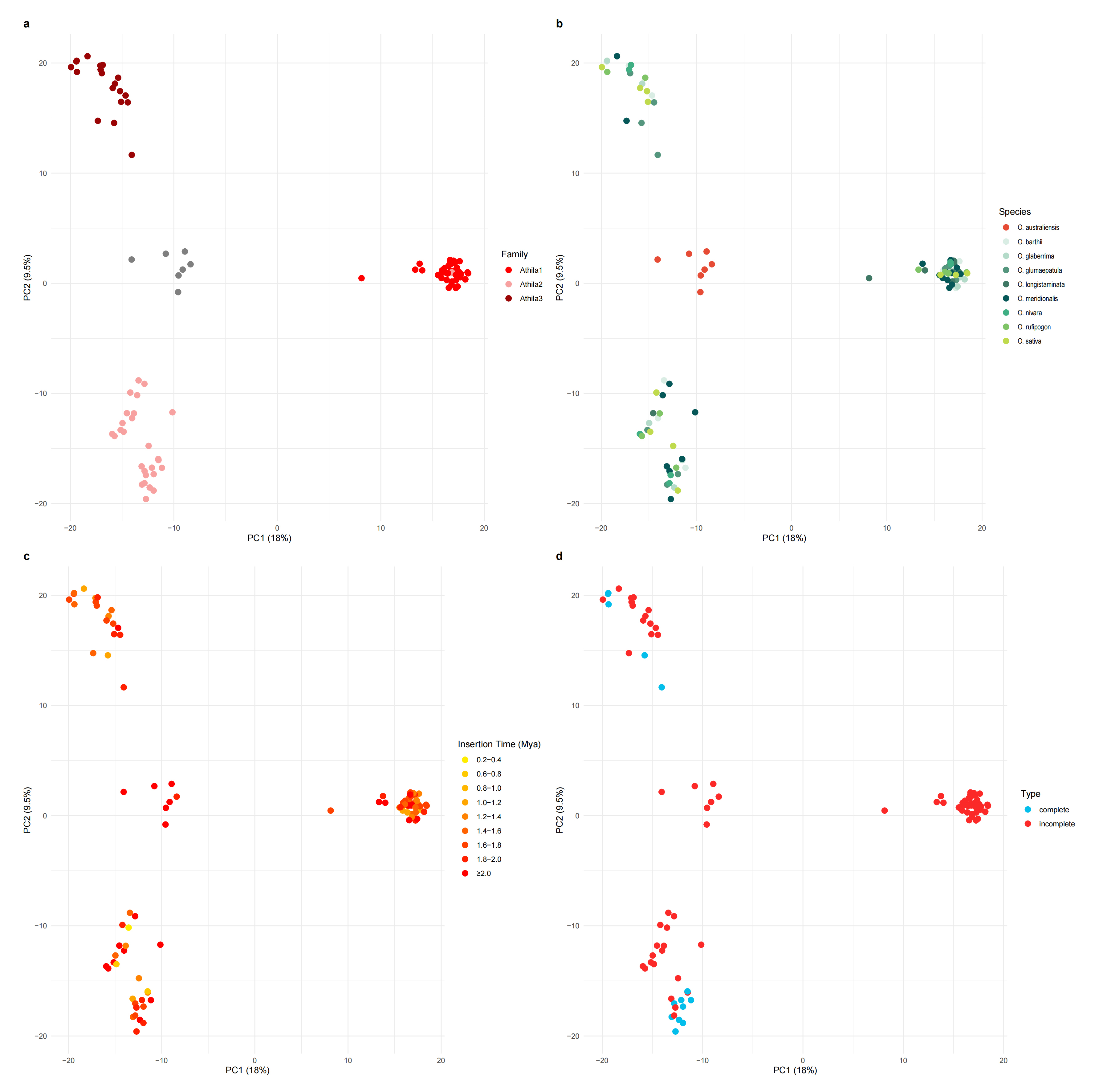

**Fig. S36. The dynamic evolution of the main LTR-RTs families in Athila lineage, in Oryza species.** **a** indicates the PCA analysis of main intact LTR-RTs families. **b** indicates the species including these intact LTR-RTs. **c** indicates the insertion time of these intact LTR-RTs. **d** indicates the autonomous intact LTR-RTs and non-autonomous intact LTR-RTs.

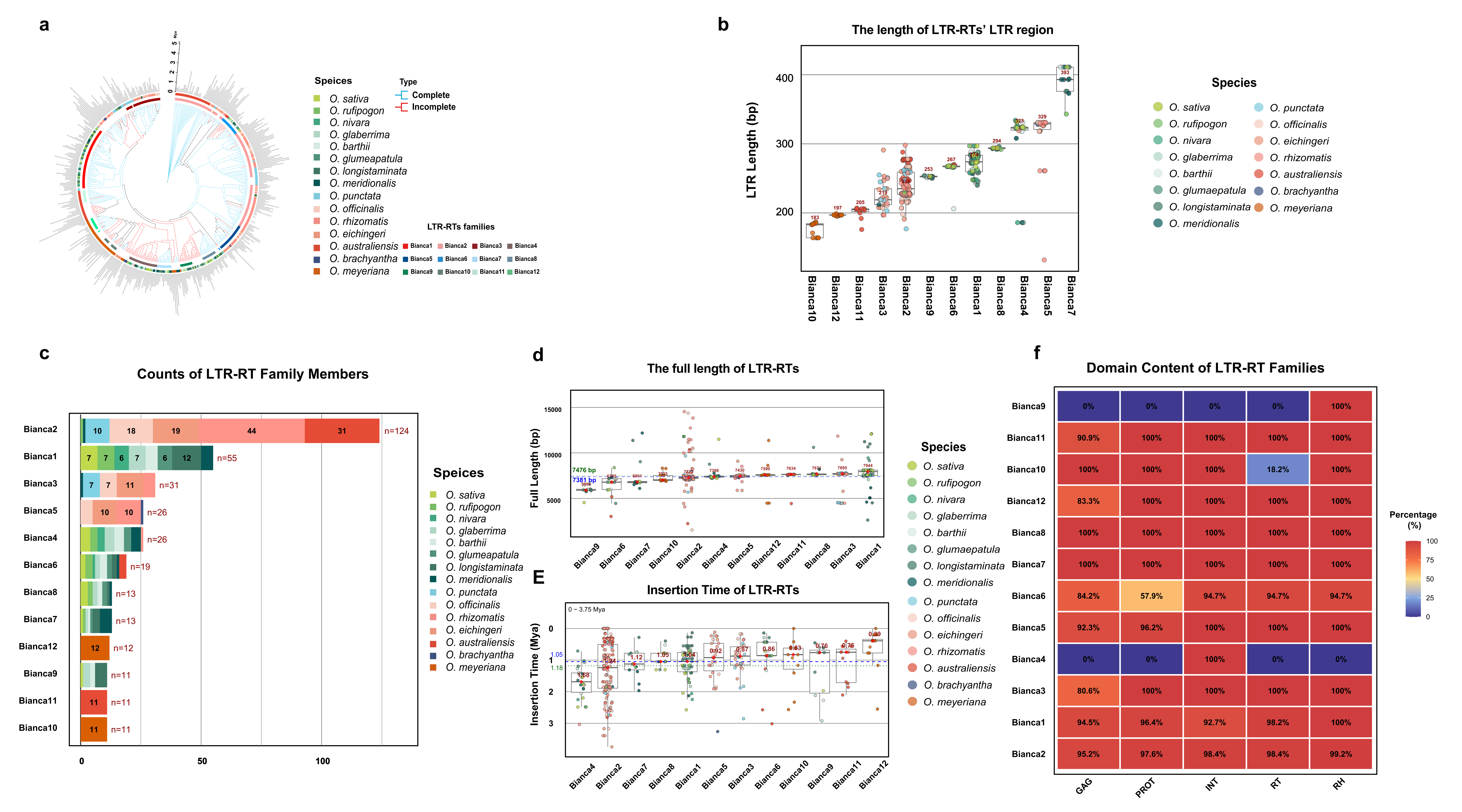

**Fig. S37. The phylogenetic tree of intact LTR-RTs and the information of main intact LTR-RTs families in Bianca lineage.** **a** is a phylogenetic tree of LTR-RTs, constructed based on the 5’ LTR region. The outermost bar plot indicates the insertion time of LTR-RTs, the middle ring layer shows the species where LTR-RTs are located, and the innermost ring represents the major families to which LTR-RTs belong. The branch colors indicate whether the domains are complete, and also distinguish whether the LTR-RTs are autonomous or non-autonomous. **b** is a boxplot showing the length of the 5’ LTR region for the major LTR-RT families. **c** shows the number and distribution of members in the major LTR-RT families. **d** is a boxplot displaying the length of members in the major LTR-RT families. **e** is a boxplot illustrating the insertion time of members in the major LTR-RT families. The green dashed line represents the mean value, while the blue dashed line indicates the median value. **f** shows the domain content of the major LTR-RT families.

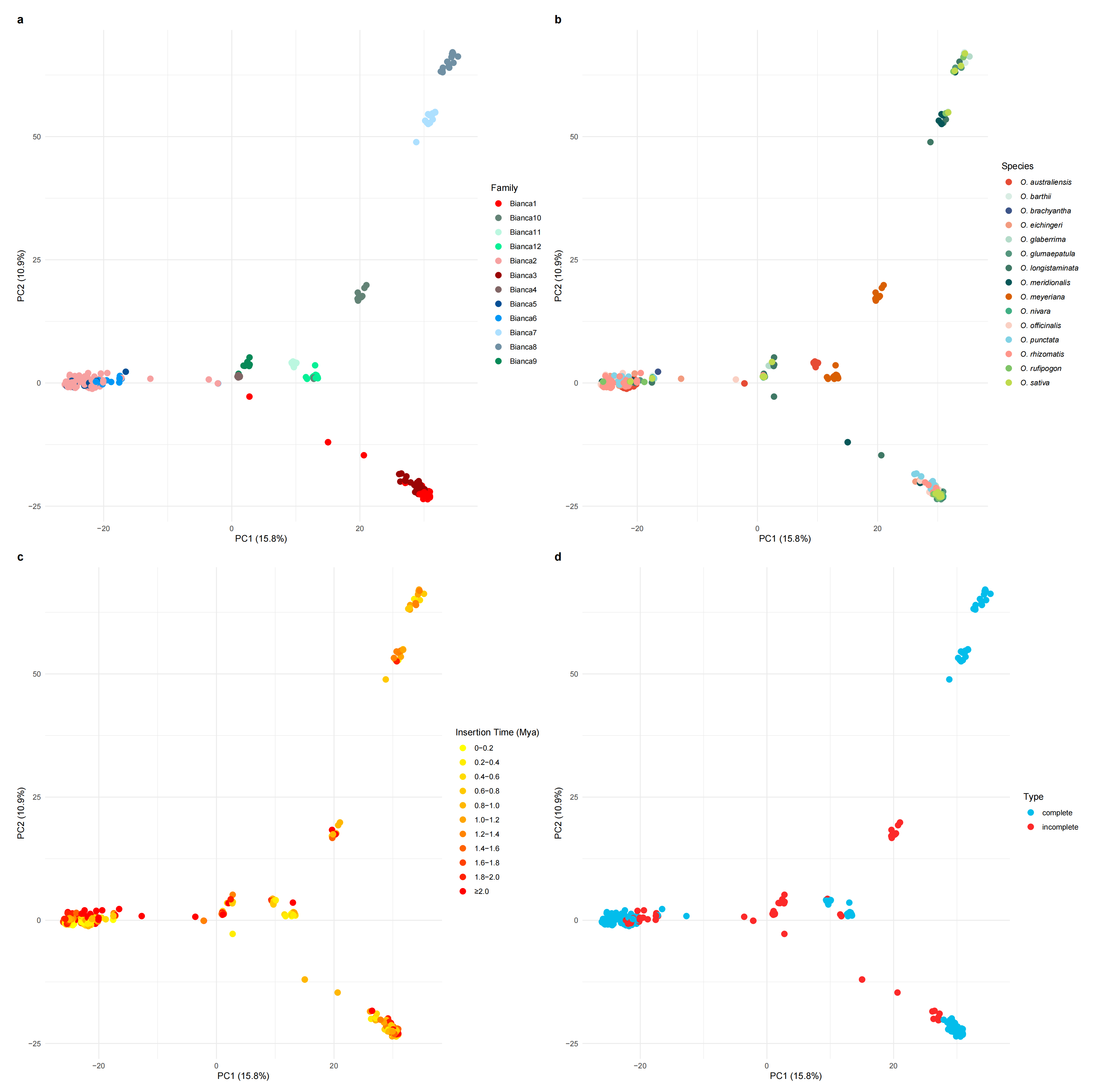

**Fig. S38. The dynamic evolution of the main LTR-RTs families in Bianca lineage, in Oryza species.** **a** indicates the PCA analysis of main intact LTR-RTs families. **b** indicates the species including these intact LTR-RTs. **c** indicates the insertion time of these intact LTR-RTs. **d** indicates the autonomous intact LTR-RTs and non-autonomous intact LTR-RTs.

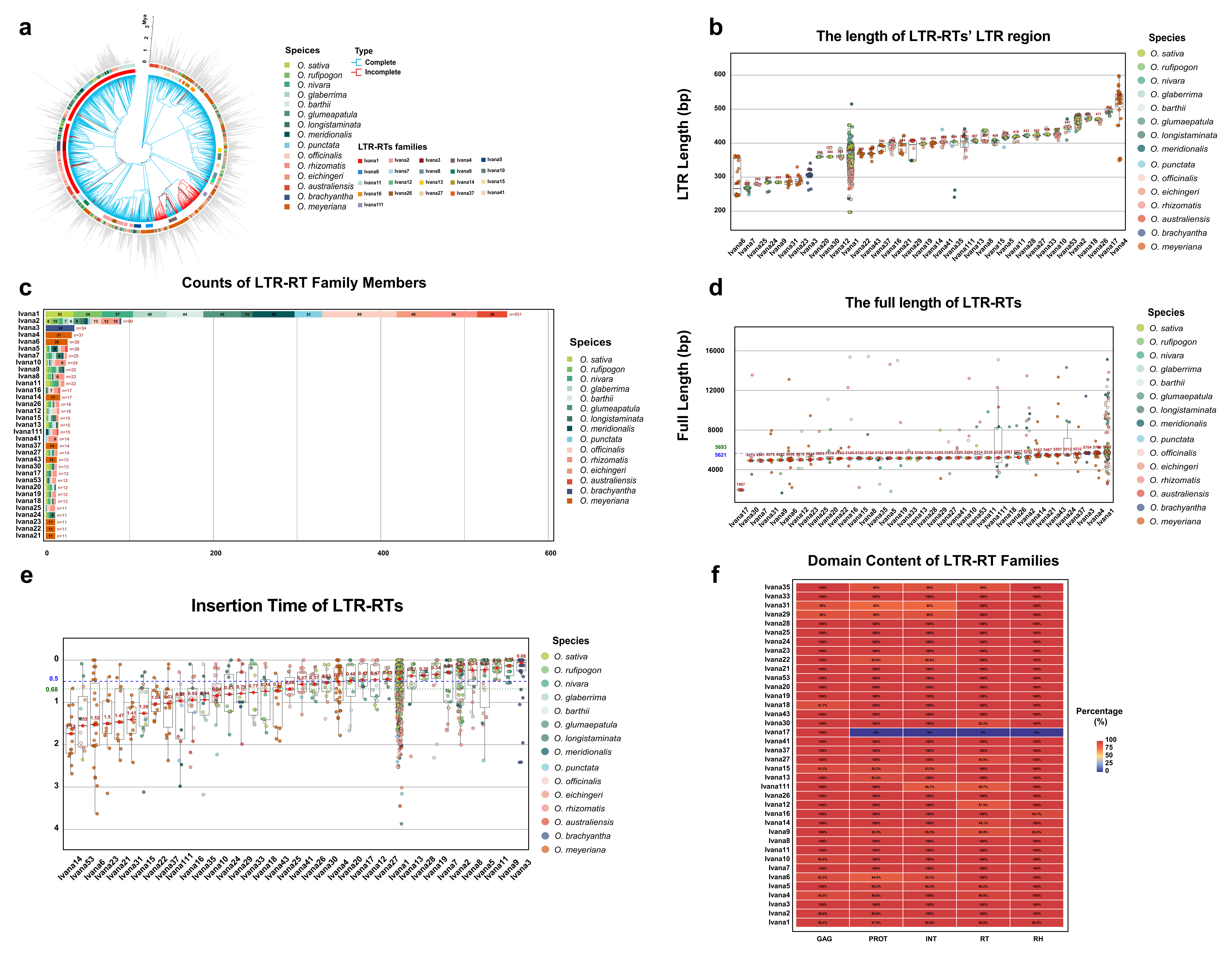

**Fig. S39. The phylogenetic tree of intact LTR-RTs and the information of main intact LTR-RTs families in Ivana lineage.** **a** is a phylogenetic tree of LTR-RTs, constructed based on the 5’ LTR region. The outermost bar plot indicates the insertion time of LTR-RTs, the middle ring layer shows the species where LTR-RTs are located, and the innermost ring represents the major families to which LTR-RTs belong. The branch colors indicate whether the domains are complete, and also distinguish whether the LTR-RTs are autonomous or non-autonomous. **b** is a boxplot showing the length of the 5’ LTR region for the major LTR-RT families. **c** shows the number and distribution of members in the major LTR-RT families. **d** is a boxplot displaying the length of members in the major LTR-RT families. **e** is a boxplot illustrating the insertion time of members in the major LTR-RT families. The green dashed line represents the mean value, while the blue dashed line indicates the median value. **f** shows the domain content of the major LTR-RT families.

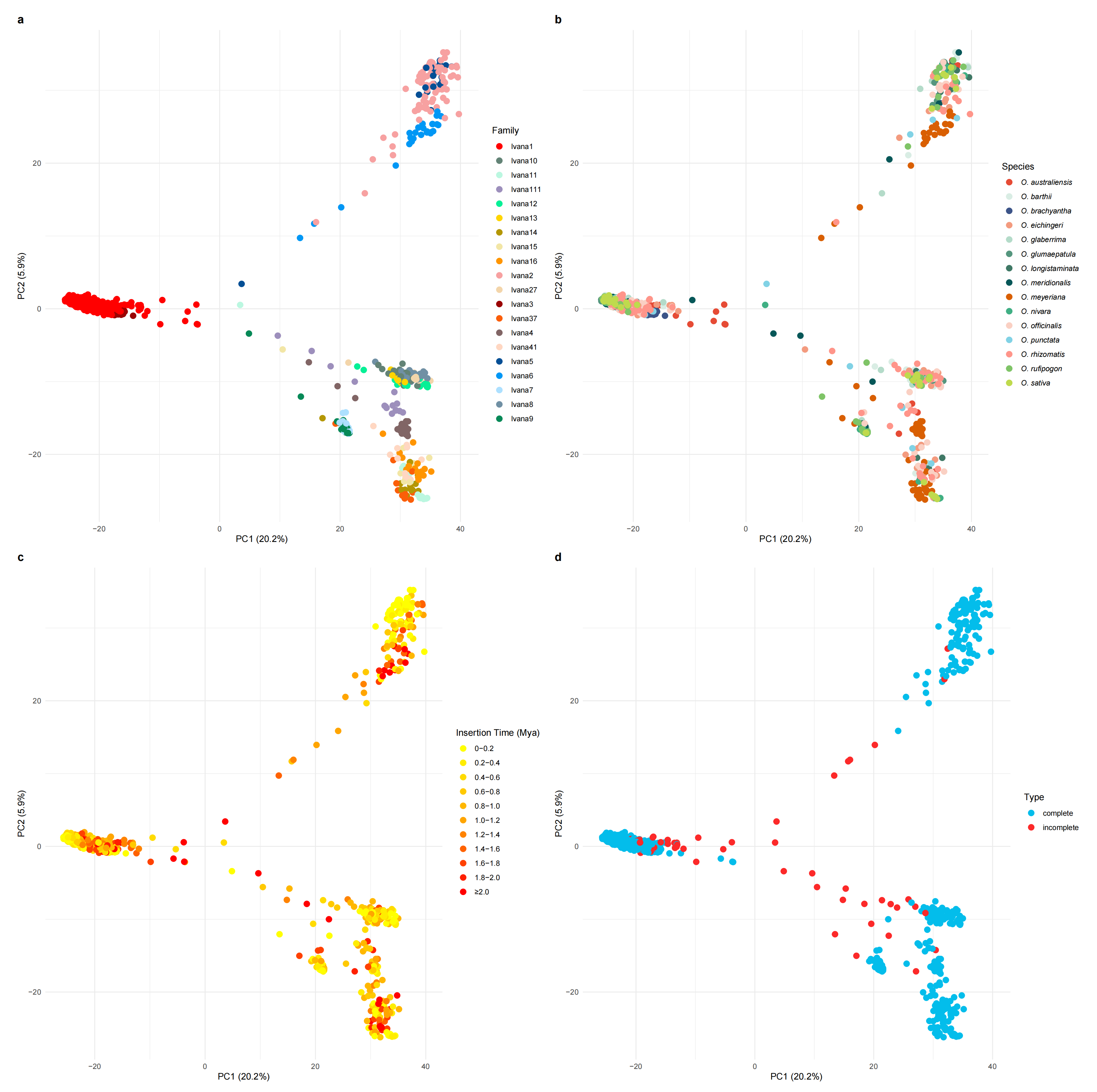

**Fig. S40. The dynamic evolution of the main LTR-RTs families in Ivana lineage, in Oryza species.** **a** indicates the PCA analysis of main intact LTR-RTs families. **b** indicates the species including these intact LTR-RTs. **c** indicates the insertion time of these intact LTR-RTs. **d** indicates the autonomous intact LTR-RTs and non-autonomous intact LTR-RTs.

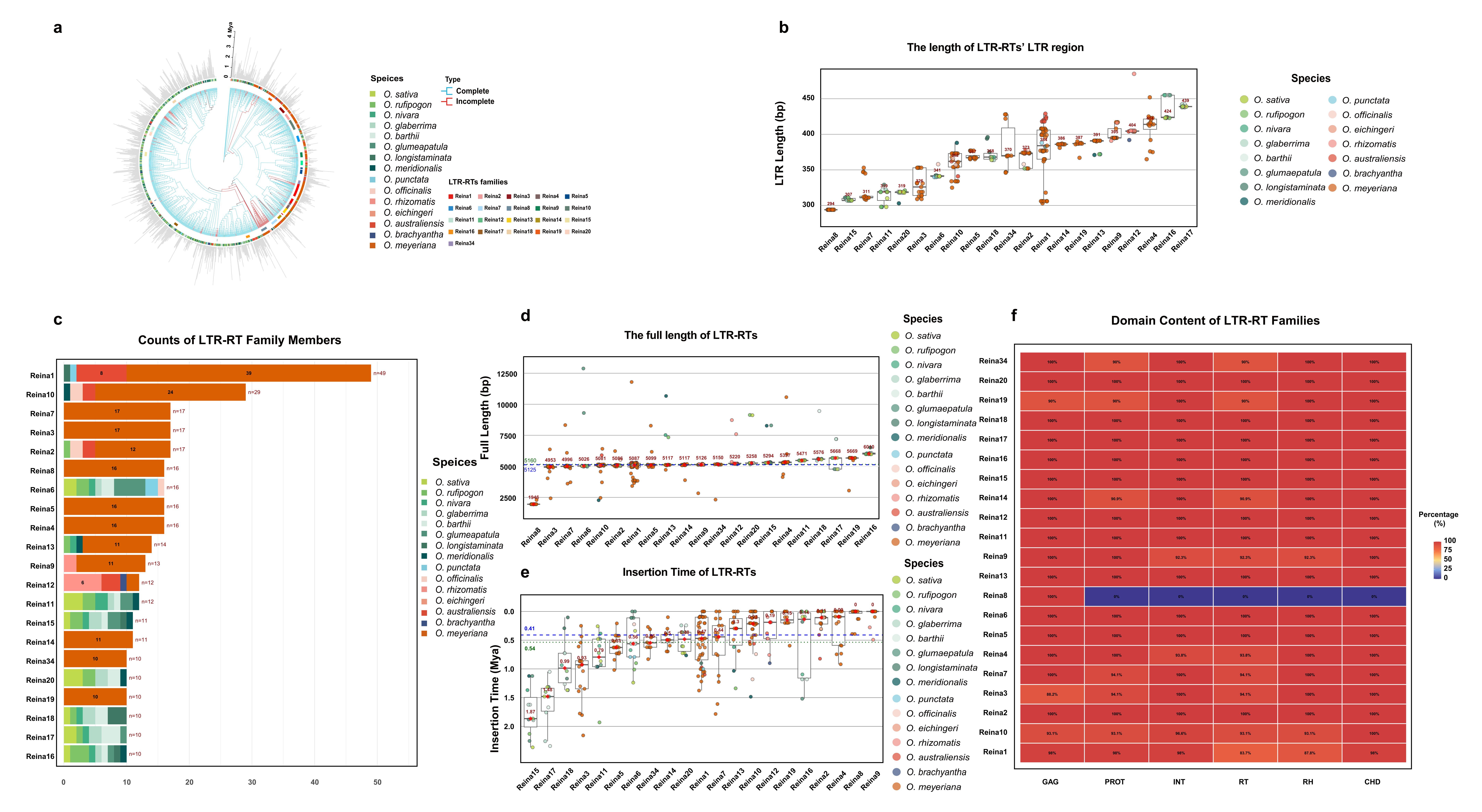

**Fig. S41. The phylogenetic tree of intact LTR-RTs and the information of main intact LTR-RTs families in Reina lineage.** **a** is a phylogenetic tree of LTR-RTs, constructed based on the 5’ LTR region. The outermost bar plot indicates the insertion time of LTR-RTs, the middle ring layer shows the species where LTR-RTs are located, and the innermost ring represents the major families to which LTR-RTs belong. The branch colors indicate whether the domains are complete, and also distinguish whether the LTR-RTs are autonomous or non-autonomous. **b** is a boxplot showing the length of the 5’ LTR region for the major LTR-RT families. **c** shows the number and distribution of members in the major LTR-RT families. **d** is a boxplot displaying the length of members in the major LTR-RT families. **e** is a boxplot illustrating the insertion time of members in the major LTR-RT families. The green dashed line represents the mean value, while the blue dashed line indicates the median value. **f** shows the domain content of the major LTR-RT families.

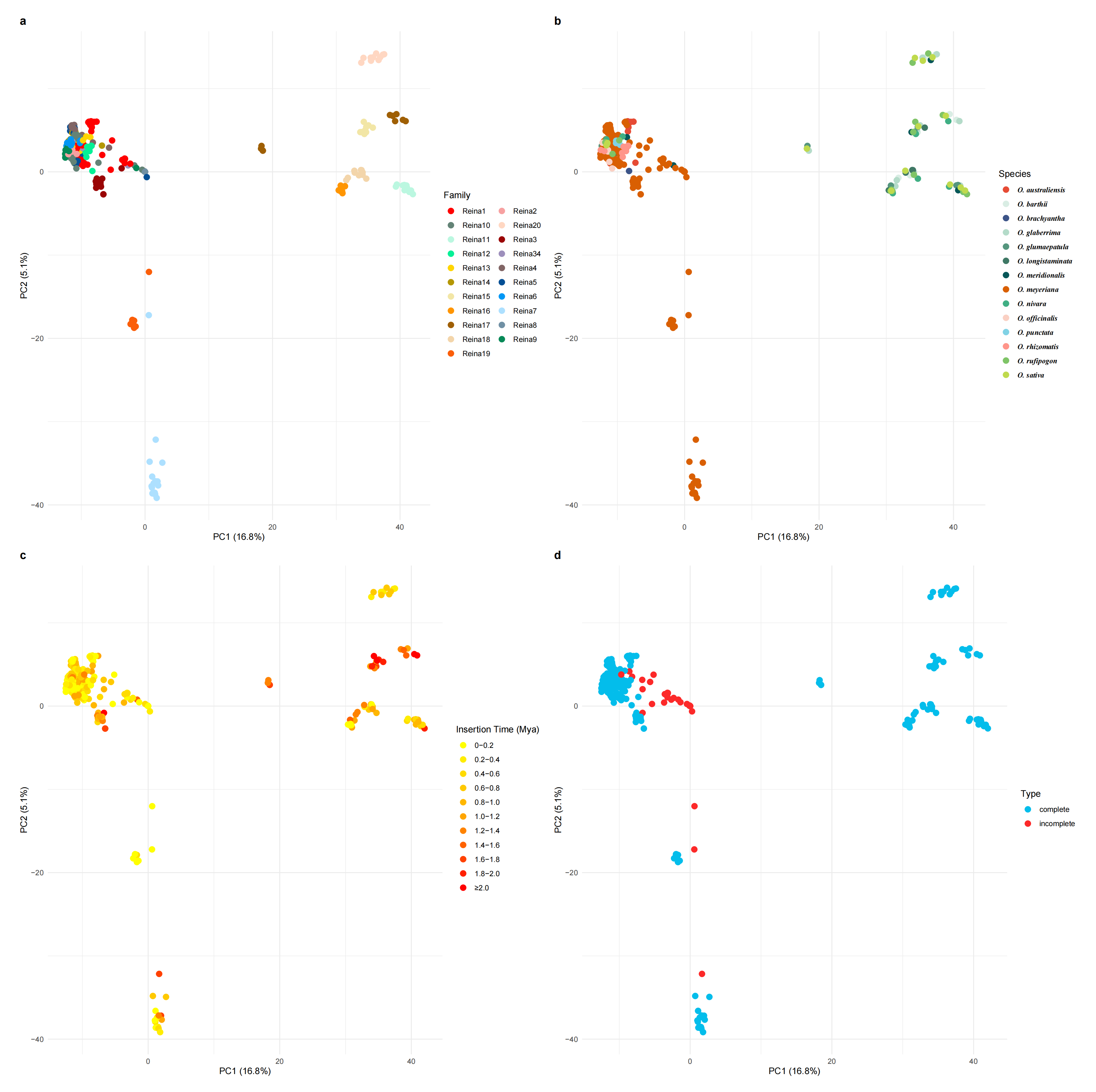

**Fig. S42. The dynamic evolution of the main LTR-RTs families in Reina lineage, in Oryza species.** **a** indicates the PCA analysis of main intact LTR-RTs families. **b** indicates the species including these intact LTR-RTs. **c** indicates the insertion time of these intact LTR-RTs. **d** indicates the autonomous intact LTR-RTs and non-autonomous intact LTR-RT.

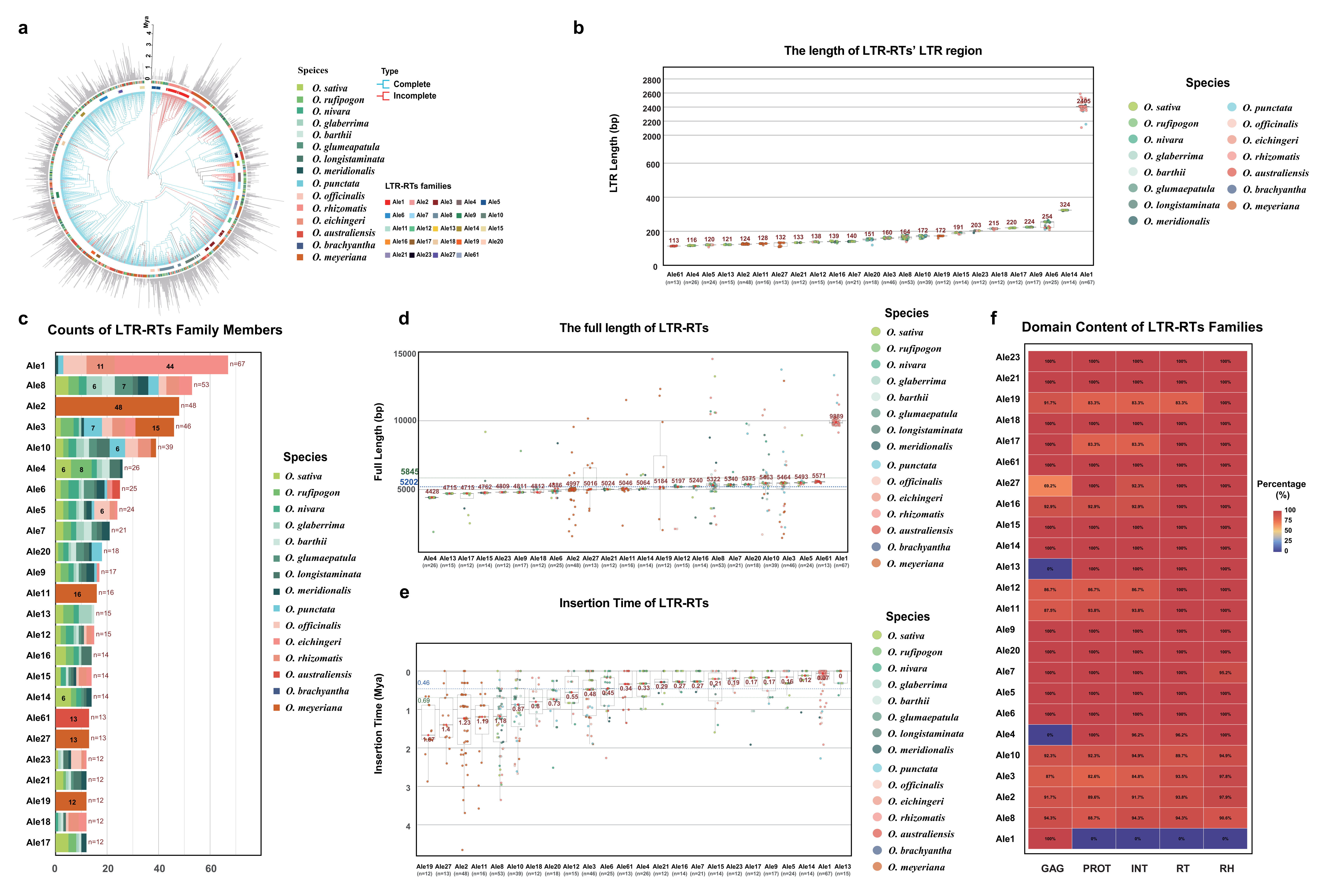

**Fig. S43. The phylogenetic tree of intact LTR-RTs and the information of main intact LTR-RTs families in Ale lineage.** **a** is a phylogenetic tree of LTR-RTs, constructed based on the 5’ LTR region. The outermost bar plot indicates the insertion time of LTR-RTs, the middle ring layer shows the species where LTR-RTs are located, and the innermost ring represents the major families to which LTR-RTs belong. The branch colors indicate whether the domains are complete, and also distinguish whether the LTR-RTs are autonomous or non-autonomous. **b** is a boxplot showing the length of the 5’ LTR region for the major LTR-RT families. **c** shows the number and distribution of members in the major LTR-RT families. **d** is a boxplot displaying the length of members in the major LTR-RT families. **e** is a boxplot illustrating the insertion time of members in the major LTR-RT families. The green dashed line represents the mean value, while the blue dashed line indicates the median value. **f** shows the domain content of the major LTR-RT families.

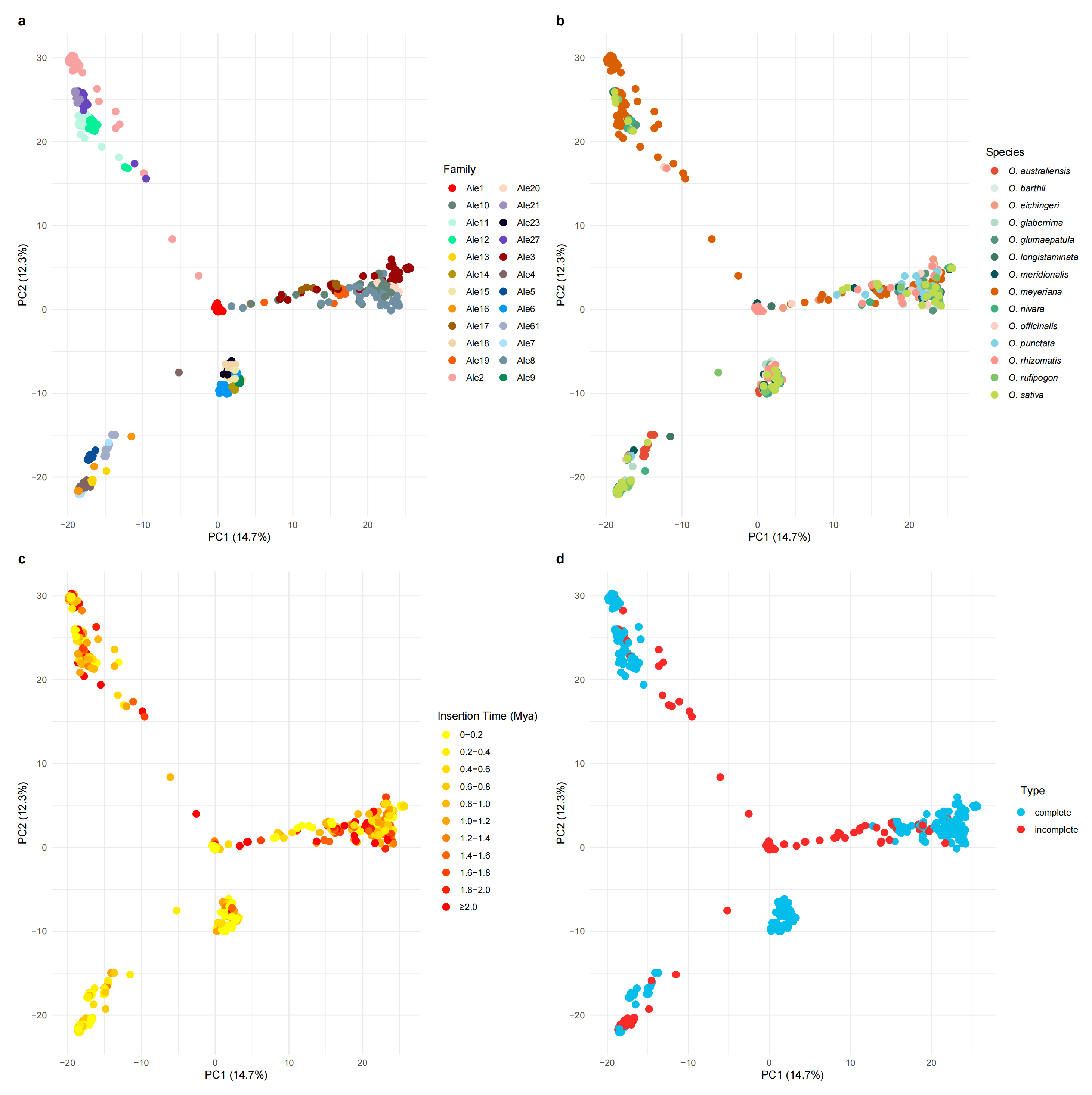

**Fig. S44. The dynamic evolution of the main LTR-RTs families in Ale lineage, in Oryza species.** **a** indicates the PCA analysis of main intact LTR-RTs families. **b** indicates the species including these intact LTR-RTs. **c** indicates the insertion time of these intact LTR-RTs. **d** indicates the autonomous intact LTR-RTs and non-autonomous intact LTR-RT.

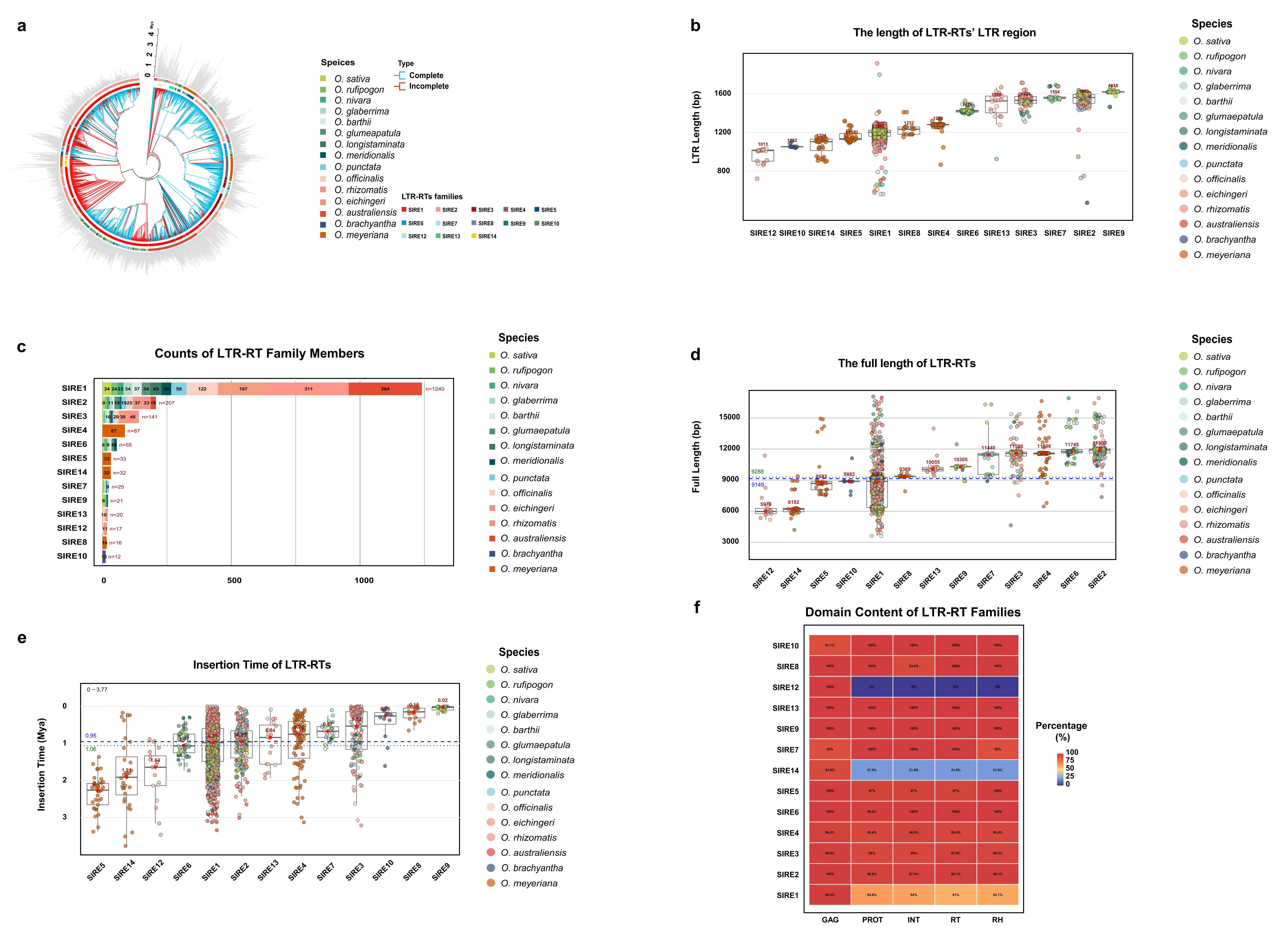

**Fig. S45. The phylogenetic tree of intact LTR-RTs and the information of main intact LTR-RTs families in SIRE lineage.** **a** is a phylogenetic tree of LTR-RTs, constructed based on the 5’ LTR region. The outermost bar plot indicates the insertion time of LTR-RTs, the middle ring layer shows the species where LTR-RTs are located, and the innermost ring represents the major families to which LTR-RTs belong. The branch colors indicate whether the domains are complete, and also distinguish whether the LTR-RTs are autonomous or non-autonomous. **b** is a boxplot showing the length of the 5’ LTR region for the major LTR-RT families. **c** shows the number and distribution of members in the major LTR-RT families. **d** is a boxplot displaying the length of members in the major LTR-RT families. **e** is a boxplot illustrating the insertion time of members in the major LTR-RT families. The green dashed line represents the mean value, while the blue dashed line indicates the median value. **f** shows the domain content of the major LTR-RT families.

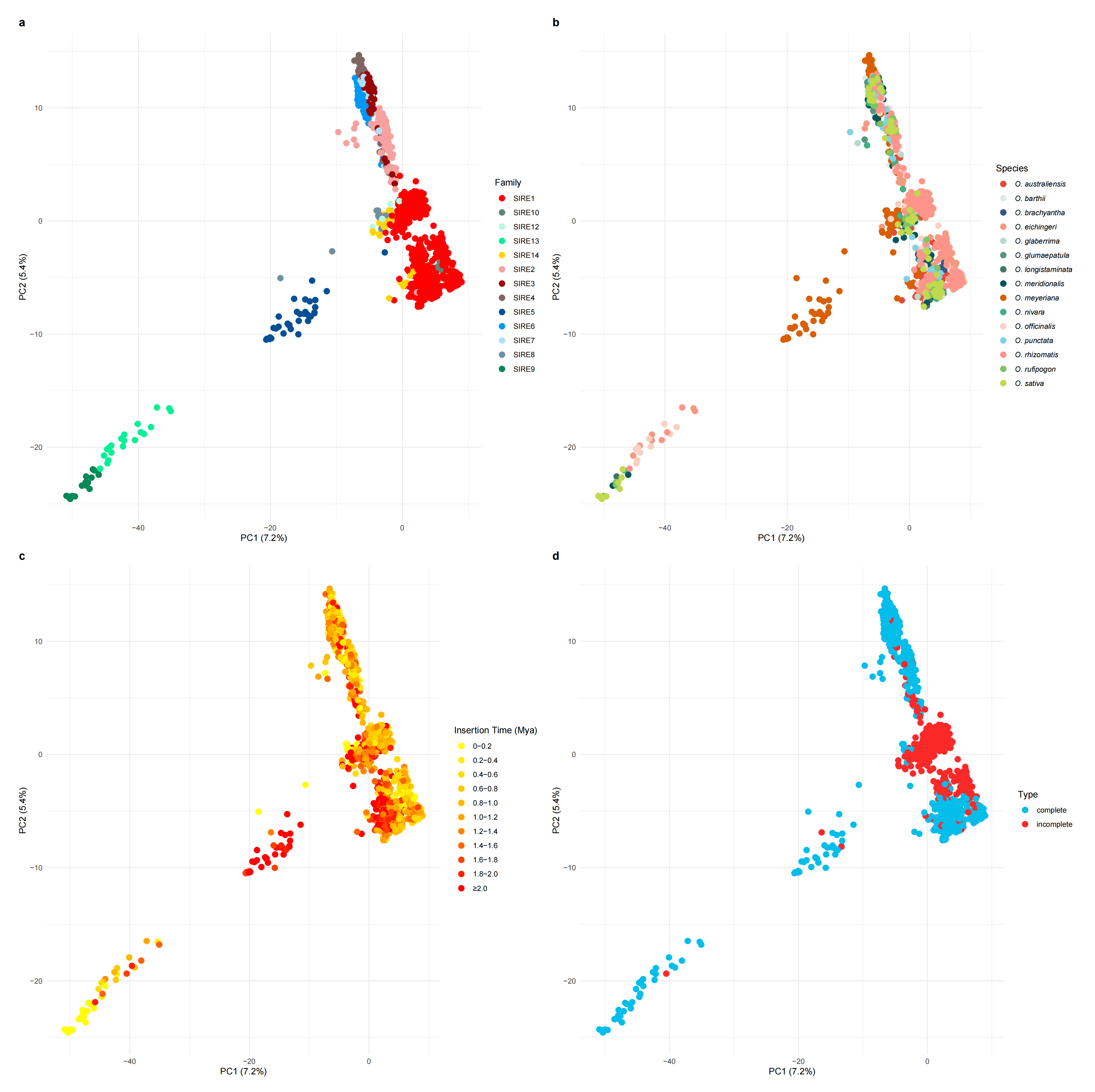

**Fig. S46. The dynamic evolution of the main LTR-RTs families in SIRE lineage, in Oryza species.** **a** indicates the PCA analysis of main intact LTR-RTs families. **b** indicates the species including these intact LTR-RTs. **c** indicates the insertion time of these intact LTR-RTs. **d** indicates the autonomous intact LTR-RTs and non-autonomous intact LTR-RTs.

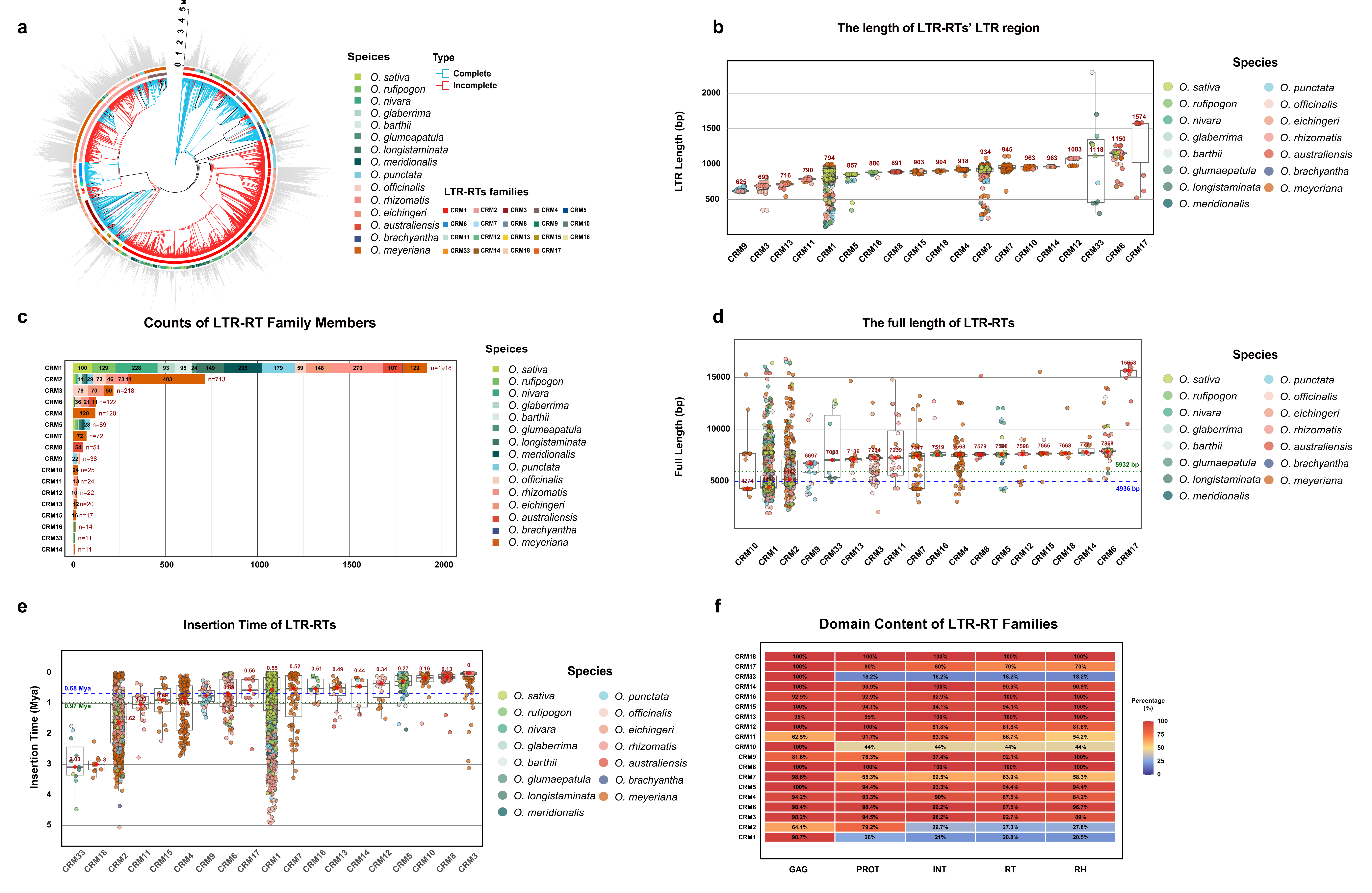

**Fig. S47. The phylogenetic tree of intact LTR-RTs and the information of main intact LTR-RTs families in CRM lineage.** **a** is a phylogenetic tree of LTR-RTs, constructed based on the 5’ LTR region. The outermost bar plot indicates the insertion time of LTR-RTs, the middle ring layer shows the species where LTR-RTs are located, and the innermost ring represents the major families to which LTR-RTs belong. The branch colors indicate whether the domains are complete, and also distinguish whether the LTR-RTs are autonomous or non-autonomous. **b** is a boxplot showing the length of the 5’ LTR region for the major LTR-RT families. **c** shows the number and distribution of members in the major LTR-RT families. **d** is a boxplot displaying the length of members in the major LTR-RT families. **e** is a boxplot illustrating the insertion time of members in the major LTR-RT families. The green dashed line represents the mean value, while the blue dashed line indicates the median value. **f** shows the domain content of the major LTR-RT families.

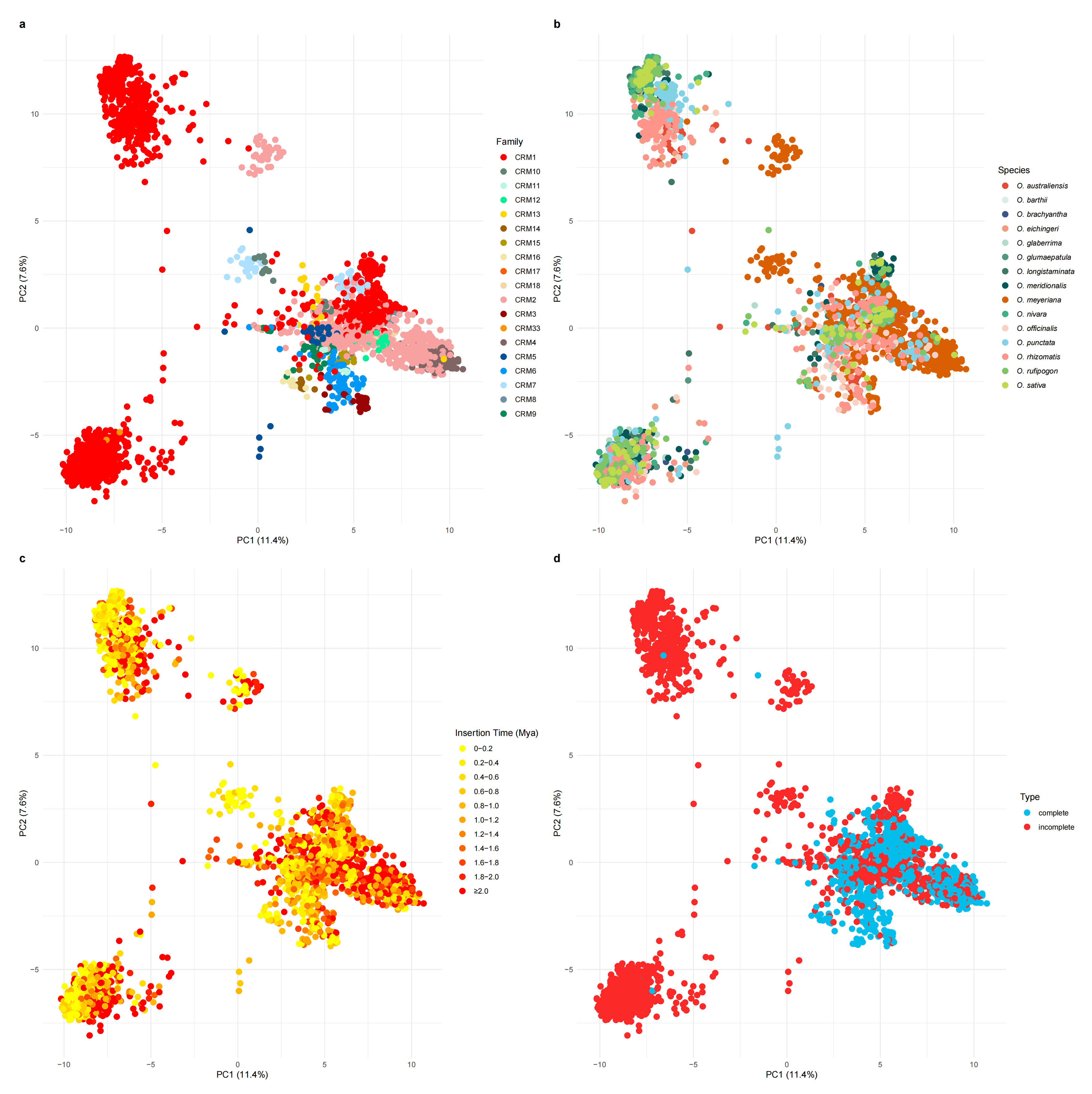

**Fig. S48. The dynamic evolution of the main LTR-RTs families in CRM lineage, in Oryza species.** **a** indicates the PCA analysis of main intact LTR-RTs families. **b** indicates the species including these intact LTR-RTs. **c** indicates the insertion time of these intact LTR-RTs. **d** indicates the autonomous intact LTR-RTs and non-autonomous intact LTR-RTs.

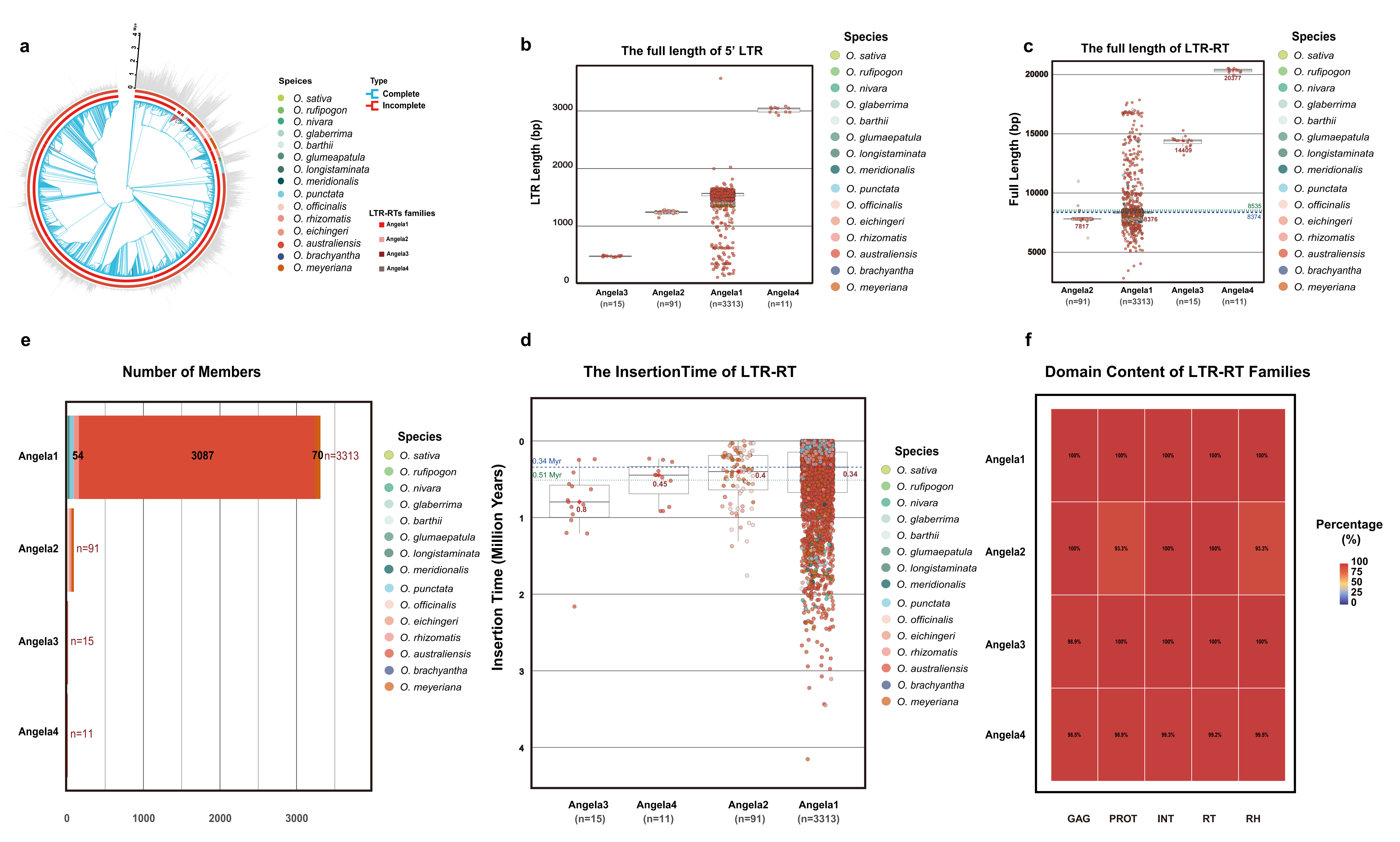

**Fig. S49. The phylogenetic tree of intact LTR-RTs and the information of main intact LTR-RTs families in Angela lineage.** **a** is a phylogenetic tree of LTR-RTs, constructed based on the 5’ LTR region. The outermost bar plot indicates the insertion time of LTR-RTs, the middle ring layer shows the species where LTR-RTs are located, and the innermost ring represents the major families to which LTR-RTs belong. The branch colors indicate whether the domains are complete, and also distinguish whether the LTR-RTs are autonomous or non-autonomous. **b** is a boxplot showing the length of the 5’ LTR region for the major LTR-RT families. **c** shows the number and distribution of members in the major LTR-RT families. **d** is a boxplot displaying the length of members in the major LTR-RT families. **e** is a boxplot illustrating the insertion time of members in the major LTR-RT families. The green dashed line represents the mean value, while the blue dashed line indicates the median value. **f** shows the domain content of the major LTR-RT families.

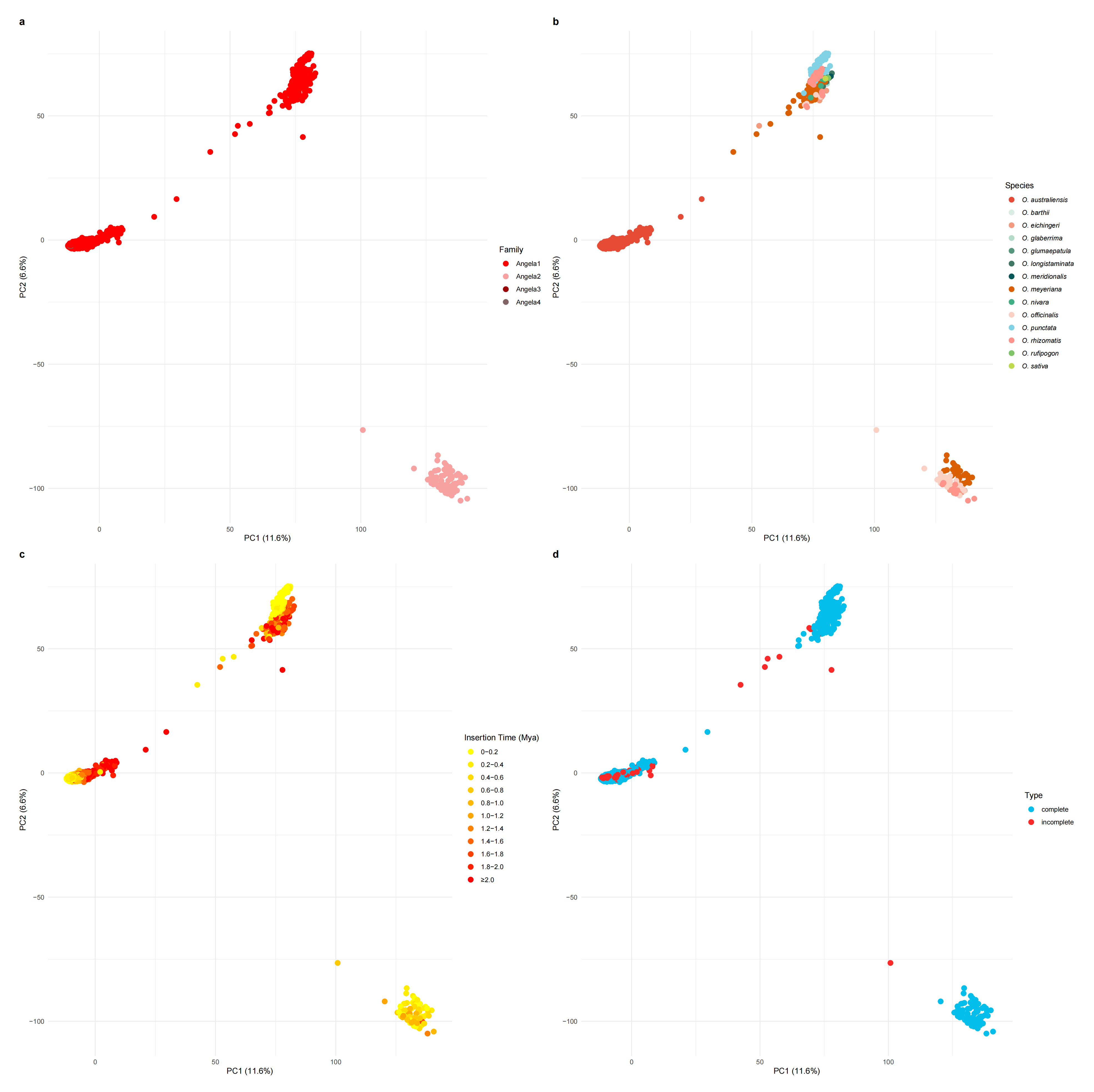

**Fig. S50. The dynamic evolution of the main LTR-RTs families in Angela lineage, in Oryza species.** **a** indicates the PCA analysis of main intact LTR-RTs families. **b** indicates the species including these intact LTR-RTs. **c** indicates the insertion time of these intact LTR-RTs. **d**, indicates the autonomous intact LTR-RTs and non-autonomous intact LTR-RTs.

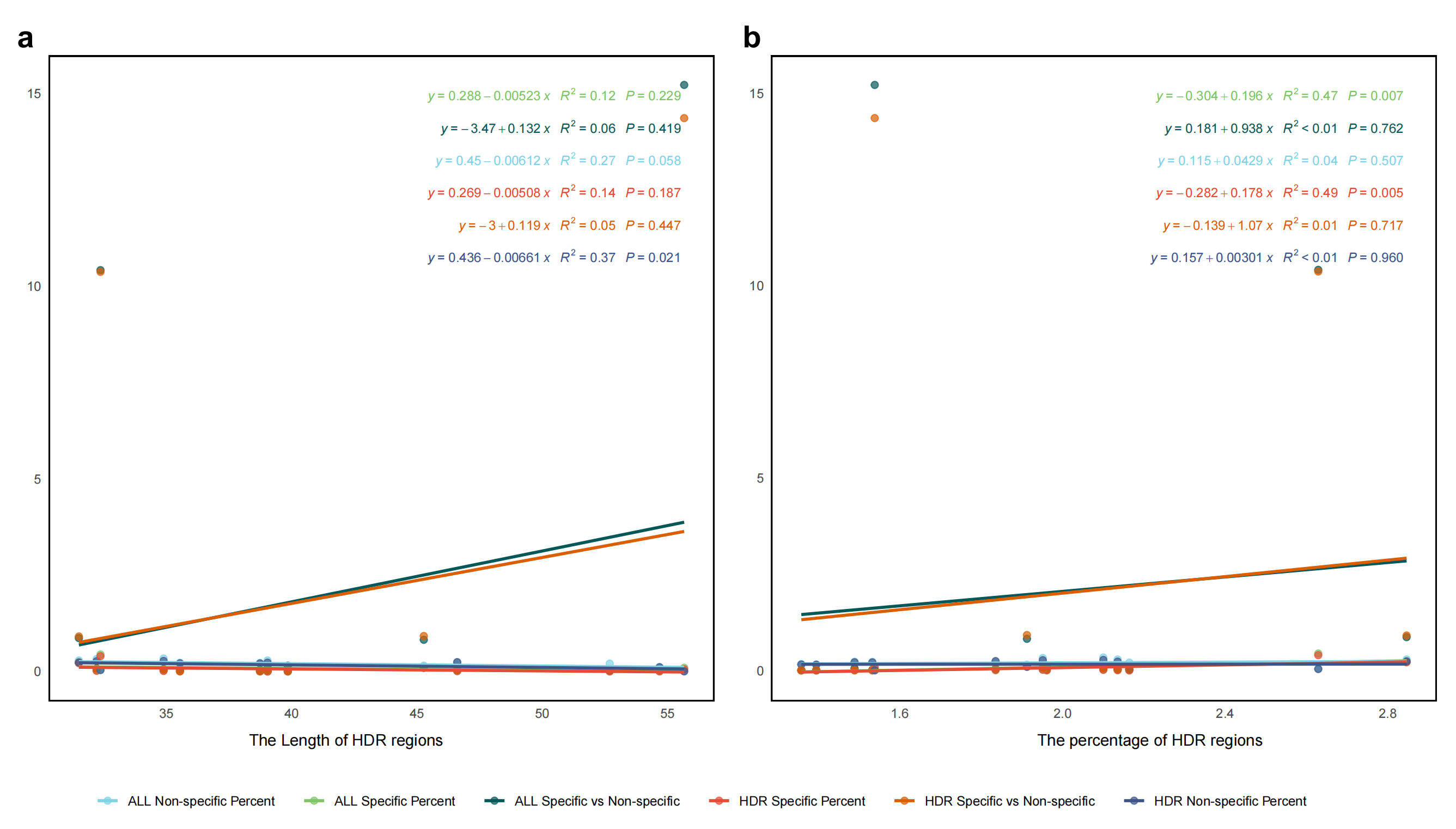

**Fig. S51.** **The pearson correlation analysis between the content of different LTR-RTs families in all region.** **a,** the length of HDRs and **b,** percentage of HDRs.

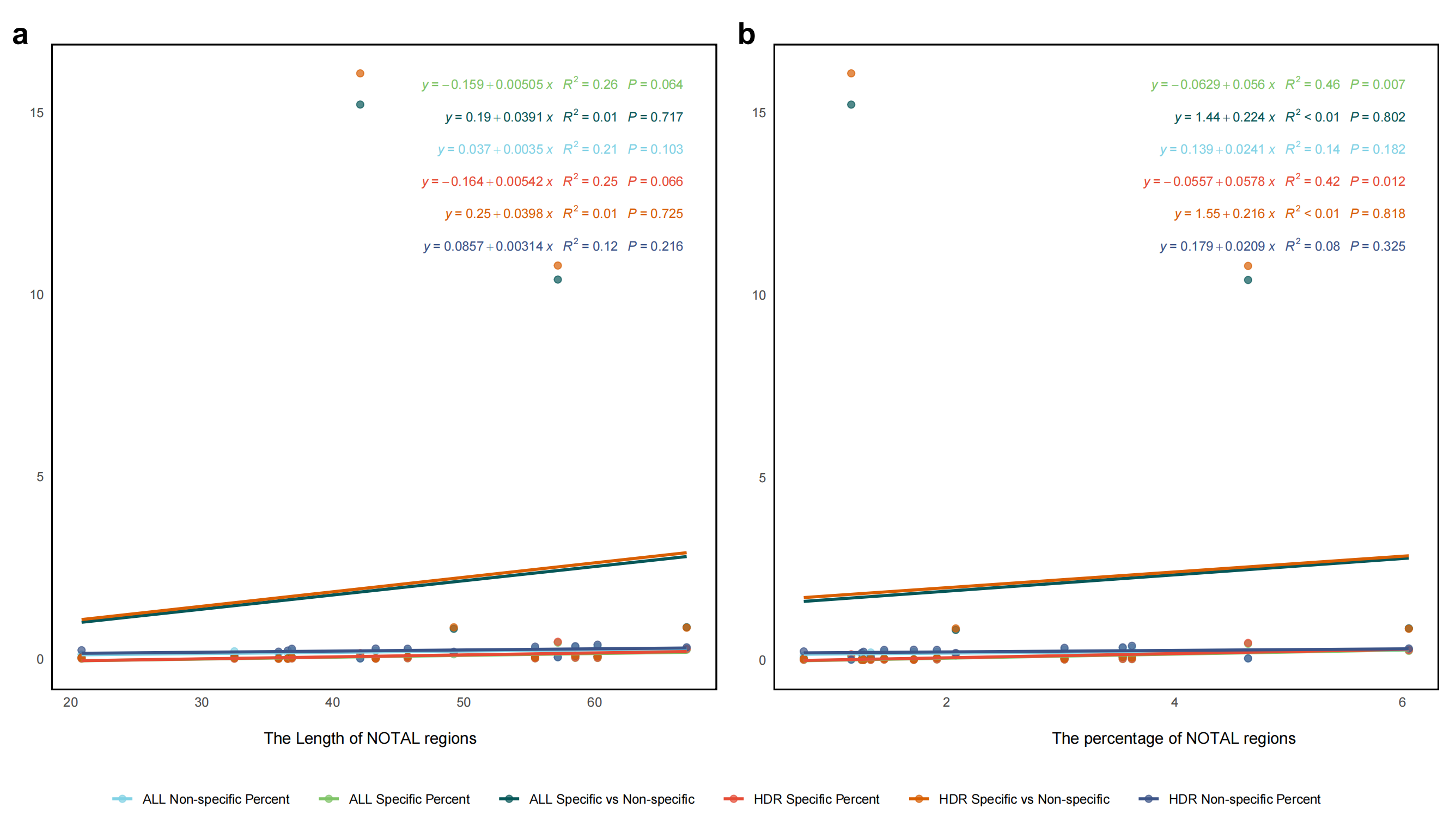

**Fig. S52.** **The pearson correlation analysis between the content of different LTR-RTs families in all region.** **a,** the length of NOTALs and **b,** percentage of NOTALs.

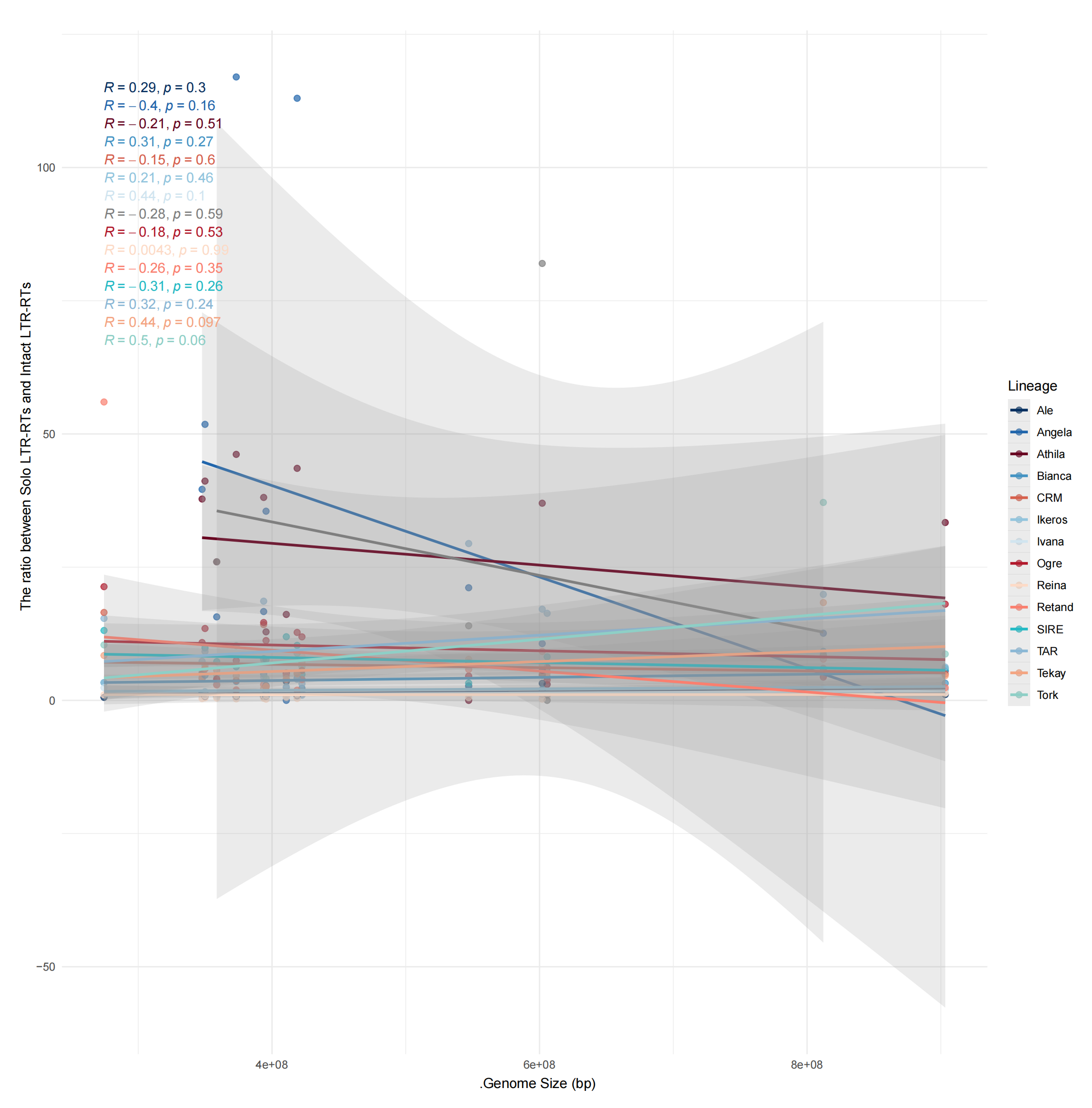

**Fig. S53**. **The pearson correlation analysis between the deletion rate of LTR-RTs from different lineages and genome size in different species.**

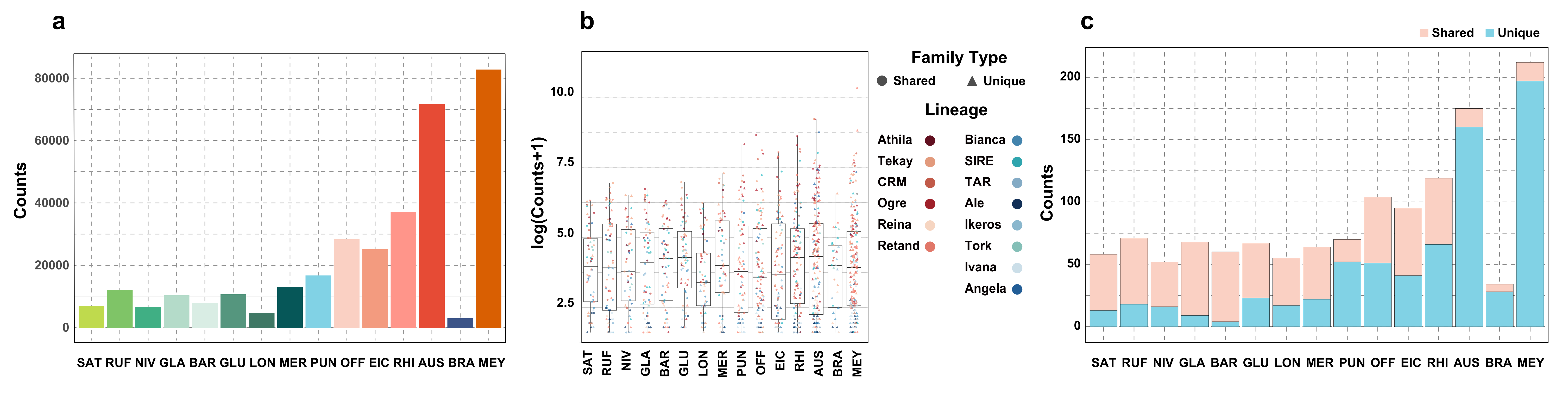

**Fig. S54.** **The variation of the member counts and family counts of high removal rate of LTR-RTs families in different species.** **a** is the member counts of high removal rate of LTR-RTs families in different species. **b** is the member counts of high removal rate of LTR-RTs families from different lineages in different species. **c** is the family counts of high removal rate of LTR-RTs families belonging to species-specific LTR-RTs families or non-species-specific LTR-RTs families in different species. Shared: non-species-specific LTR-RTs families. Unique: species-specific LTR-RTs families.

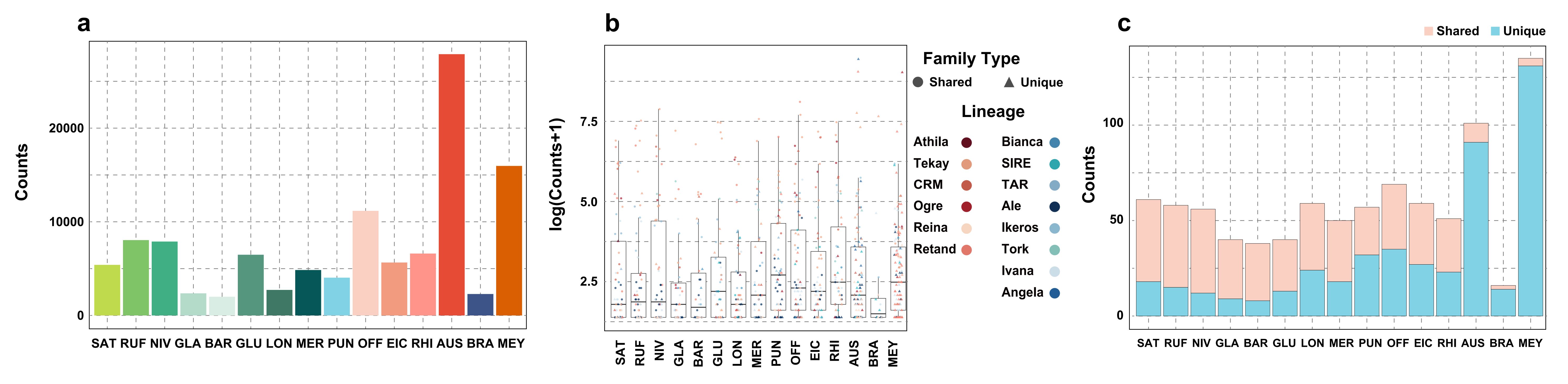

**Fig. S55.** **The variation of the member counts and family counts of medium removal rate of LTR-RTs families in different species.** **a** is the member counts of high removal rate of LTR-RTs families in different species. **b** is the member counts of high removal rate of LTR-RTs families from different lineages in different species. **c** is the family counts of high removal rate of LTR-RTs families belonging to species-specific LTR-RTs families or non-species-specific LTR-RTs families in different species. Shared: non-species-specific LTR-RTs families. Unique: species-specific LTR-RTs families.

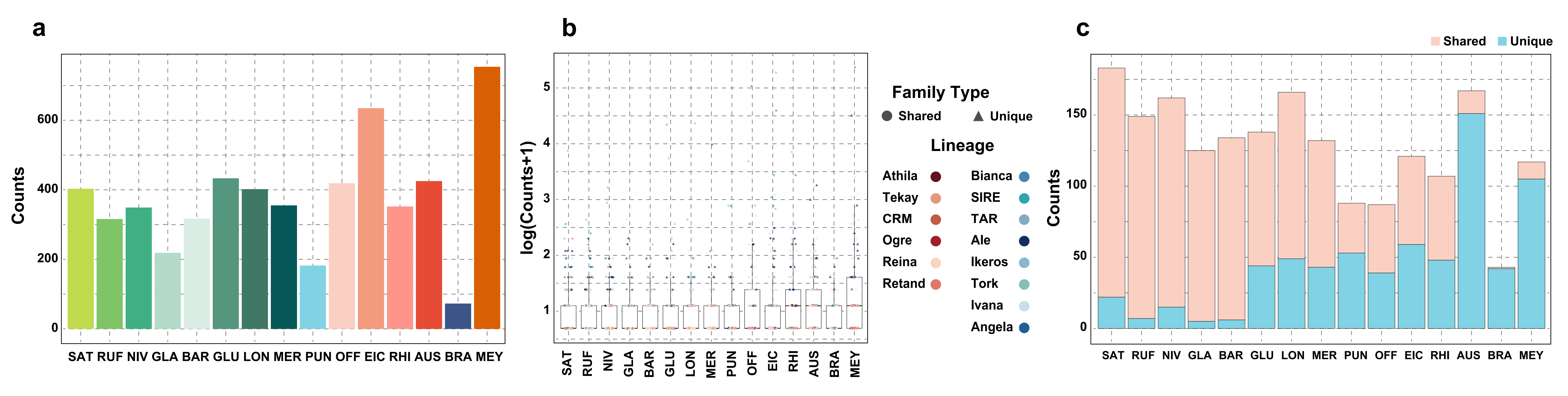

**Fig. S55.** **The variation of the member counts and family counts of low removal rate of LTR-RTs families in different species.** **a** is the member counts of high removal rate of LTR-RTs families in different species. **b** is the member counts of high removal rate of LTR-RTs families from different lineages in different species. **c** is the family counts of high removal rate of LTR-RTs families belonging to species-specific LTR-RTs families or non-species-specific LTR-RTs families in different species. Shared: non-species-specific LTR-RTs families. Unique: species-specific LTR-RTs families.
